# IsoGraph resolves coordinated transcript choice and gene abundance in the aging human brain

**DOI:** 10.64898/2026.10.04.756439

**Authors:** Alexis Bennett, Elisa Kain Johnson, Kynon J M Benjamin

## Abstract

Relative transcript usage can change without a corresponding change in total gene expression, but resolving coordinated transcript choice across genes while separating it from abundance remains challenging. We present IsoGraph, a gene-centered framework that represents isoform usage and abundance as paired network channels, identifies gene modules, and quantifies their channel-specific contributions. In synthetic benchmarks, IsoGraph preferentially recovered network structure when coordinated transcript usage carried the planted signal, whereas abundance-based representations were more informative in abundance-dominated and RNA-degradation settings. Applied to BrainSEQ and GTEx adult brain transcriptomes, IsoGraph identified age-associated transcript-usage signals that persisted after conditioning on gene abundance and, in a region-dependent manner, after adjustment for estimated cellular composition. Modules were stable across donor subsets, and frozen module axes retained age associations across cohorts even when exact gene partitions differed. IsoGraph module context captured predictive information about held-out differential transcript usage beyond gene-wise evidence and conventional co-expression. Convergent evidence from structural annotation, short-read junction usage, independent long-read sequencing, and within-donor allelic analyses supported constituent transcript events. Disease colocalization further prioritized candidate isoform events at neurological and psychiatric disease loci, including an alternative terminal-exon event at *PRDM2* associated with amyotrophic lateral sclerosis. Together, these results establish coordinated transcript usage as a reproducible component of transcriptomic organization and provide a framework for resolving its contribution alongside gene abundance and for tracing network-level signals to candidate transcript events.

## Introduction

Variation in gene regulation involves both total gene abundance and the relative use of alternative transcript isoforms. Summing transcripts into a gene-level measurement can obscure changes in transcript choice that alter coding sequences, untranslated regions (UTRs), or other regulatory features without a corresponding change in total expression. Transcript quantification enables these dimensions to be studied in human cohorts^1,2^, and brain studies have identified extensive isoform regulation across development, anatomical regions, and disease^3–7^. In aging brain tissue, separating variation in transcript choice from variation in gene abundance is further complicated by cellular composition, while differences in cohort composition and RNA-sequencing workflows challenge the reproducibility of transcript-usage signals across studies. These challenges raise a broader question: whether age-associated changes in transcript choice form coordinated programs across genes that can be distinguished from abundance effects and reproduced across human brain cohorts.

Differential transcript usage (DTU) methods characterize changes within individual genes, and structural annotation can identify their predicted consequences^8,9^. Existing network approaches already extend isoform analysis across genes. Saha *et al*. developed Transcriptome-Wide Networks that represent total expression and relative isoform abundance as distinct nodes within a common sparse network, allowing relationships among transcriptional and splicing features to be modeled directly^10^. A human neocortex study subsequently constructed separate weighted gene co-expression network analysis (WGCNA)^11^ networks of gene expression, isoform expression, and isoform usage^12^, while the acorde framework detects co-differential isoform usage across cell types, including relationships not detected from gene-level expression^13^. More recent work has also constructed isoform co-expression networks in adult human cortex^14^. Together, these studies establish that transcript abundance and isoform usage contain complementary network information, but existing approaches do not directly resolve their respective contributions to shared gene-level network structure. We therefore asked how transcript choice and total abundance contribute to the same gene modules, which transcripts underlie usage-driven structure, and whether these patterns transfer across cohorts.

Here, we introduce IsoGraph, an open-source Python package that represents within-gene transcript composition and total gene abundance as separate network channels. It identifies gene modules, assigns channel-specific roles to their members, and relates usage-associated network coordinates to contributing transcripts and annotated structures. We evaluated the framework using synthetic benchmarks and human brain RNA-sequencing data from BrainSEQ^15,16^ and the Genotype-Tissue Expression (GTEx) project^17^. We asked whether transcript usage captures age-associated variation beyond gene abundance and estimated cellular composition; whether gene modules and their age associations remain stable across donor subsets and transferable across cohorts; and whether module context improves the prioritization of transcript-usage events beyond gene-wise evidence. We then examined molecular support for constituent transcript events using long-read sequencing^18^, short-read junction usage, and allelic junction analyses^19^, and assessed their disease relevance through colocalization^20,21^ and complementary credible-set posterior _overlap_^22^. Together, these analyses evaluate whether coordinated transcript usage constitutes a reproducible dimension of gene-network organization that can be separated from abundance and traced from module-level structure to specific transcript events and disease-associated loci.

## Results

### IsoGraph resolves coordinated isoform usage alongside gene abundance

IsoGraph represents each gene through a within-gene isoform-usage coordinate and a separate abundance coordinate. These paired features are used to infer a gene network and identify modules, after which genes are assigned channel-specific roles according to the contributions of switching and abundance to module membership (*Methods*; **Fig. 1a,b**). Transcript loadings and structural annotations connect the gene-level switch coordinate to candidate transcript pairs (**Fig. 1a**). The switch coordinate captures continuous variation in relative transcript usage, including changes that do not involve exchange of the dominant isoform (**Fig. 1c**). Usage and abundance are modeled as distinct channels while retaining their empirical dependence.

**Fig. 1.**
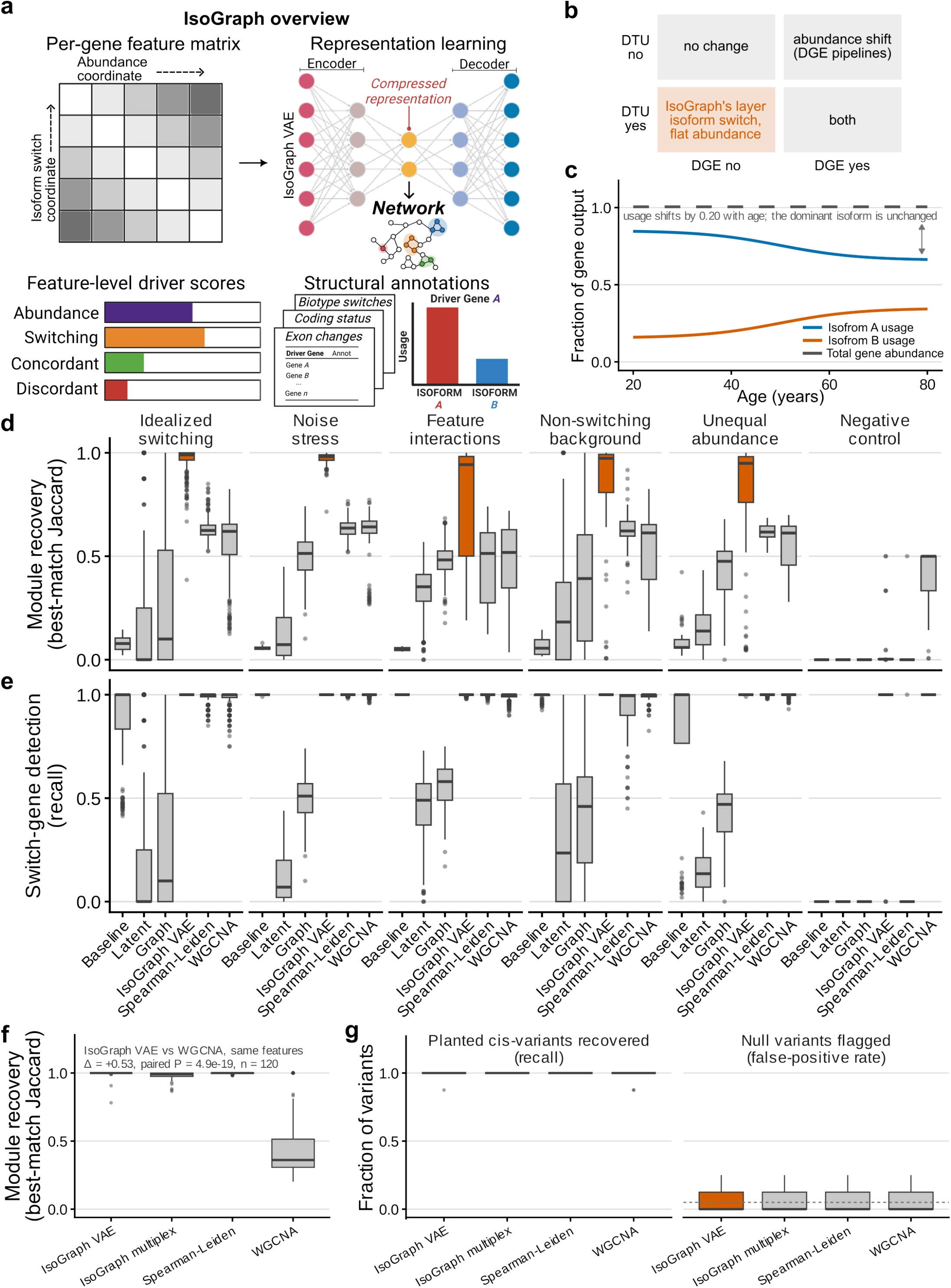
IsoGraph jointly represents isoform usage and gene abundance and recovers planted switching structure in synthetic data. **a**. Overview of the IsoGraph framework. Transcript-level measurements are decomposed into a within-gene isoform-usage coordinate and a separate gene-abundance coordinate. These features are jointly encoded into a compressed representation by a variational autoencoder (VAE) and used to construct a gene network and identify modules. Channel-specific contributions distinguish usage- and abundance-associated gene signals, while transcript loadings and structural annotations link gene-level usage coordinates to candidate transcript pairs and associated transcript features. Created in *BioRender*.*com*. **b**. Conceptual relationship between isoform-usage and gene-abundance changes. Genes may exhibit neither signal, predominantly abundance change, predominantly isoform-usage change, or both. IsoGraph retains usage and abundance as distinct channels within a shared representation. **c**. Schematic illustrating continuous variation of isoform usage at constant gene abundance. Relative transcript usage can change continuously without requiring a binary exchange of the dominant isoform. **d–e**. Performance across synthetic scenarios comprising idealized switching, noise stress, feature interactions, non-switching background, unequal abundance, and a negative control (*n* = 40–360 simulation runs per scenario and method; Table S1). **d**. Recovery of planted gene modules, quantified by best-match Jaccard similarity (1 indicates perfect alignment with the planted module). **e**. Recall of planted switch genes. Orange denotes the switch-only IsoGraph VAE. **f–g**. Synthetic-genetics benchmark using 120 matched simulated datasets with identical input features. **f**. Recovery of planted gene modules by IsoGraph and weighted gene co-expression network analysis (WGCNA), quantified by best-match Jaccard similarity. **g**. Detection of planted genetic associations, shown as recall of planted *cis*-variants and false-positive rate for unlinked null variants; the dashed horizontal line indicates a false-positive rate of 0.05. Boxplots show the median and interquartile range; whiskers extend to 1.5 × the interquartile range, and points denote individual simulation replicates.

In simulations with known generating transcripts, transcript loading ranked the planted switch transcript first in 99.9% of switching genes (mean top-1 accuracy across 40 multi-isoform simulations; chance 0.25), and switch strength distinguished switching from non-switching genes with a mean area under the receiver operating characteristic curve of 1.00 under idealized switching (**Fig. S1**). In mixed switch–abundance simulations, the multiplex representation retained abundance-shift detection as abundance mixing increased, while its inferred role composition shifted toward abundance-associated assignments (**Fig. S2**). These results support the intended interpretation of the paired coordinates and their channel-specific roles.

Synthetic benchmarks evaluated module recovery across switching, abundance, degradation, and scale settings (**Fig. 1d,e** and **Fig. S3**). We scored recovery primarily by best-match Jaccard similarity, where 1 indicates perfect alignment with the planted module (**Table S1**). IsoGraph outperformed WGCNA^11^ in the idealized-switching scenarios (mean difference 0.396), and this advantage persisted across stress tests evaluating count dispersion, feature interactions, and unequal isoform abundance (**Fig. S4** and **Fig. S5**). Residualization also improved robustness to simulated cell-composition, batch, and library-depth effects (**Fig. S6**). In a separate synthetic-genetics arm, both tools detected the planted genetic signal, but IsoGraph recovered module organization more accurately (mean recovery 0.997 versus 0.466 for WGCNA; one-sided paired Wilcoxon *P* = 4.9 × 10^−19^; **Fig. 1f,g**). Complementary metrics (adjusted Rand index, homogeneity, completeness) supported these recovery patterns. Pure-noise negative controls showed little recovery of the planted partition, supporting specificity of the module-recovery metric (**Fig. S7** and **Fig. S8**).

IsoGraph was less effective when degradation or abundance, rather than coordinated transcript usage, dominated the planted structure (**Fig. S6**). Abundance-dominated and degradation-coupled scenarios favored WGCNA over IsoGraph (mean differences −0.344 and −0.559, respectively). However, when degradation affected switching but abundance retained the planted structure, incorporating the abundance channel substantially rescued module recovery from approximately 0.17–0.21 to 0.67–0.69 across degradation levels (**Fig. S9**). Finally, at transcriptome scale, the recovery difference between methods was approximately zero, and the implementation showed efficient runtime and memory scaling (**Fig. S10**; **Table S2**). Overall, these benchmarks show that IsoGraph preferentially recovers network structure driven by coordinated transcript usage, while the paired abundance channel preserves information when network organization shifts toward gene-level expression.

### Phenotype-associated isoform usage captures signals beyond gene abundance across human brain regions

We next asked whether phenotype-associated transcript-usage signals in human brain were distinguishable from changes in total gene abundance. We applied IsoGraph across 16 aging analyses spanning three BrainSEQ regions (caudate, hippocampus, and dorsolateral prefrontal cortex [DLPFC])^15,16^ and 13 GTEx brain regions^17^, alongside a schizophrenia comparison in BrainSEQ caudate^16^ (**Fig. 2a**; **Table 1**; **Data S1**). BrainSEQ and GTEx were processed independently, and donors were partly shared across regions within each cohort (**Fig. S11**).

**Table 1.** BrainSEQ sample sets by analytical purpose. Sample counts are listed in the order caudate, hippocampus, and dorsolateral prefrontal cortex unless a single region is stated. The ASE resource is the set of samples with allele-aware alignments available for junction recounting; the number of donors fitted in each allelic test was determined separately for each transcript pair by informative fragment counts and lead-heterozygous donor coverage. ASE, allele-specific expression; BAM, binary alignment map; VCF, variant call format; QTL, quantitative trait locus; GWAS, genome-wide association study.

| Analysis or resource | Eligibility | Samples by region | Use |
| --- | --- | --- | --- |
| Aging discovery | Neurotypical controls, age $\geq 18$ years | 238, 238, 222 | Network discovery and aging analyses |
| Schizophrenia discovery | Controls and schizophrenia cases, age $\geq 18$ years | 390 in caudate (238 controls, 152 cases) | Diagnosis analysis |
| Switch-QTL and abundance-QTL mapping | All available diagnoses, age $> 13$ years, genotype matched | 420, 362, 360 | QTL mapping; a European-ancestry subset was used for GWAS colocalization |
| Available junction-specific ASE resource | ASE-grade BAM, index, and phASER VCF available | 486, 451, 498 | Starting resource for junction recounting, not the number fitted in each allelic test |

**Fig. 2.**
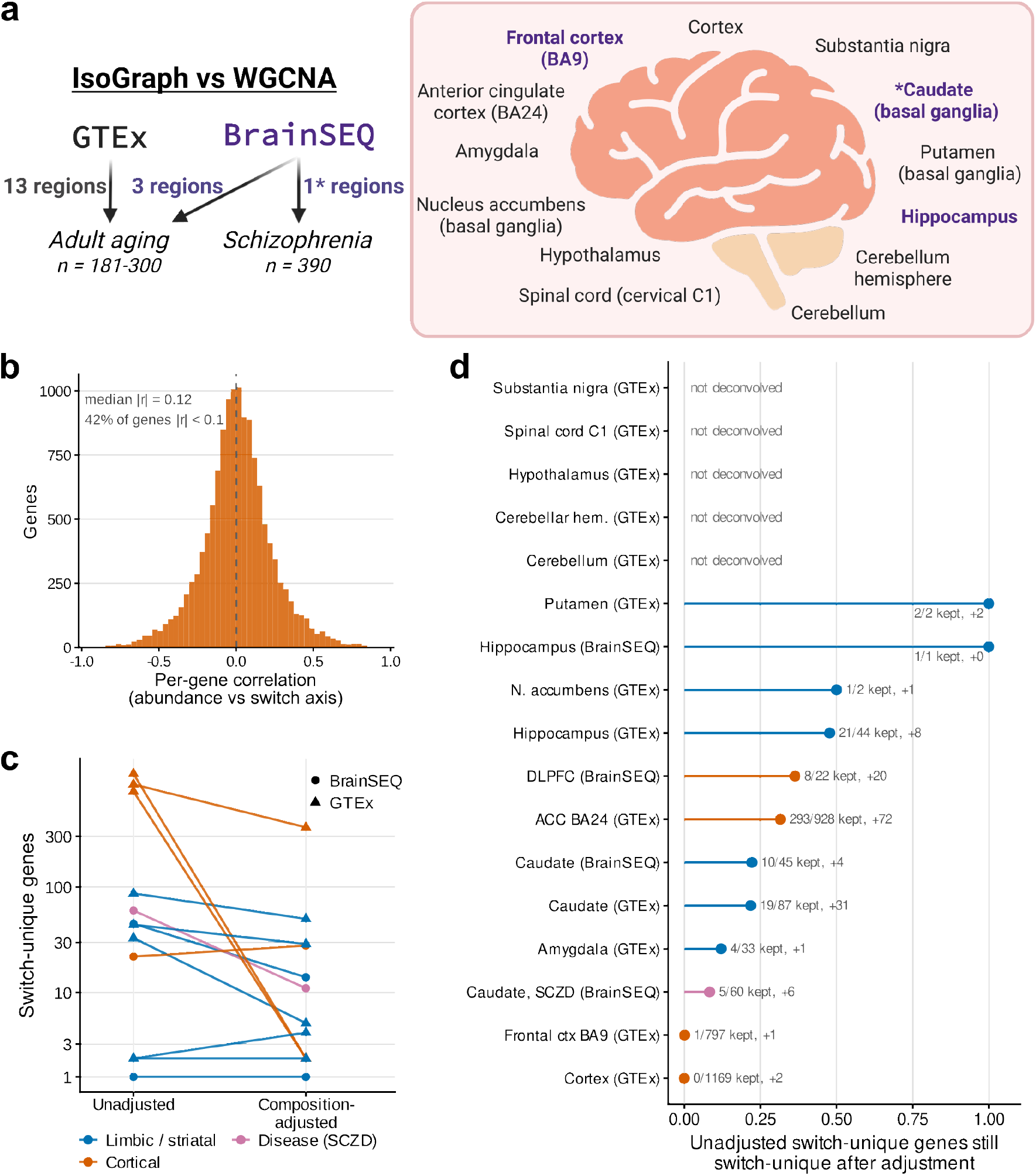
IsoGraph identifies isoform-usage structure distinct from gene abundance across adult human brain datasets. **a**. Study design. IsoGraph and weighted gene co-expression network analysis (WGCNA) were applied to adult-aging RNA-seq data from 13 Genotype-Tissue Expression (GTEx) brain regions and three BrainSEQ regions (caudate, hippocampus, and dorsolateral prefrontal cortex [DLPFC]), with a separate BrainSEQ caudate schizophrenia analysis. Adult-aging analyses included 181–300 donors per region; the schizophrenia analysis included 390 samples. BrainSEQ regions are highlighted in purple. Created in *BioRender*.*com*. **b**. Distribution of per-gene correlations between gene-abundance and isoform-switch coordinates for the 13,222 genes tested in the BrainSEQ caudate schizophrenia analysis. **c**. Switch-unique genes (conditional switch association at false discovery rate [FDR] < 0.10 without a conditional abundance association) before and after adjustment for estimated cellular composition, for the 12 analyses with an anatomically matched single-nucleus reference. Connected points denote the same analysis; circles indicate BrainSEQ and triangles GTEx. The *y* axis is logarithmically scaled. **d**. Fraction of unadjusted switch-unique genes that remain switch-unique after adjustment; labels give the number retained and the number newly detected after adjustment. Five GTEx regions lacking a matched reference are labeled as not deconvolved. Colors denote limbic/striatal regions (blue), cortical regions (orange), and the BrainSEQ schizophrenia analysis (pink).

Switch and abundance coordinates captured largely distinct aspects of within-gene variation (**Fig. 2b**). In the BrainSEQ caudate, the median absolute switch–abundance correlation was 0.12 across 13,222 genes, with 42% of genes showing weak correlation (|*r*| < 0.1); a similar separation was observed across all 16 aging analyses (median |*r*| = 0.11–0.24; **Fig. S12a**; **Table S3**). To isolate isoform-usage signals independent of total expression, we fit paired conditional models and defined “switch-unique” genes as those with a conditional switch association at false discovery rate (FDR) < 0.10 without a corresponding conditional abundance signal. For example, *MEGF6* showed schizophrenia-associated isoform usage in caudate without a detectable conditional gene-abundance association (**Fig. S12b**). Because sample size, variance, and feature filtering differed across analyses (**Fig. 2c**; **Fig. S12c**), switch-unique counts were interpreted within rather than quantitatively between regions.

We next tested whether switch-unique associations persisted after adjustment for estimated cellular composition in the 12 analyses with an anatomically matched single-nucleus reference^23^ (11 aging analyses and the schizophrenia analysis; **Fig. 2c,d**; **Data S2**). The effect of adjustment was strongly region dependent: substantial subsets of switch-unique associations persisted in anterior cingulate cortex and several BrainSEQ regions, whereas signals were markedly attenuated in GTEx frontal cortex and cortex. Adjustment changed the identity, not only the number, of detected genes. In BrainSEQ DLPFC, for example, 28 genes were detected after adjustment, but only 8 overlapped the 22 identified before adjustment. These qualitative patterns were stable across FDR thresholds of 0.05–0.20 (**Fig. S13**).

Thus, age-associated isoform usage showed heterogeneous relationships with estimated cellular composition across brain regions. In several regions, a subset of associations remained detectable after conditioning on both gene abundance and estimated cell proportions, whereas in others the signal was largely attenuated. Persistent associations therefore represent tissue-level signals robust to the estimated cell-composition covariates; bulk data do not resolve whether these effects are cell intrinsic. Conversely, attenuation after adjustment may reflect either confounding or biologically meaningful age-related changes in cellular composition (**Fig. S14a–c**). Consistent with the idea that IsoGraph modules were not simply proxies for cell-type composition, only 12 of 2,061 module-by-cell-type tests showed marker enrichment (FDR < 0.05), and none showed depletion (**Fig. S14d**), indicating that the modules were not generally enriched for the tested cell-type marker sets.

### IsoGraph modules are stable across donor subsets and transfer aging-associated axes across cohorts

We next evaluated the reproducibility of IsoGraph modules across donors and cohorts using three complementary criteria: stability of module structure across donor subsets, transfer of frozen module axes across cohorts, and independent recovery of similar gene partitions across cohorts (**Fig. 3a**; **Data S3**). These analyses complemented phenotype-association comparisons with WGCNA, for which IsoGraph did not show a higher overall fraction of associated modules (**Fig. S15**; **Table S4** and **Data S4**).

**Fig. 3.**
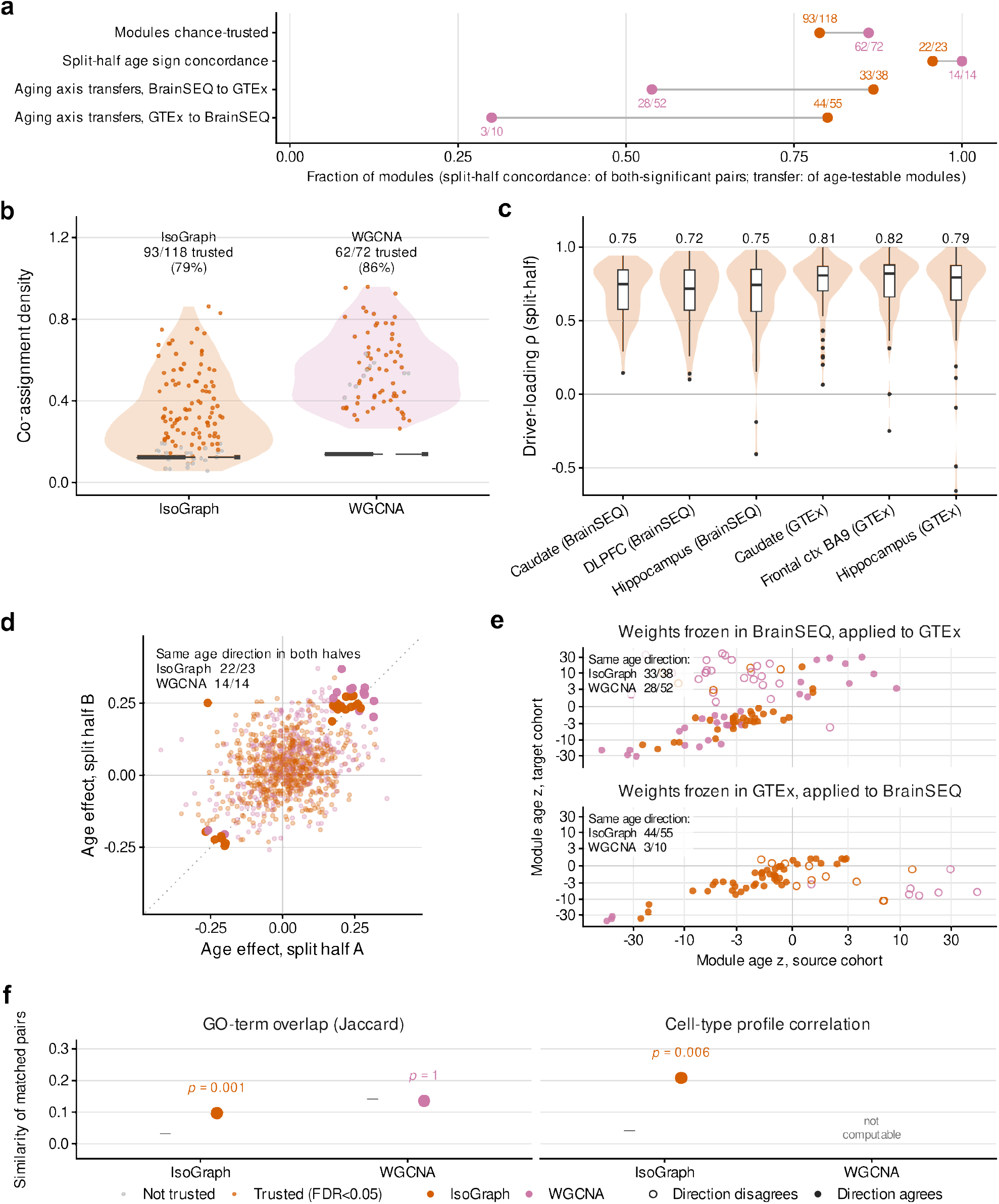
IsoGraph modules are reproducible across donor subsets and transfer age-associated structure across cohorts. **a**. Summary of four reproducibility measures for IsoGraph (orange) and for the matched weighted gene co-expression network analysis (WGCNA; purple) analyses; these method colors apply throughout. Numerators and denominators are shown for each measure because the eligible module sets differ across analyses. **b**. Split-half co-assignment density for every module of both methods across the six anatomically paired cohort-region analyses. Points are individual modules, colored orange when their density exceeded a size-matched permutation null (false discovery rate [FDR] < 0.05) and gray otherwise; the dashed line marks the median null. **c**. Spearman correlation of shared-gene switch-driver loadings between matched split halves, by analysis; values above each violin are regional medians. Classical WGCNA has no transcript-level drivers and is not shown. **d**. Module age effect in split half *A* against split half *B*. Large opaque points are the pairs whose age effect was significant in both halves, the only pairs on which direction agreement is assessed. **e**. Cross-cohort transfer of frozen eigengene weights. Each module’s age association is expressed as a *z* score relative to matched random projections in the source and target cohorts. Filled symbols agree in direction and both axes are pseudo-log scaled. **f**. Similarity of matched cross-cohort module pairs relative to size-matched random pairings. Gray denotes the null mean ± SD (standard deviation); *P* values are permutation based. Cell-type-profile similarity is shown only for IsoGraph, for which these profiles were available.

Across six cohort-region analyses spanning three anatomically matched BrainSEQ–GTEx region pairs, 93 of 118 IsoGraph modules showed split-half co-assignment above a size-matched permutation null at FDR < 0.05, compared with 62 of 72 abundance-WGCNA modules (**Fig. 3b**; **Data S5**). Thus, a large majority of modules from both representations were stable across donor subsets, despite IsoGraph producing a finer partition. Module internal architecture was also reproducible: IsoGraph switch-driver loadings correlated strongly across halves (regional median Spearman ρ = 0.72–0.82; **Fig. 3c**). For age-associated modules, effect directions agreed for 22 of 23 IsoGraph pairs and all 14 WGCNA pairs (**Fig. 3d**). These split-half patterns were similar across Leiden^24^ resolutions from 0.5 to 20 (mean adjusted Rand index 0.37–0.47; 0.42 at the production resolution), and no single resolution performed best in every region, indicating that module stability was not strongly dependent on the chosen production resolution (**Data S6**).

We next asked whether module axes learned in one cohort retained age-related structure when transferred to the other cohort. Source-cohort module weights were frozen and applied unchanged to aligned features in the target cohort, and each projected age association was standardized against matched random projections. Nearly all projected modules exceeded their matched random expectations (37 of 38 BrainSEQ-to-GTEx and 53 of 55 GTEx-to-BrainSEQ projections for IsoGraph; 49 of 52 and 10 of 10 for WGCNA). IsoGraph projections also showed strong directional concordance of the null-standardized age associations, with agreement in 33 of 38 BrainSEQ-to-GTEx and 44 of 55 GTEx-to-BrainSEQ projections (*P* = 2.13 × 10^−6^ and 4.35 × 10^−6^), compared with 28 of 52 and 3 of 10 WGCNA projections (**Fig. 3e**; **Data S7**). Thus, frozen IsoGraph axes preserved age-associated structure across cohorts even when the raw regression coefficient was not directly comparable between datasets. Because IsoGraph and WGCNA use different feature representations, this comparison reflects transferability of the overall representation rather than network inference alone.

Cross-cohort reproducibility was stronger for transferred module axes than for independently reconstructed gene partitions. Gene-membership-matched modules showed no excess of age-concordant pairs over the permutation null (IsoGraph 1/38, *P* = 0.91; WGCNA 2/52, *P* = 0.15; **Fig. S16**; **Data S8**). Nevertheless, matched IsoGraph modules shared more functional annotation and cell-type profiles than expected under size-matched re-pairing (Gene Ontology [GO] term Jaccard 0.097 versus 0.030, *P* = 0.001; cell-type-profile correlation 0.208 versus 0.040, *P* = 0.006; **Fig. 3f**; **Data S9**). Thus, cross-cohort reproducibility was stronger for the source-derived module axis and its associated biological structure than for exact recovery of the same gene partition.

### Module context predicts held-out DTU beyond gene-wise evidence and co-expression

Switch-module genes first showed concordance with an established gene-wise DTU model: continuous satuRn^8^ evidence was elevated among switch-module genes in 11 of 13 testable analyses (rank-biserial effect sizes 0.04–0.45; **Fig. S17**; **Data S10**), indicating that module assignment agreed with independent gene-wise DTU evidence. We next tested whether switch-module genes were also supported by junction-level usage measured from the underlying short-read alignments (percent-spliced-in [PSI] events in BrainSEQ and within-gene junction usage in GTEx). Switch-module genes were enriched for genes with junction-level age associations in five of the nine aging analyses containing switch-module genes (transcript-count-adjusted odds ratio 2.24–3.28; *P* ≤ 0.0032; **Data S11**).

IsoGraph module context predicted held-out DTU beyond a gene’s own discovery evidence and conventional co-expression in three of six cohort-region analyses. To evaluate this across three matched regions in BrainSEQ and GTEx, we repeatedly split donors into discovery and held-out halves and estimated DTU independently in each. For each gene, we then tested whether the discovery-half DTU evidence of its module neighbors predicted its held-out DTU, adjusting for the gene’s own discovery evidence, isoform count, overall expression, minor-isoform usage, and DTU evidence among its most co-expressed genes. After these adjustments, module context remained associated with held-out DTU in BrainSEQ caudate (mean partial *r* = 0.056; 8/10 directions positive), GTEx hippocampus (*r* = 0.074; 9/10), and GTEx frontal cortex BA9 (*r* = 0.086; 10/10), with permutation *q* = 0.002 in each analysis (**Fig. S18a**; **Data S12**). This relationship was not detected in the other three cohort-region analyses, indicating that the predictive contribution of module context was cohort- and region-dependent.

In the same three positive analyses, module context was enriched for held-out DTU among genes that fell below the discovery-half significance threshold (Benjamini–Hochberg *q* ≥ 0.05; per-analysis median adjusted odds ratios 1.39–1.72; **Fig. S18b**). This association persisted after removing the largest module in each split. Together, these analyses show that IsoGraph module context captures held-out DTU information not contained in a gene’s own discovery evidence or conventional co-expression, with reproducible gains in three of six cohort-region analyses.

### Age-associated switches recurrently distinguish transcripts with different regulatory and coding architectures

We next asked whether age-associated switch pairs preferentially distinguished transcripts with different molecular architectures. Across the nine aging analyses containing annotatable age-associated switch genes, UTR differences were enriched relative to random within-gene transcript pairs in all nine analyses (pooled enrichment 1.247, 95% CI 1.22–1.28), while coding-sequence differences were enriched in eight of nine analyses, although more weakly (1.040; **Fig. 4a**). Exon-boundary differences were not enriched, whereas differences in nonsense-mediated-decay annotation, transcript biotype, and coding status were depleted. UTR and coding-sequence enrichment was similar among switches from GO-visible and GO-invisible modules (**Fig. 4b**), indicating that these structural patterns were not restricted to modules with readily detectable functional annotation.

**Fig. 4.**
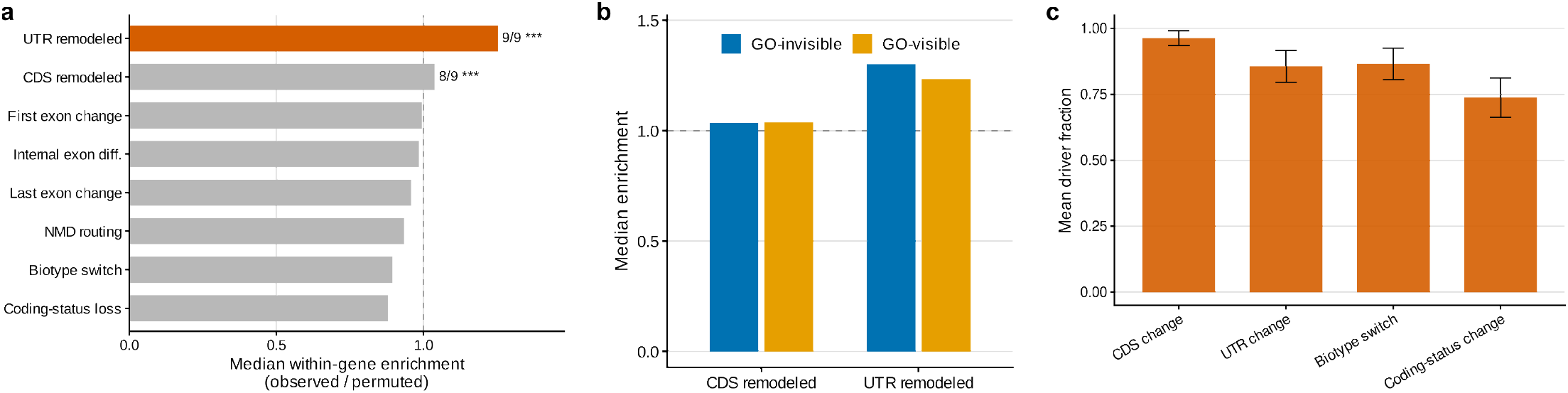
Molecular architecture of age-associated transcript switches. **a**. Enrichment of each structural difference among switch pairs relative to random transcript pairs from the same genes, as the median observed/permuted ratio across the nine annotatable aging analyses. Labels give the number of analyses with nominal enrichment (*P* < 0.05); asterisks mark the Fisher-combined *P* (^***^ < 0.001). The dashed line marks no enrichment. **b**. The coding-sequence and untranslated-region enrichments of **a**, split according to whether individual switch modules showed Gene Ontology (GO) enrichment; eight analyses contained GO-visible switches and six contained GO-invisible switches. **c**. Descriptive fraction of top-ranked driver transcript pairs differing in each structural class across 19 age-associated trusted modules; error bars show SEM (standard error of the mean) across modules. Enrichment relative to within-gene null pairs is evaluated in **a**.

Switch genes were also more loss-of-function constrained than the genome-wide background (median gnomAD^25^ loss-of-function observed/expected upper-bound fraction [LOEUF] of 0.729 across the ten annotatable analyses). In contrast, exons unique to one member of a switch pair were depleted for ClinVar^26^ pathogenic or likely-pathogenic entries relative to other exons of the same transcripts (median density ratio 0.16; depletion in 10 of 10 analyses; **Fig. S19**; **Table S5**). Thus, switch genes tended to occur in constrained loci, while the alternatively represented exons themselves were not preferentially enriched for known pathogenic variation.

Consistent with the enrichment analyses, top-ranked driver pairs from the 19 age-associated trusted modules frequently differed in coding sequence (mean fraction 0.96) and in UTR structure, transcript biotype, or coding status (0.74–0.87; **Fig. 4c**). Exploratory motif and enhanced crosslinking and immunoprecipitation (eCLIP) analyses^27,28^ further nominated candidate *trans*-acting RNA-binding proteins (RBPs) (**Fig. S20**; **Data S13** and **S14**). Together, these analyses show that age-associated IsoGraph switches recurrently distinguish transcripts with different coding and regulatory architectures, providing a molecular basis for tracing network-level usage signals to specific transcript consequences.

### Genetic analyses support *cis* regulation and prioritize disease-associated isoform switch events

We first asked whether IsoGraph transcript events showed within-donor evidence of *cis* regulation. Allele-aware junction recounting identified significant haplotype differences in relative support for isoform-discriminating junctions for 590 switch pairs in 185 genes in caudate, 247 pairs in 92 genes in hippocampus, and 98 pairs in 37 genes in DLPFC (FDR < 0.05; **Fig. 5a**; **Data S15**). These associations were well calibrated across negative controls and alternative statistical tests, and their directions agreed with between-donor genotype-dosage associations for 82–87% of significant pairs (**Fig. S21a–d**). Because the allelic contrasts compare haplotypes within the same donor, they provide evidence for *cis*-regulated transcript choice that is robust to between-donor differences in cellular composition.

**Fig. 5.**
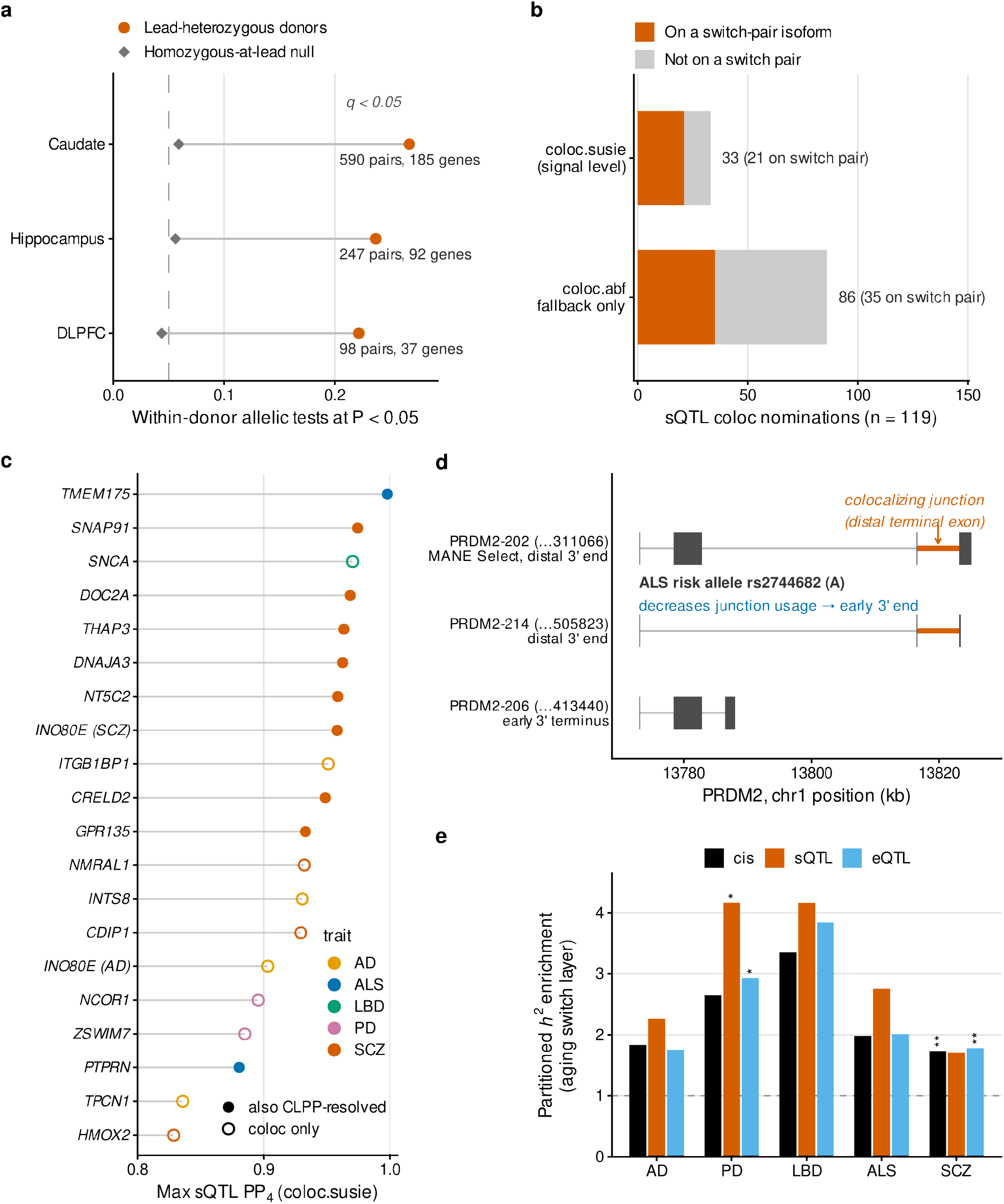
Genetic analyses support *cis* regulation of isoform choice and prioritize disease-associated switch events. **a**. Within-donor allelic test in BrainSEQ caudate, hippocampus, and dorsolateral prefrontal cortex (DLPFC). For each switch pair, fragments crossing isoform-specific junctions were counted on the two haplotypes of donors heterozygous at the lead variant associated with the switch and compared with a beta-binomial mixed model (*Methods*). Orange points give the fraction of fitted pairs with nominal *P* < 0.05; gray points give the same fraction for donors homozygous at the lead, which should fall at 0.05 (dashed line). Labels give the pairs and genes at Benjamini–Hochberg-adjusted *P* (*q*) < 0.05; calibration checks are in **Fig. S21. b**. The 119 splicing quantitative trait locus (sQTL) colocalization nominations (posterior probability of a shared causal variant [PP4] ≥ 0.8) across five genome-wide association study (GWAS) traits, by estimator tier: signal level `coloc.susie`in at least one tissue) or the `coloc.abf` fallback only. Orange marks nominations whose colocalizing junction is carried by an isoform of a tissue-matched IsoGraph switch pair (56 nominations, 115 events). **c**. Maximum sQTL PP4 for the 20 switch-mapped events evaluated with `coloc.susie`, colored by trait. Filled points also met the switch-mapping criterion of the credible-set posterior-overlap score (CLPP_overlap_). Events supported only by `coloc.abf` are not shown. **d**. *PRDM2* at the amyotrophic lateral sclerosis (ALS) locus. The colocalizing junction (orange) joins the distal terminal exon and is spliced by *PRDM2-202* (MANE Select) and *PRDM2-214*. The ALS risk allele A at rs2744682 was associated with reduced usage of the distal terminal junction, corresponding to relatively greater representation of early-terminating transcript structures. CLPP_overlap_ is 0.056 and the junction-to-switch correlation is 0.11. **e**. Partitioned heritability enrichment of variants that are Genotype-Tissue Expression (GTEx) brain QTLs of the aging switch genes (sQTLs, expression QTLs [eQTLs], or either), each tested in a single-annotation model conditional on baselineLD v2.2. Stars denote Benjamini–Hochberg-adjusted significance across the 15 single-annotation tests (^*^ *q* < 0.05, ^**^ *q* < 0.01). Joint sQTL/eQTL models are in **Data S22**.

We next asked whether genetically regulated transcript events intersected disease-associated variation. We colocalized GTEx brain splicing quantitative trait loci (sQTL) and expression QTL (eQTL) signals with genome-wide association studies (GWAS) of Alzheimer’s disease^29^, Parkinson’s disease^30^, dementia with Lewy bodies^31^, amyotrophic lateral sclerosis (ALS)^32^, and schizophrenia^33^. We prioritized signal-level colocalization when credible sets could be resolved for both traits and otherwise used regional colocalization under a single-causal-variant assumption (*Methods*). At a posterior probability of a shared causal variant (PP4) ≥ 0.8, sQTL colocalization nominated 119 gene-by-trait pairs in 116 genes, including 33 supported at signal level in at least one tissue (**Fig. 5b**; **Data S16**).

Across all 6,342 jointly evaluated gene-analysis combinations, eQTL-only colocalization was more frequent than sQTL-only colocalization (160 versus 67; McNemar *P* = 5.9 × 10^−10^; **Data S17**). A similar asymmetry was observed for IsoGraph-derived coordinates in BrainSEQ, with 16 genes colocalizing only on the switch axis and 94 only on the abundance axis (McNemar *P* = 1.3 × 10^−14^; **Fig. S22a**; **Data S18**). At the level of QTL architecture, module genes were depleted of both *cis*-eQTLs and *cis*-sQTLs relative to the tested background (pooled odds ratios 0.84 and 0.87 across 17 analyses), although the paired within-analysis sQTL-to-eQTL odds-ratio ratio was modestly above one (1.03, 95% CI 1.01–1.06, *P* = 0.018 across all modules; 1.07, 1.01–1.12, *P* = 0.020 in phenotype-associated modules). A matched multiplex WGCNA comparator showed a similar contrast (1.08, 1.03–1.15, *P* = 0.0044), indicating that this relative sparing of sQTLs reflected the paired feature representation rather than IsoGraph network inference alone (**Fig. S23**; **Tables S6** and **S7**). These analyses therefore identify a subset of disease loci with splicing-related genetic evidence within a broader landscape dominated by abundance-associated regulation.

Disease-colocalizing splicing signals mapped directly to IsoGraph switch-pair transcripts in 55 genes, representing 56 of the 119 gene-by-trait nominations. Twenty-one were signal-level nominations, and for 20 the switch-mapped event itself was evaluated (**Fig. 5b,c**; **Data S19**). Summary-data-based Mendelian randomization (SMR)^34^ passed its Bonferroni-corrected threshold for 278 of 313 instrumented primary sQTL probes, while the heterogeneity in dependent instruments (HEIDI) test rejected a single shared-variant model for 13 and did not reject it for 263 (**Fig. S22b**; **Data S20**). A complementary credible-set posterior-overlap score (CLPP_overlap_), calculated from shared GWAS and QTL posterior inclusion probabilities (PIPs) following the posterior-product principle of eCAVIAR^22^, nominated 160 genes at CLPP_overlap_ ≥ 0.01. Thirty had sQTL events mapping to tissue-matched switch pairs, including 12 that also belonged to the colocalization-derived switch-mapped set (**Data S21**). Together, these complementary analyses converged on a subset of disease-associated loci in which genetic evidence could be traced to specific IsoGraph transcript events.

Among these events, *PRDM2* showed the strongest convergence across the evaluated evidence types as a candidate transcript-choice association with ALS (**Fig. 5d**). Nine colocalization events in seven genes mapped to a switch pair in both GTEx and the anatomically matched BrainSEQ region; *PRDM2* had the highest PP4 among these and was the only event also supported by CLPP_overlap_ ≥ 0.05. At the ALS locus, cortex sQTL colocalization yielded PP4 = 0.964 with CLPP_overlap_ = 0.056, whereas eQTL evidence was weaker (PP4 = 0.494). The ALS risk allele *A* at rs2744682 was associated with reduced usage of a distal-terminal-exon junction carried by *PRDM2-202/214*, consistent with relative shifting toward early-terminating 3′ transcript structures represented by *PRDM2-206/203*. These alternative forms were also detected independently in long-read data. The convergence of genetic association, cross-cohort switch mapping, and transcript-level detection therefore prioritizes *PRDM2* alternative terminal-exon usage as a candidate ALS-associated transcript event.

At a broader annotation level, sQTLs linked to aging switch genes were enriched for Parkinson’s disease heritability^35,36^ (4.17-fold; *q* = 0.023 across the 15 single-annotation tests of five traits and three annotations; **Fig. 5e**; **Data S22**). The corresponding eQTL annotation was also enriched when tested alone (2.93-fold; *q* = 0.023); in a joint model, the sQTL coefficient remained nominally significant (*P* = 0.013), whereas the eQTL coefficient did not (*P* = 0.067). Schizophrenia heritability was enriched in the eQTL (1.78-fold; *q* = 0.0097) and combined *cis*-QTL (1.73-fold; *q* = 0.0097) annotations, but not in the sQTL annotation after multiple-testing correction (1.71-fold; *P* = 0.035, *q* = 0.088), consistent with broader *cis*-regulatory rather than splicing-specific enrichment for this trait.

Disease-genetic support was stronger at the level of specific transcript events than at the level of whole modules. Genes with significant within-donor allelic effects were not preferentially concentrated in modules with stronger age associations (pooled correlation between module fraction of allelic-effect genes and age-association rank = −0.183; permutation *P* = 0.47; **Fig. S21e**; **Table S8**), and module-level colocalization-convergence tests were null across all nine testable trait-by-cohort combinations (permutation *P* = 0.20–0.86; **Fig. S24**; **Data S23**). Significant results from competitive gene-set tests using MAGMA across eight brain-disorder GWAS were likewise concentrated among large modules: 24 of the 37 module-by-trait sets significant at FDR < 0.05 contained at least 900 genes, with similar size dependence at Leiden resolution 5.0 and in classical abundance-only WGCNA (**Fig. S25**; **Data S24**). Together, these analyses indicate that genetic support is most clearly resolved at specific transcript events rather than as a general property of entire aging-associated modules.

### Disease-prioritized switch events show independent transcript-level support

We next asked whether disease-prioritized IsoGraph switch events were independently supported by transcript-level measurements. Among genetically prioritized genes, switch pairs more often showed reciprocal usage in independent long-read data than abundance-matched IsoGraph pairs from other genes (58.1% versus 24.5%; empirical *P* = 5 × 10^−4^; **Fig. 6a,b**; **Data S25**). We evaluated an Oxford Nanopore dataset from human DLPFC (*n* = 12 donors; six with Alzheimer’s disease, six controls) quantified with Bambu^18,37^. The analysis focused on 55 genetically prioritized genes in which a disease-colocalizing sQTL junction was carried by at least one transcript in a tissue-matched IsoGraph switch pair. After requiring long-read detection of both transcripts, 191 switch pairs were testable.

**Fig. 6.**
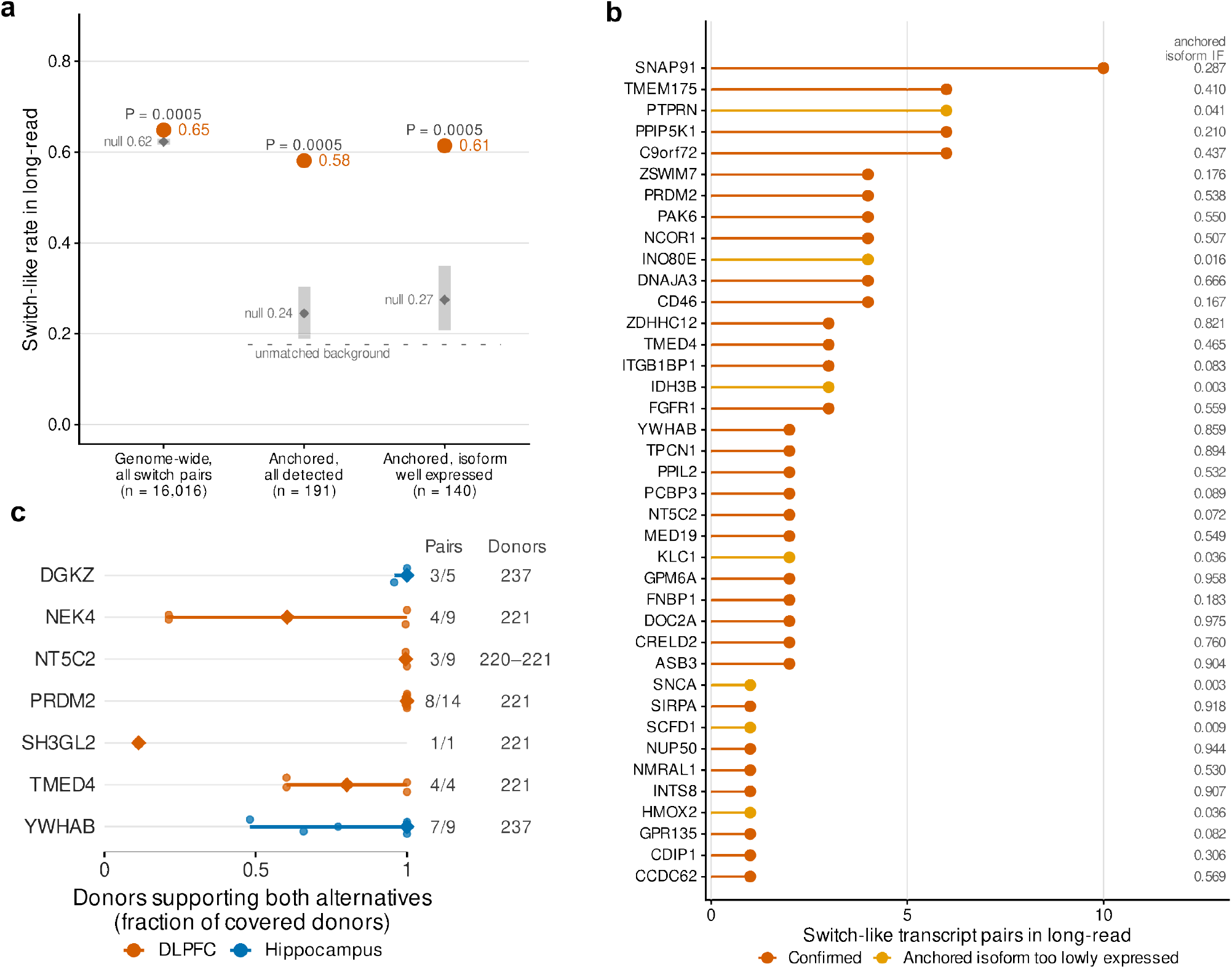
Disease-prioritized IsoGraph switch events show independent transcript-level support. **a**. Switch-like reciprocal usage in Oxford Nanopore long-read data from dorsolateral prefrontal cortex (DLPFC; *n* = 12 donors). A transcript pair was classified as switch-like when the within-gene fractions of its two isoforms were negatively rank-correlated across donors meeting the detection criteria (*Methods*). Shown are all detected IsoGraph switch pairs genome-wide, the 191 detected transcript pairs from 55 colocalization-prioritized genes, and the subset of 140 prioritized pairs whose genetically anchored isoform had a mean within-gene fraction ≥ 0.05. Orange points show observed switch-like rates; gray intervals show the 95% range from 2,000 abundance-matched random draws, and the dashed line marks the unmatched genome-wide background rate. Empirical *P* values are two-sided; 5 × 10^−4^ is the permutation floor. **b**. Gene-level distribution of long-read-supported switch pairs among the 55 prioritized genes. Bars show the number of switch-like transcript pairs per gene for the 39 genes with at least one supported pair. Genes are distinguished according to whether at least one supported pair contained a well-expressed anchored isoform (mean within-gene fraction ≥ 0.05) or whether support was observed only below this threshold; the 16 genes without a switch-like pair are listed below the axis. The right-hand column shows the highest mean within-gene fraction of an anchored isoform for each gene. **c**. Short-read junction corroboration of genetically prioritized transcript pairs in BrainSEQ DLPFC (orange) and hippocampus (blue). For each measurable transcript pair (at least 30 covered donors, each with at least 10 pair-discriminating fragments), small points show the fraction of covered adult control donors with at least one fragment supporting each transcript alternative; diamonds show gene–region medians and horizontal lines the observed range across measured pairs, not confidence intervals. Labels report the number of measured versus targeted pairs and the number of covered donors.

A pair was classified as switch-like when its two transcripts showed a negative rank correlation in within-gene fraction across donors in which both were measured (*Methods*). Overall, 111 of 191 testable pairs met this criterion, compared with 24.5% of abundance-matched switch pairs from other genes. The enrichment was similar among the 140 pairs with a well-expressed genetically anchored isoform (86 pairs, 61.4%; matched null 27.5%), and 39 of the 55 prioritized genes contained at least one switch-like pair, including 32 at the well-expressed threshold (**Fig. 6a,b**; **Data S25**). These independent long-read measurements therefore support both the presence and reciprocal usage of transcripts implicated by disease-prioritized IsoGraph switches.

Short-read junction measurements in BrainSEQ provided complementary alignment-level support for the prioritized transcript events (**Fig. 6c**; **Data S26**). Among the 55 prioritized genes, 26 had a matched BrainSEQ regional counterpart, and 30 transcript pairs across seven genes met the junction-model and coverage requirements for evaluation (*Methods*). Of these, 29 of 30 showed junction support for both transcript alternatives in at least 30 donors, with a gene-balanced mean donor support of 0.76. The implicated sQTL junction itself discriminated the two transcript alternatives for 26 of 30 pairs, and junction-based and transcript-quantification usage were positively concordant across donors (median covariate-adjusted Spearman ρ = 0.32). These patterns were similar under more stringent fragment-depth thresholds and after excluding proxy regional mappings. Because the junction and transcript measurements derive from overlapping short-read data, this analysis provides orthogonal representation of the transcript structures rather than independent cohort replication. Together with the long-read analysis, these results connect disease-associated QTL signals to alternative transcript usage observed across complementary sequencing representations.

## Discussion

IsoGraph revealed reproducible age-associated transcript-usage programs in the adult human brain that were distinct from coordinated changes in total gene abundance. The framework represents within-gene transcript composition and total abundance as paired coordinates within the same gene network. Across adult brain aging analyses, these coordinates were weakly correlated within genes, and age associations carried by transcript switching were detected in multiple regions after conditioning on abundance. A subset persisted after adjustment for estimated cellular composition. These findings show that coordinated transcript choice captures age-associated variation beyond total abundance that can be obscured by gene-level expression summaries.

This contribution is distinct from the broader application of network analysis to isoform regulation. Previous studies have constructed gene-expression, isoform-expression, and isoform-usage networks^12^, identified co-differential isoform usage across cell types^13^, built isoform co-expression networks in human cortex^14^, and integrated total-expression and relative-isoform features within common transcriptomic graphs^10^. IsoGraph adds a gene-centered decomposition: for every gene, it retains paired abundance and switching coordinates, quantifies the contribution of each channel to shared module organization, and connects the switching component back to the transcripts and structural features from which it arises. This paired representation directly asks whether shared network organization is driven by transcript choice or abundance, whether that distinction is preserved across cohorts, and which transcript events underlie the network-level signal. The synthetic benchmarks defined this specificity: IsoGraph preferentially recovered structure driven by coordinated transcript usage, while performance shifted toward abundance-based representations when abundance or degradation dominated the signal.

The reproducibility analyses further distinguished transfer of a coordinated molecular axis from recovery of identical module membership. Within cohorts, most IsoGraph modules were stable across donor splits, and switch-driver loadings were reproducible. Across cohorts, frozen module weights generally retained the direction of their age association, standardized against matched random projections, when applied without refitting. Independently reconstructed modules, by contrast, showed no excess of age-concordant gene-membership matches, although matched modules shared more functional and cell-type annotation than expected. This distinction is consistent with broader work on module preservation and transfer of latent molecular patterns^38,39^. Together, these results indicate that the transferable unit of these transcriptomic programs is the coordinated switch-and-abundance axis rather than an invariant list of genes.

Module context also carried information beyond gene-wise DTU^8^. In three of six cohort-region analyses, module-neighbor DTU evidence predicted held-out gene-wise DTU beyond each gene’s own discovery evidence, conventional co-expression and baseline transcript features. Network context can therefore contain out-of-sample information about transcript usage, although this contribution was region- and cohort-dependent. Cellular composition likewise influenced the signal differently across regions: some transcript-usage associations persisted after adjustment for estimated cell-type proportions, whereas others were attenuated and the set of detected genes changed. IsoGraph modules were not generally enriched for the tested cell-type marker sets. This regional heterogeneity indicates that IsoGraph captures both composition-sensitive and composition-robust transcript-usage structure. Bulk tissue, however, cannot determine whether composition-sensitive effects reflect molecular changes within cells, shifts in cellular populations, or both^40^.

Structural annotation and orthogonal transcript measurements connected these network-level signals to underlying transcript events. Age-associated switch pairs were consistently enriched for UTR differences and, more weakly, for coding-sequence differences, whereas differences in nonsense-mediated-decay annotation were depleted. This pattern is compatible with the prominence of alternative transcription start and termination sites among human isoform differences^41^ and identifies transcript features with plausible consequences for RNA stability, localization, or translation^42^. Switch genes were more loss-of-function constrained than the genome-wide background while their switch-specific exons were not enriched for known pathogenic variation, suggesting that age-associated transcript choice preferentially alters regulatory and coding architecture within genes that are themselves relatively constrained. Independent long-read DLPFC data^18^ further supported the genetically prioritized subset: anchored transcript pairs showed switch-like reciprocal usage more often than abundance-matched background pairs. Short-read junction usage provided a complementary check at the gene level in several aging analyses and supported transcript structures where individual events could be resolved. Together, these measurements provide convergent transcript-level support for the events represented by composite IsoGraph switch coordinates.

Genetic analyses revealed a separation between disease-associated transcript events and whole-network aging programs. Within-donor allelic imbalance supported *cis* regulation of isoform choice and agreed directionally with between-donor genotype-dosage associations while reducing sensitivity to between-donor differences in cellular composition^19^. Colocalization identified disease-linked sQTL events, including a subset supported at signal level^43^, but these event-level associations did not aggregate into evidence that aging modules share a predominantly splicing-mediated genetic architecture. eQTL colocalization was more frequent overall, the relative sparing of sQTLs in phenotype-associated modules was not specific to IsoGraph network inference, and module-level colocalization and gene-set analyses provided little evidence for coherent disease-genetic enrichment beyond effects associated with large modules. Heritability analyses likewise distinguished trait-specific patterns: Parkinson’s disease showed enrichment among both sQTL and eQTL annotations linked to aging switch genes, whereas schizophrenia showed broader *cis*-regulatory enrichment. Disease-associated genetic variation therefore resolved more convincingly at constituent transcript events than at the level of entire aging programs. This separation is biologically informative: reproducible network organization can emerge from coordinated molecular variation without requiring a single shared genetic architecture across its constituent genes.

*PRDM2* illustrates how convergence across these evidence layers can prioritize a specific disease-associated transcript hypothesis. *PRDM2* showed the strongest convergence across genetic, cross-cohort, and long-read evidence, consistent with altered 3′ terminal-exon choice at an ALS locus, although the colocalization required a single-causal-variant assumption because no GWAS credible set was available. A recent preprint reported experimentally confirmed cryptic-exon inclusion in *PRDM2* in human forebrain organoids carrying the TDP-43 K181E mutation associated with ALS and frontotemporal dementia (FTD)^44^. Although distinct from the terminal-exon event identified here, this finding independently implicates *PRDM2* RNA processing in an ALS/FTD context. Together, these observations prioritize altered *PRDM2* terminal-exon choice for mechanistic investigation. More broadly, convergence across genetic localization, transcript mapping, and orthogonal molecular measurements can move a network association toward a specifically defined transcript event that can be tested experimentally.

Several limitations define the scope of these conclusions. First, at the level of transcript measurement, short-read quantification remains uncertain for highly similar isoforms and is sensitive to sequencing and quantification choices^45^. Junction measurements provide a complementary view of transcript structure but are not fully independent because they derive from overlapping read alignments. Second, at the level of tissues and cohorts, BrainSEQ and GTEx differ in cohort composition and quantification pipeline, and the WGCNA comparison reflects differences in feature representation as well as network inference. Differences in reconstructed module membership may therefore reflect both biological and technical variation. Bulk tissue also cannot cleanly distinguish within-cell transcript regulation from shifts in cellular composition, and deconvolution results depend on the reference and method; matched composition estimates were unavailable for several GTEx regions. The independent long-read cohort was small and restricted to DLPFC, providing orthogonal support for transcript behavior but not a direct test of age dependence or switching magnitude. Third, genetic interpretation depends on QTL coverage, linkage disequilibrium (LD), fine-mapping assumptions, and instrument availability. Mapping an sQTL junction to an IsoGraph switch pair links genetic regulation to the implicated transcript structure but does not establish that the mapped junction itself generates the composite switch coordinate. More broadly, the molecular and genetic evidence prioritizes transcript-level hypotheses but does not by itself establish downstream effects on RNA stability, translation, or protein output.

IsoGraph represents transcript choice and gene abundance as paired contributions to gene-network organization. In adult human brain, this framework identified reproducible age-associated transcript-usage programs whose molecular axes transferred across cohorts more readily than exact gene partitions, while genetic evidence resolved most clearly at the level of individual transcript events. These network signals could in turn be traced to specific transcript structures and disease-associated loci. By separating coordinated transcript choice from coordinated abundance within the same network, IsoGraph reveals transcript-level organization that gene-abundance networks alone do not resolve and provides a route from reproducible network structure to specific molecular hypotheses for functional investigation.

## Methods

### Study design

We developed IsoGraph to identify modules of genes with coordinated changes in relative transcript usage while retaining total gene abundance as a separate feature channel. The study combined synthetic benchmarks, module discovery in adult postmortem human brain RNA sequencing, evaluation of reproducibility across donors and cohorts, orthogonal transcript-level validation and genetic analyses. Analyses followed a committed, stage-wise workflow and are described below in the order of the Results: synthetic benchmarking, module discovery and characterization, stability and cross-cohort transfer, held-out and orthogonal validation of switch events, structural annotation, within-donor allelic tests, colocalization and heritability analyses, and candidate regulator nomination. Unless otherwise stated, hypothesis tests were two-sided, and the FDR was controlled using the Benjamini–Hochberg procedure^46^.

We distinguish four uses of switching throughout these analyses. A switch coordinate summarizes within-gene transcript composition. Module-member genes belong to an IsoGraph module; membership alone does not establish a gene-level usage association or an active switch-channel role. In the genetic analyses, “selected switch genes” denotes all members of phenotype-associated modules, irrespective of channel role. Candidate transcript pairs are enumerated contrasts for downstream annotation and testing, not necessarily anticorrelated transcripts or transcripts retained during network fitting. A pair is termed switch-like in the long-read analysis only when its measured within-gene fractions are negatively rank-correlated under the criteria below.

No participants were prospectively recruited and no new biospecimens were collected. BrainSEQ and GTEx data were obtained under the consent, institutional review, and data-access conditions of the original studies^15,16,47^. Analyses used de-identified data. Controlled-access BrainSEQ molecular and genotype data were analyzed under the applicable data-use agreements.

### BrainSEQ cohorts

BrainSEQ sample eligibility differed by analytical purpose. **Table 1** separates discovery cohorts, genotype-matched mapping cohorts, and the available allele-specific expression (ASE) resource; counts are ordered as caudate, hippocampus, and DLPFC.

Allelic-test eligibility was determined separately for each pair by informative fragment counts and lead-heterozygous donor coverage, as specified below.

BrainSEQ analyses used bulk RNA sequencing from postmortem caudate nucleus, hippocampus, and DLPFC. The source resource included phenotype and clinical metadata, raw paired-end RNA-sequencing reads, region-level sequencing quality metrics, and SNP-array genotypes. We restricted each region to its RiboZero rRNA-depletion dataset: BrainSEQ Phase 3 caudate, BrainSEQ Phase 2 hippocampus and BrainSEQ Phase 2 DLPFC. This restriction excluded the overlapping Phase 1 poly(A)-selected DLPFC libraries. Caudate and DLPFC libraries used RiboZero Gold; the hippocampus dataset combined RiboZero HMR (195) and RiboZero Gold (43) libraries (**Data S1**).

Samples were first required to occur in both the transcript-count matrix and the RNA-sequencing metadata. We then retained the canonical region-specific dataset, age at death ≥18 years and one of the prespecified diagnosis labels. The aging analyses were restricted to neurotypical controls. They contained 238 caudate samples (71 female and 167 male; age 18.05–89.91 years), 238 hippocampus samples (72 female and 166 male; age 18.04–84.16 years) and 222 DLPFC samples (66 female and 156 male; age 18.04–84.16 years). The schizophrenia analysis retained controls and individuals diagnosed with schizophrenia in caudate and contained 390 samples (238 controls and 152 cases; 121 female and 269 male; age 18.05–96.92 years). Samples with other diagnoses, samples marked for exclusion by the BrainSEQ quality-control flag, non-adult donors and libraries from a noncanonical region/ preparation dataset were excluded.

Each donor contributed at most one RNA-sequencing sample within a region-specific analysis. Donors could occur in more than one region (**Fig. S11**); all discovery and phenotype models were therefore fit separately by region. Genetic analyses additionally required an unambiguous RNA-to-genotype donor mapping and available TOPMed-imputed^48^ genotypes. Analyses requiring an RNA-sequencing quality covariate used samples with the corresponding region-level metric; missing continuous discovery covariates were handled as described below rather than filtering on the tested phenotype.

For same-tissue BrainSEQ switch-QTL and abundance-QTL mapping, switch and abundance phenotypes were recomputed in a larger, prespecified genotype-matched cohort rather than reusing the control-only discovery matrix. These analyses retained the canonical region-specific RiboZero dataset, age >13 years, all available diagnoses, and required presence in the current TOPMed genotype panel. Diagnosis was included as a QTL covariate. The primary within-gene modality contrast used all qualifying donors; a European-ancestry subset with ancestry-specific genotype principal components was used for colocalization with European-ancestry GWAS. This same-cohort QTL analysis was treated as endogenous genetic anchoring, not independent replication.

BrainSEQ reads were aligned to GRCh38.p14 with GENCODE release 47 annotation^49^ using STAR v2.7.11b^50^. A two-pass procedure first discovered splice junctions across samples, removed junctions supported only by multimapping reads or by fewer than two uniquely mapped reads, and supplied the retained junctions to the second pass. Alignment quality was assessed with RNA-SeQC v5.0.4 ^51^. Gene counts were obtained from coordinate-sorted BAM files with featureCounts v2.0.8 using paired-end, reverse-stranded exon counting against the GENCODE v47 GTF `-p --countReadPairs -s 2 -t exon -g gene_id`)^52^. Transcript counts and transcripts per million (TPM) were estimated with Salmon v1.9.0 using inward-facing, reverse-stranded libraries and validated mappings `-l ISR --validateMappings`)^2^. Count matrices, feature annotation and quality-control metrics were stored as `RangedSummarizedExperiment` objects^53^.

BrainSEQ SNP-array genotypes were quality controlled by array platform with PLINK v1.9, harmonized to 1000 Genomes Project Phase 3 sites, lifted from GRCh37 to GRCh38 with CrossMap, normalized with bcftools and imputed separately by platform through the TOPMed Imputation Server using Eagle2 pre-phasing and Minimac4 against TOPMed Freeze 10^48,54–63^. Downstream analyses retained variants with imputation INFO *R*^2^ ≥0.8. Autosomal variants used for ancestry principal components had minor-allele frequency, ≥0.05 missingness ≤ 0.05 and Hardy–Weinberg *P* > 10^−6^, and were pruned for LD within chromosome with PLINK2^55^ `--indep-pairwise 200 100 0.2`). The first ten approximate principal components (PCs) were calculated after merging pruned variants across chromosomes 1–22.

### BrainSEQ ASE data generation

BrainSEQ ASE data were generated from matched paired-end RNA-sequencing reads and phased genotype data on GRCh38. The genome FASTA, sequence dictionary, GENCODE v47 annotation, STAR index, and genotype VCFs used the same reference build and chromosome naming. A frozen sample manifest linked each RNA sample to its donor and genotype sample and recorded the FASTQ files, final BAM, sex, region, and processing exceptions. Before counting, genotype sample names and RNA/ genotype mappings were checked, BAMs were required to have an index and to pass `samtools` `quickcheck`^58^, and heterozygous sites were required to be callable in the matched genotype.

Paired-end FASTQ files were the primary input. Reads were aligned with the region-specific two-pass STAR workflow described above, with read-group information retained and coordinate-sorted BAM output. The ASE-final alignments were generated with STAR’s WASP-aware mode `--waspOutputMode SAMtag`) to control reference-mapping bias^64^. Reads overlapping polymorphic sites carried a `vW`tag; only primary, mapped, uniquely aligned reads `NH:i:1`) passing WASP `vW:i:2`(`vW:i:1`) were eligible for ASE interpretation. Reads or fragments with on either mate were excluded. Duplicate flags were retained and summarized for quality control rather than silently removed. A conventional two-pass BAM without WASP-aware handling was not considered ASE-final when the source FASTQs were available.

For each sample, high-confidence heterozygous variants were extracted from the matched, phased genotype VCF with `bcftools`; multiallelic or low-quality sites excluded by genotype quality control were not counted. Site-level reference and alternate reads were quantified from the ASE-grade BAM with GATK `ASEReadCounter`^19,65^ using the GRCh38 reference, minimum depth 1, mapping quality 255 and base quality 10. The same counting policy was applied to all samples. Site counts were aggregated with phASER^66^ using the phased genotypes and GENCODE annotation to produce per-sample allelic counts, haplotype blocks, haplotypic counts, variant connections, and feature-level allelic-expression summaries. phASER’s genome-wide phased genotype `PW`) defined haplotype 1 and haplotype 2 in downstream analyses.

The model-ready release contained cohort-level `site_ase`, `gene_ae`, `haplotypes` `allelic_counts`, `haplotypic_counts` and `variant_connections`, tables together with a column-name map. Quality-control tables recorded the validated sample manifest, alignment and duplicate metrics, unique-alignment proxies, WASP pass/fail counts, genotype heterozygosity, the number and depth of counted ASE sites, allele-balance distributions, phASER yield, and RNA/genotype concordance. The exact FASTA, GTF, STAR index, genotype VCF, software versions, and input checksums were recorded with the release. Per-sample ASE and phASER outputs were retained until cohort aggregates had passed row-count validation and checksum-based archival.

For the junction-specific allelic analysis, the validated resource comprised 486 caudate, 498 DLPFC and 451 hippocampus samples with an ASE-grade BAM, corresponding BAM index and phASER VCF. These broader region-level ASE resources were used only after intersecting donors with the sample and genotype requirements of each downstream test; their sizes therefore differ from the adult control-only IsoGraph discovery cohorts.

### GTEx brain cohorts

We analyzed GTEx v11 RSEM transcript expected counts^17,47,67^, RNA-SeQC gene read counts, sample annotations, and subject annotations from 13 brain regions: amygdala, anterior cingulate cortex BA24, caudate basal ganglia, cerebellar hemisphere, cerebellum, cortex, frontal cortex BA9, hippocampus, hypothalamus, nucleus accumbens basal ganglia, putamen basal ganglia, cervical spinal cord C1 and substantia nigra. Region-specific sample sizes after construction and expression filtering ranged from 181 to 300 (**Data S1**). A sample was included when it was present in the region’s transcript expected-count matrix, gene read-count matrix and sample annotation and had the age and sex fields required by the analysis. Matrix order defined sample order, and construction stopped rather than dropping a sample if its gene-count column was absent. GTEx is an adult resource; no diagnosis-based exclusion was applied. Metadata included age, sex, RNA integrity number (RIN), ischemic time, and mapping rate. Exact age, which the release top-codes at 70 years, was used when available and the midpoint of the released age interval was used otherwise. GENCODE release 47 was used for gene and transcript annotation. GTEx transcript expected counts, rather than TPM, were supplied to IsoGraph to match the count scale used for BrainSEQ.

### Analysis bundles and expression filtering

Each cohort-by-region input was converted into a self-contained IsoGraph bundle containing sample metadata, gene and transcript annotations, gene and transcript count matrices, and a provenance manifest. Filtering occurred in two explicit stages. During bundle construction, genes were retained when counts per million (CPM) were at least 1 in at least max(10, [0.10*n*]) samples within the region. BrainSEQ CPM was calculated from gene read counts and GTEx CPM from RNA-SeQC gene read counts. Transcript matrices were then restricted to transcripts mapping to these expressed genes (**Data S1**).

Immediately before module fitting, the same phenotype-blind, switching-oriented prevalence filter based on the principles implemented by DRIMSeq `dmFilter`^68,69^, was applied to all production IsoGraph and matched-feature baseline fits. For each sample, transcript counts were summed within gene. A gene passed when this total was at least 10 in at least 70% of samples. A transcript passed when its count was at least 10 in at least 10% of samples and its share of the corresponding gene was at least 0.10 in at least 10% of samples. Prevalence requirements were rounded upward to the nearest whole sample. The filter was applied without reference to age or diagnosis and used identical code for BrainSEQ and GTEx. Genes left with one retained transcript remained eligible for the abundance channel but did not receive a switch coordinate; genes with at least two retained transcripts received both channels. The same resulting gene universe was supplied to gene-abundance, switch-only and multiplex WGCNA comparisons. Filter definitions, dimensions before and after filtering and retained feature identifiers were written to the bundle or fit provenance.

### IsoGraph software implementation

IsoGraph v0.1.5, the version recorded in the analysis environment (see Statistical analysis and reproducibility), is an open-source Python package distributed under the Apache-2.0 license through GitHub, PyPI, and Zenodo^70^. The package supports Python 3.11–3.14 and exposes a command-line interface for dataset validation, model fitting, benchmark execution, module explanation, and structural annotation. The package depends on NumPy, pandas, SciPy, scikit-learn, NetworkX ^71^, PyArrow, Patsy, and Hydra; PyTorch^72^ is an optional, platform-specific dependency for the variational autoencoder (VAE) backend, and the WGCNA backend requires R and WGCNA^11^.

IsoGraph accepts a versioned dataset bundle containing a manifest, sample metadata, feature tables, and count matrices. The manifest declares matrix orientation and dimensions, transcript-to-gene mappings, sample identifiers and provenance. Input validation checks required columns, matrix dimensions, finite values and agreement among sample and feature identifiers before model fitting. Providing transcript counts activates the gene-local switch channel; providing gene counts activates a distinct abundance channel. IsoGraph implements baseline precision-network, factor-analysis, graph-diffusion, VAE, and WGCNA backends through a shared model interface. The VAE backend was used for the production real-data network, while the other backends were used as benchmark or matched-feature comparators.

Each fit writes machine-readable module assignments, typed edges, module–trait associations, raw feature scores, reconstructed features, module eigengenes, per-gene channel roles, node-fate diagnostics, residualization diagnostics, calibration metadata and, when requested, a VAE checkpoint. Node-fate diagnostics distinguish assignment to a retained module, membership in a component below the minimum module size, loss after switch reliability weighting and isolation below the edge threshold. The `explain-module` workflow calculates gene drivers, transcript polarity, high-versus-low score contrasts and optional VAE decoder or Integrated Gradients attribution^73^. The `annotate-structure` workflow compares switch-pair transcripts with a supplied GTF to label exon, coding sequence (CDS), UTR, biotype, and coding status differences.

### Isoform-switch coordinates

For a retained gene *g* with *k*_*g*_ ≥ 2 transcripts, let *A*_*gtj*_ denote the count for transcript *t* in sample *j*. We added a pseudocount of 0.5 and calculated within-gene transcript usage (Eq. 1) as

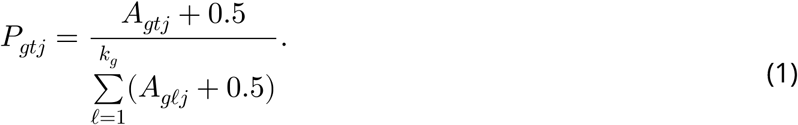

Usage values from Eq. 1 were clipped below 10^−8^at and centered-log-ratio transformed (Eq. 2)^74^:

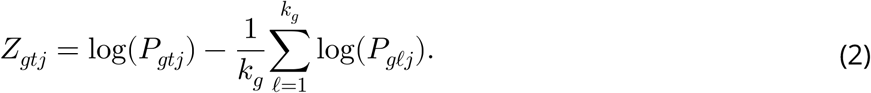

The sample-by-transcript CLR matrix from Eq. 2 was centered across samples and decomposed by singular value decomposition. The first right singular vector defined a gene-local loading axis, and its sample scores defined the switch coordinate *s*_*g*_. Switch coordinates were not scaled to unit variance before network inference. Transcript loadings and transcript–module correlations were retained to map a gene-level coordinate back to its contributing isoforms.

The sign of a singular vector is arbitrary, and no biological direction was imposed on it. Each gene’s axis was instead oriented by a fixed computational rule: the transcript with the largest-magnitude loading was given a positive loading, with ties (as for two-transcript genes, whose loadings are ±*a*) resolved to the first such transcript in annotation order. A higher switch coordinate therefore denotes greater relative usage of that pivot transcript. The rule makes the orientation reproducible for a given transcript set and invariant to sample order, but the pivot, and with it the sign, can differ between cohorts, split-half refits and transcript filters; about 45% of the genes whose axes can be matched between paired BrainSEQ and GTEx analyses carry opposite signs in the two cohorts (**Data S27**). Downstream analyses accordingly treat the sign as a convention rather than a measurement: split-half driver comparisons use absolute loadings, cross-cohort projection re-orients each gene’s axis by the cosine of its shared-transcript loadings, BrainSEQ switch-QTL slopes were pinned to the discovery orientation, and the direction of an age or diagnosis association was interpreted through the retained transcript loadings rather than through the sign of the coordinate. Module inference itself is not invariant to the convention (see Network inference and module definition).

For downstream structural annotation, candidate transcript pairs were enumerated per gene by ranking transcripts by the across-sample variance of pseudocount-regularized usage, retaining at most six transcripts and forming all unordered pairs among them, up to 15 pairs per gene. The structural pair table can therefore contain several pairs per gene; it does not select a single positive-loading versus negative-loading pair, does not require negative pairwise correlation, and ties in variance were not broken by transcript identifier. These pairs are candidate transcript contrasts, and evidence of switching comes from the downstream usage tests rather than from pair enumeration. Pairs were drawn from all bundle transcripts of module genes, that is, after the expressed-gene filter of bundle construction but before the transcript-prevalence filter applied at module fitting, so a pair can include a transcript that did not contribute to the gene’s switch coordinate. The structural-enrichment null likewise sampled annotated transcript pairs of the same genes without a prevalence filter, so observed and null pairs share this property.

For every retained gene, an abundance coordinate was calculated as log_2_(CPM + 0.5) and standardized across samples. Abundance and switch features were stacked into a typed multiplex feature matrix. Switch–switch and abundance–abundance edges used separate thresholds. Switch-to-abundance edges between genes possessing both channels were suppressed so that the two measurements could not substitute for one another; genes with only an abundance channel could still connect to the graph. The switch threshold was fixed at an absolute similarity of 0.50. The abundance threshold was chosen independently within each fit from {0.70,0.75, 0.80, 0.85, 0.90, 0.95} as the smallest threshold that added abundance edges without joining two switch-defined components, including indirect joining through abundance-only genes. If every candidate merged components, 0.95 was used. This trait-blind criterion allowed abundance information to connect genes without collapsing independently identified switch programs.

### Covariate adjustment

Technical and ancestry covariates were residualized from features used for module discovery by ordinary least squares (OLS): RIN, mapping rate, mitochondrial mapping rate, and SNP PCs 1–5 in BrainSEQ, and RIN, ischemic time, and mapping rate in GTEx. Biological covariates and the traits were not removed before clustering. Sex and, in BrainSEQ, manner of death were instead adjusted jointly with the trait in the association models, together with the discovery covariates. Missing numeric covariates were mean-imputed and missing categorical values were represented as an explicit category.

For the VAE, residualization was restricted to module discovery and saved feature scores were unresidualized. Adjusted diagnosis, age-spline and gene-level incremental models included covariates jointly with the trait. The separately reported linear-age Pearson analysis did not include covariates and must not be interpreted as covariate-adjusted inference. Variance retained after residualization and covariate *R*^2^ before and after adjustment were recorded for each feature.

The adjustment strategy depended on the analytical task. Module discovery used the paired switch and abundance features without a trait model, residualized by OLS for the technical and ancestry covariates listed above, so that modules were inferred without removing age or diagnosis. Primary module trait tests related the saved, unresidualized arithmetic-mean module score to a natural age spline or a linear diagnosis term, adjusting for the cohort-specific discovery covariates plus sex and, in BrainSEQ, manner of death; diagnosis models also included age. These tests selected phenotype-associated modules. Gene-level incremental models took one channel as the outcome and the other as a predictor alongside the age spline or diagnosis term, with the same trait-model covariates and, in the composition-adjusted arm, estimated cell fractions; these models identified conditional usage and abundance associations. The split-half and matched-module Pearson analyses related module scores to continuous age without covariates to assess signed age-association concordance. Cross-cohort projection related the frozen principal-component score to continuous age, again without covariates, with correlations standardized against matched random projections to assess transfer relative to each cohort’s random-projection background.

The matched-module sensitivity analysis additionally evaluated linear and spline age models under the covariate modes described in the trait-association section.

### Network inference and module definition

Production real-data analyses used the IsoGraph beta-VAE^75,76^. The encoder and decoder were multilayer perceptrons with ELU activations and no dropout. Two hidden layers contracted from 256 to 128 units in the encoder and expanded symmetrically in the decoder. The encoder returned Gaussian posterior means and log variances and the decoder reconstructed the complete typed feature matrix. The objective combined mean-squared reconstruction error, scaled by the number of input features, with a beta-weighted Kullback–Leibler (KL) term. Beta was linearly annealed from zero to 1 during the first quarter of training and then held at 1.

Models used Adam^77^ with learning rate 10^−3^, weight decay 10^−5^, default momentum parameters and global gradient-norm clipping at 1.0. A ReduceLROnPlateau scheduler used patience 20 and factor 0.5. Real-data fits used hidden dimension 256, latent dimension 32 and at most 500 epochs. Twenty percent of samples were assigned to validation using the configured random seed. Inputs with more than 2,000 features used automatic minibatching with batch size min(64, ⌈*n*/4⌉); smaller inputs used full-batch training. The checkpoint with the lowest post-warm-up validation loss was retained. Training stopped after 50 epochs without an improvement greater than 10^−4^, and non-finite validation loss triggered restoration of the best checkpoint. NumPy and PyTorch seeds and the selected CPU or CUDA device were recorded. Latent dimensions with mean KL divergence below 0.01 were flagged as collapsed; fits with at least half their latent dimensions collapsed were flagged for review.

Gene similarity was calculated by double-centering decoded feature reconstructions across samples and features, *L*_2_-normalizing each feature and taking their dot product, equivalent to Pearson similarity after centering. For each gene, mean non-dominant transcript usage was 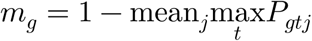. Its switch reliability was min(1, *m*_*g*_/0.10). Switch–switch edge weights were multiplied by the geometric mean of the two gene reliabilities before thresholding; abundance edges were not downweighted. This estimability correction used neither phenotype nor sample covariates. Candidate feature pairs were screened by absolute similarity and projected to one edge per gene pair by retaining the strongest typed association. Community detection retained only edges whose signed weight was positive; the retained weights were not replaced by absolute correlations. Thus a negative association could pass absolute-similarity screening but was excluded from community detection. Because a switch–switch similarity changes sign when either gene’s axis is flipped, this step depends on the orientation convention described above: two genes whose oriented coordinates are negatively correlated are not connected, even though their transcript usage covaries. Module membership is therefore reproducible under the fixed rule but reflects the implementation rather than an intrinsic direction of switching. The resulting graph was partitioned by seeded, weighted Leiden community detection^24^ at canonical resolution 2.0, and communities with fewer than 20 genes were discarded.

Resolution 5.0 and a phenotype-blind resolution sweep (**Data S6**) were retained as sensitivity analyses. Resolution was never selected from trait-association results. Each fit saved modules, edges, traits, feature scores, reconstructed features, module gene roles, calibration values and configuration metadata.

The module score used in every module-level age and diagnosis test was the arithmetic mean of the saved feature rows of member genes (`eigengene_table` in IsoGraph v0.1.5). In a multiplex fit this mean includes both switch and abundance rows, including those of abundance-only genes, and no iterative sign alignment is applied, so it is a combined module score rather than a switch-only eigengene. Analyses that separate the two channels (incremental models and channel roles) computed switch-only and abundance-only scores explicitly, as described below.

Three distinct summaries are used for different purposes. The production module score `eigengene_table`) is the arithmetic mean of the saved member-feature rows, including both channels in multiplex fits, and is used in the module-level age and diagnosis tests. The projection eigengene is the first principal component of source-standardized member-feature rows with frozen weights and is used for cross-cohort projection. The channel-specific reference is the first component of the switch rows or of the abundance rows taken separately and is used for gene channel-role assignment.

Channel roles were calculated separately. Within each module, the first component of all switch features and the first component of all abundance features defined channel-specific references. A gene’s switch and abundance scores were correlated with the corresponding references and called active at |*r*| ≥ 0.2. Genes with only an active switch channel were `switch_only`, those with only active abundance were `abundance_only`, and those with neither were `inactive`. Genes with both channels active were `coupled`when their within-gene switch–abundance correlation was at least 0.2 `discordant` and otherwise.

### Comparator networks

WGCNA^11^ was used in three comparator spaces: gene abundance, switch-only features, and the same multiplex switch-plus-abundance features supplied to IsoGraph. All comparator fits used the same production gene universe. Matched-feature comparisons therefore varied network inference while holding the input representation fixed; abundance WGCNA represented a conventional gene-expression analysis. WGCNA first assigned typed features to modules. Feature identifiers were then mapped back to genes and duplicate gene–module rows removed. If a gene’s two channels belonged to different modules, both memberships were retained; an abundance-only gene inherited its abundance feature’s module. The resulting multiplex comparator is therefore an overlapping collection of gene sets rather than a single gene partition, and partition-based comparisons with it require an explicit overlap-handling rule. WGCNA used signed topology, a soft-thresholding power selected at scale-free topology fit *R*^2^ ≥ 0.85, minimum module size 20 in real-data comparisons and merge cut height 0.25. Reported comparisons used per-module rates where raw counts would be determined by partition granularity (**Fig. S15**; **Table S4**; **Data S4**).

### Synthetic benchmarks

Synthetic data tested recovery under known ground truth. Core datasets contained 400 genes and 160 samples; scale experiments extended to 16,000 genes and 300 samples. Samples were balanced between control and schizophrenia labels. Ages spanned 25–85 years with Gaussian perturbation, and sex labels alternated. Switching genes were assigned approximately evenly among planted modules, and each module received a latent variable with independently sampled age and diagnosis effects. Scenario grids varied switch-signal noise, count dispersion, and feature interactions (**Fig. S3**), non-switching background (**Fig. S4**), isoform imbalance (**Fig. S5**), abundance-driven structure (**Fig. S2**), batch, cell composition, library depth, and RNA degradation (**Fig. S6**; **Fig. S9**), null structure (**Fig. S8**), and dataset scale (**Fig. S10**); all scenario parameters are listed in **Table S9**.

For two-transcript simulations, transcript-1 usage followed a logistic model of the planted switch signal and an isoform-imbalance parameter. Gene means were sampled from a log-normal distribution, sample depth from a second log-normal distribution and total counts from a gamma– Poisson negative-binomial model. Transcript-1 counts were binomially sampled conditional on the total, and transcript-2 counts were the remainder. Dataset identifiers, parameters and random seeds were recorded.

A genetic-anchoring scenario assigned one biallelic *cis*-variant to each planted module. Genotypes followed Hardy–Weinberg equilibrium at minor-allele frequency 0.30 and shifted the corresponding module latent variable additively. An equal number of unlinked variants served as null controls. The per-allele effect was varied over 0.5, 1.0 and 2.0 and switch noise over 0.25 and 0.40 across 20 seeds. Each predicted-module eigengene was regressed on genotype, and the best *R*^2^ and Bonferroni-corrected anchor call were recorded for each planted variant.

Module recovery was the mean, over true modules, of the maximum Jaccard similarity to any predicted module. Additional endpoints included adjusted Rand and mutual-information measures (**Fig. S7**), switch-gene recall, background-gene assignment, driver-transcript ranking (**Fig. S1**), wall-clock time and maximum memory (**Table S1**; **Table S2**). Means and 95% confidence intervals were obtained from 10,000 non-parametric bootstrap resamples. Methods were compared on shared simulated datasets using paired Wilcoxon signed-rank tests; unmatched rank-sum tests were used only for diagnostic summaries lacking paired runs. Benjamini–Hochberg correction was applied across the complete prespecified scenario-by-metric-by-method family.

### Trait association and incremental models

Each fit saved an unadjusted linear-age analysis using Pearson correlation and a separate covariate-adjusted age-spline analysis. The covariate-adjusted spline was used to select age-associated modules and genes. The covariate-free linear correlation was used only for the reproducibility analyses (split-half sign agreement, cross-cohort concordance and projection), where it gives a signed effect comparable across cohorts. Repeating the gene-membership-matched cross-cohort concordance with the spline sensitivity model did not change the conclusion: under the matching null and `complement` covariate mode, IsoGraph had 0 of 38 concordant pairs (*P* = 1) and WGCNA had 5 of 52 (*P* = 0.161). This sensitivity analysis used a three-degree-of-freedom natural-spline block test, with direction supplied by the corresponding linear age coefficient. The `complement` mode added covariates outside the discovery residualization set; `full` and `none` modes were also reported (**Data S8**). For primary trait selection, the spline analysis fit the module score as a function of a natural cubic spline of age, with one internal knot at the median and boundaries at the observed minimum and maximum, plus the cohort-specific covariates above. A partial *F*-test compared this model with the covariate-only model. The diagnosis analysis fit the linear model in Eq. 3,

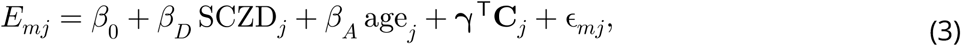

where *E*_*mj*_ is the saved module score, SCZD_*j*_ indicates a schizophrenia diagnosis, and *C*_*j*_ contains sex, manner of death, RIN, mapping rate, mitochondrial mapping rate, and SNP PCs 1–5. The SCZD coefficient, relative to controls, was tested by a two-sided ordinary-least-squares-test using complete cases. Benjamini–Hochberg adjustment was performed across modules within each trait and model. The unadjusted Welch test that IsoGraph also writes was not used for diagnosis inference.

For genes with both feature channels, separability was summarized by the Pearson correlation between raw switch and standardized abundance scores (**Table S3**; **Fig. S12**). Incremental models included at least 30 complete samples. In the schizophrenia cohort, switch scores were regressed on diagnosis, the same gene’s abundance score, age, sex, manner of death, RIN, mapping rate, mitochondrial mapping rate, and SNP PCs 1–5; the reverse model exchanged switch and abundance outcomes. Diagnosis terms were assessed by two-sided ordinary-least-squares *t*-tests. For aging, age was represented by the natural spline described above and assessed by a partial *F*-test comparing full and reduced models. BrainSEQ models used the BrainSEQ covariates above, and GTEx models used sex, RIN, ischemic time, and mapping rate. Correction was applied separately to switch-primary and abundance-primary tests within an analysis. FDR below 0.10 defined switch-unique, abundance-unique and jointly associated genes for this characterization; other thresholds are shown in **Fig. S13**.

Cell-type proportions were estimated with MuSiC^23^ using prespecified Tran/LIBD single-nucleus RNA-sequencing references^78^. GTEx deconvolution was restricted to eight regions with a defensibly matched reference; direct anatomical matches were used where available, striatal regions were mapped to the nucleus accumbens reference and cortical regions to the DLPFC reference. Incremental models were repeated with estimated cell-type fractions added as covariates. Results with and without composition adjustment were retained; no module or gene was selected using the composition-adjusted result before the unadjusted analysis was run (**Data S2**).

To judge the risk of over-adjustment, each estimated cell-type proportion was correlated with donor age by Spearman correlation, with Benjamini–Hochberg correction across cell types within each analysis. As a control independent of the covariate model, module genes were tested for enrichment and depletion of each cell type’s 25 top-ranked mean-ratio markers from the matched reference, by one-sided hypergeometric tests against the analysis’s module-gene background and with Benjamini– Hochberg correction across module-by-cell-type tests within each analysis (**Fig. S14**).

### Module annotation and GO enrichment

GO Biological Process enrichment^79,80^ was performed with `goatools`^81^ against the analysis-specific tested-gene universe. Modules with fewer than five mapped genes were not tested, and Benjamini–Hochberg adjustment was applied to terms within the specified analysis family. A phenotype-associated module (age or diagnosis FDR below ≤0.05) was termed GO-invisible when it had no enriched Biological Process term at FDR and GO-visible otherwise. The number and strength of anticorrelated transcript pairs and their structural consequences were calculated independently of GO status.

### Module stability and replication

Within each of six anatomically paired BrainSEQ and GTEx aging analyses, we performed five random split-half refits. For every production module, module-restricted co-assignment density was the proportion of within-module gene pairs assigned together across split-half fits. This statistic was compared with size-matched random gene sets, and modules with FDR below 0.05 were called trusted (**Data S3**; **Data S5**). Driver reproducibility was the Spearman correlation of per-gene switch-axis importance, the largest absolute transcript loading of each gene, between each trusted production module and its best-matching split-half module over shared genes; absolute loadings were used because the axis sign is fixed independently in every fit.

Trusted BrainSEQ modules were matched to GTEx modules from the corresponding brain region using gene and top-driver overlap. Replication required directionally concordant age associations in both cohorts and was evaluated against a matched permutation null (**Fig. S16**; **Data S8**). Raw best-match Jaccard similarity was not used as a trust criterion because it increases with module size.

For cross-cohort projection, the frozen eigengene of each trusted source module was defined as the first principal component of its member feature rows (switch and abundance rows for IsoGraph, abundance rows for the WGCNA comparator), each row standardized across source samples, with the unit-norm weight vector oriented so that the eigengene correlated positively with the mean of the member features. This projection eigengene is distinct from the production module score used in the trait tests. The weight vector was applied unchanged to the same features in the target cohort, and the projected eigengene was standardized to unit variance. Switch axes were aligned before projection using shared-transcript loadings: the cosine between a gene’s source and target loading vectors set the sign of its target row, and genes with fewer than two shared transcripts or an absolute cosine below 0.5 were excluded from switch rows. IsoGraph projections used switch and abundance rows, whereas the projection comparator was classical abundance-only WGCNA rather than the same-feature WGCNA comparator used elsewhere in the study. Preservation was quantified by the mean signed feature-to-eigengene correlation, that is, the mean over member features of each target feature’s correlation with the projected eigengene multiplied by the sign of its frozen weight. It was compared with 1,000 random projections (seed 13) that applied the same weight vector to randomly drawn target features matched in number and feature type, giving a one-sided empirical *P* value with Benjamini–Hochberg correction across modules. For the age comparison, each cohort’s frozen-eigengene age correlation was standardized against the age correlations of the same-weight random projections in that cohort, *z* = (*r* − *μ*_null_)/*σ*_null_, and concordance was assessed between these standardized statistics. Raw age-sign concordance was retained separately (**Data S7**; **Data S27**). No covariates entered the projection-age test; null standardization does not make this a covariate-adjusted analysis.

Because matched modules shared few genes, functional preservation was assessed at three further levels. For each matched BrainSEQ–GTEx pair we calculated the Jaccard index of the two modules’ enriched GO Biological Process term sets, the Pearson correlation of their cell-type marker-enrichment profiles (log_2_ of observed over expected marker genes, each offset by 0.5) and the Pearson correlation of their switch coding-consequence class rates; profile correlations required at least three shared cell types or classes. All three measures increase with module size, so the null re-paired each BrainSEQ module with a random GTEx module from the gene-count decile of its match (1,000 permutations, seed 13). The observed mean over pairs was compared with the null distribution of means by a one-sided empirical *P* value, for the linear and spline age models separately. Cell-type profiles were available only for IsoGraph modules, and consequence profiles only in regions with phenotype-significant switch genes; measures without data were reported as untested (**Data S9**).

### Gene-wise DTU concordance and junction-usage corroboration

Genes of phenotype-associated IsoGraph switch modules were compared with satuRn DTU results^8^ fitted to the same transcript-count matrix and trait design. A gene’s continuous DTU evidence was of its −log_10_ minimum transcript-level *P* value. Evidence in switch-module genes was compared with that in other tested genes by a one-sided Mann–Whitney test and rank-biserial effect size; analyses without switch-module genes or without satuRn results were not testable. A logistic model additionally adjusted for the number of tested transcripts per gene (**Fig. S17**; **Data S10**).

Junction-level corroboration used splicing summaries derived from split reads of the same alignments rather than from the transcript quantifier: PSI events in BrainSEQ and within-gene junction usage from STAR junction counts in GTEx. Each event was tested for association with the trait under the covariate model of the corresponding analysis. Per gene, the minimum event *P* value was Bonferroni-adjusted for the number of events and then Benjamini–Hochberg-adjusted across genes; a gene was PSI-positive at FDR ≤0.05. Switch-positive genes were members of a trait-associated IsoGraph module (module set) or, in BrainSEQ caudate, genes whose own switch coordinate was trait-associated at FDR ≤0.05 (marginal set). Enrichment of PSI-positive genes among switch-positive genes was tested by a one-sided Fisher exact test and by a logistic model of PSI status on switch status adjusted for log transcript count. In BrainSEQ, junctions implicated by colocalization events that resolved to a switch pair were additionally matched to PSI events by coordinate overlap, and the event’s trait association was compared with the gene’s switch-coordinate association (**Data S11**).

### Held-out module-context prediction of DTU

For each of six matched cohort-region analyses, satuRn^8^ was fit separately in both halves of five sample splits. Both discovery-to-held-out directions were evaluated, using only the discovery-half partition to define neighbors. Genes had to be tested in both halves and belong to a discovery module with at least five eligible genes. For each gene, evidence was *e* = −log_10_(*P*_Simes_), where Simes’ method^82^ combined its tested transcript-level *P* values. The outcome was held-out evidence and the context predictor was the leave-one-out mean of discovery evidence among eligible module members.

The adjustment model contained an intercept, a centered natural cubic spline of the focal gene’s discovery evidence (four degrees of freedom), log tested transcript count, log-one-plus mean gene count and minor-isoform usage. Mean gene count was the mean per-sample sum of tested transcript counts in the discovery half. Minor usage was one minus the largest transcript’s mean within-gene usage, excluding zero-total samples when averaging usage. The statistic was the partial correlation of context and held-out evidence after residualizing both against this adjustment model.

The region-level statistic was the mean partial correlation across the ten split directions; medians, ranges, and the number of positive directions were also retained. For each of 1,000 permutations, discovery module labels were shuffled among genes within discovery-expression deciles crossed with minor-usage tertiles. This preserved module sizes and stratum composition; DTU evidence and adjustment variables stayed attached to their genes. Leave-one-out context was recomputed after every shuffle. Each direction was permuted independently and the ten-direction mean formed one null statistic. One-sided empirical *P* values used (1 + #{*T*_*null*_ ≥ *T*_*obs*_})/1001, with Benjamini–Hochberg correction across the six regions. Split directions were not treated as independent studies in a meta-analysis (**Fig. S18**; **Data S12**).

The abundance comparator averaged discovery evidence among the 50 genes with the highest signed correlations in covariate-residualized discovery log-CPM, restricting neighbors to genes with available evidence. The focal gene was excluded from both module and abundance neighborhoods. In addition to testing the abundance context alone, the joint model tested module context conditional on abundance context using the same stratified module-label null. Missing abundance context was replaced by the mean focal-gene discovery evidence in the analysis frame.

For descriptive prioritization, discovery gene-level Simes *P* values were Benjamini–Hochberg-adjusted over genes tested in both halves. Among eligible genes with discovery *q* ≥ 0.05, module-context ranks were divided into tertiles within each split direction, breaking ties by row order. We compared the fraction with held-out nominal Simes *P* < 0.05 in the top and bottom tertiles. A logistic model, excluding the middle tertile, estimated the top-versus-bottom odds ratio conditional on the same four-degree-of-freedom spline of discovery evidence, log transcript count, log-one-plus mean count and minor-isoform usage. This descriptive model did not additionally condition on co-expression context. Directions with fewer than 30 sub-threshold genes were not evaluated, and failed logistic fits contributed no odds ratio. Rates were averaged and fitted odds ratios summarized by their median across the ten directions. Here, held-out support denotes nominal evidence, not an FDR-significant replication call. Post hoc sensitivities recomputed the continuous statistic after excluding modules of at least 900 genes or each split’s largest module.

The analysis code specified a study-level decision rule for the model before co-expression adjustment: *q* < 0.05 in at least four of six analyses, including at least one from each cohort. Four analyses met that rule. The additional co-expression-adjusted model retained three significant analyses and is reported separately; the four-of-six criterion was not met after that adjustment.

### Long-read transcript support and short-read junction corroboration

Long-read transcript support was assessed using published Oxford Nanopore cDNA sequencing from aged human DLPFC (BA9/46; 12 donors, six with Alzheimer’s disease and six controls)^18^. Bambu transcript counts^37^ were obtained from Zenodo^83^; raw reads were not reprocessed. A transcript was detected at five or more counts in at least three samples, and a pair was detected when both transcripts were. A detected pair was switch-like when isoform fractions were negatively rank-correlated in at least four jointly measured samples. Anchored events were the colocalization events whose junction was carried by a tissue-matched switch-pair isoform (55 genes); the overlap-score events (30 genes) were scored separately as a sensitivity analysis. For the genetically anchored pairs, each anchored isoform was paired with every other transcript in its gene’s tissue-matched switch pairs, and repeat observations across events, traits, and tissues were collapsed to one row per anchored–partner orientation. Because detection, the usage correlation and the abundance stratum are symmetric in the two transcripts, orientations were then collapsed to distinct unordered transcript pairs, the primary unit of analysis; a pair whose two members both carried the colocalizing junction contributes one distinct pair and two orientations, and the orientation-level test was retained as a sensitivity analysis. A distinct pair was well expressed when an anchored isoform had a mean within-gene isoform fraction of at least 0.05 across the 12 samples. Because within-gene fractions are compositional, negative correlation was compared with a null matched on the abundance decile of the better-expressed transcript. For all IsoGraph switch pairs genome-wide, the null was detected non-switch transcript pairs from the same genes, scored in the same samples; for the anchored pairs, it was IsoGraph switch pairs from genes outside the prioritized set, matched on abundance but not on detection. Thus, the prioritized-pair comparison controls abundance but does not ensure equal transcript-detection rates in the observed and background sets. Two thousand matched draws were performed with seed 13. This analysis tested isoform existence and switch-like usage, not age association in the 12-donor cohort (**Data S25**).

Short-read junction corroboration (**Fig. 6c**; **Data S26**) used the same 55 signal-colocalization-prioritized genes as the long-read analysis. For each nomination, we retrieved the tissue-specific IsoGraph switch pairs containing a transcript that carried the implicated junction. Transcript pairs were matched to the existing BrainSEQ junction recount models by exact, versioned transcript identifiers, allowing reversal of pair order. Primary anatomical mappings were GTEx BA9 to BrainSEQ DLPFC, hippocampus to hippocampus, and caudate to caudate; generic cortex to DLPFC was included as an anatomical proxy. Eligibility required discriminating junctions for both alternatives and the implicated junction to be present in the pair model, with exact chromosome, strand and intron endpoints after converting exon-flanking coordinates to closed intron coordinates (start + 1, end − 1). Repeated nominations were collapsed to one unordered transcript pair per BrainSEQ region, prioritizing the closest anatomical match.

We reused the fragment counts described under Allele-specific junction analysis, restricting donors to the adult control aging cohorts with completed recounts. Counts were summed across both haplotypes and fragments without haplotype assignment, without filtering on heterozygosity, allelic significance or allelic-test eligibility. Missing recounts were excluded rather than assigned zero counts. For each pair, covered donors had at least ten total pair-discriminating fragments, and a pair was measurable when at least 30 donors met this threshold. The primary endpoint was the fraction of covered donors with at least one fragment supporting each transcript alternative. The panel shows individual pair fractions, gene–region medians and observed pair ranges; ranges are descriptive, not confidence intervals. Nominations without a matched region, an eligible recount model or sufficient coverage were retained in the coverage summary rather than classified as unsupported.

Sensitivity analyses used coverage thresholds of five and 20 fragments, required at least two fragments per alternative, excluded the generic-cortex proxy, or restricted pairs to those for which the implicated junction itself discriminated the alternatives. Adjacent mappings (BA24 to DLPFC and putamen or nucleus accumbens to caudate) and secondary cortical mappings to caudate were evaluated separately. Junction models distinguish the two members of a pair but need not distinguish either member from every other transcript of the gene. Because counts aggregate across model junctions, support does not establish direct observation of the particular colocalizing junction or a full-length transcript. These measurements reuse BrainSEQ reads and provide complementary measurement corroboration, not independent phenotype replication.

The separate supplementary junction-usage analysis of overlap-score-anchored switch pairs used the BrainSEQ PSI event catalog in the anatomically matched region (DLPFC for cortical GTEx tissues, with caudate carried alongside). For each candidate event containing the anchored junction, usage was summarized as (*median PSI*, 1 – *median PSI*), which is invariant to undocumented PSI orientation. An event had junction-usage support when this minor-form usage was at least 0.05, establishing detectable alternative usage rather than replication of a phenotype association. Genes nominated only in GTEx tissues without a matched BrainSEQ region were retained as not assessable rather than removed (**Data S28**).

### Structural consequences of switching

Analyses were annotated when they contained at least one phenotype-significant switch gene (FDR < 0.05); nine aging analyses and the schizophrenia analysis met this criterion, and aging and diagnosis analyses were pooled separately. Observed switch-pair transcripts were compared with GENCODE v47 models for changes in first exon, last exon, internal exons, UTRs, coding sequence, transcript biotype, coding status, and transcript length. A coding consequence was defined by a coding-sequence or coding-status difference. Nonsense-mediated decay status was assigned using the strand-aware 50-nucleotide rule^84,85^. To control for transcript number and gene architecture, the observed number of pairs for each gene was sampled with replacement from all annotated transcript pairs of that gene. Consequence rates were recomputed over 1,000 permutations with seed 13. One-sided empirical *P* values were calculated as (1 + #{*T*_null_ ≥ *T*_obs_})/1001. Region-level values were combined by Fisher’s method, and enrichment ratios (observed over permuted consequence rate) were also pooled across annotatable analyses by DerSimonian–Laird random-effects meta-analysis^86^ of the log ratio, with *I*^2^ reported. For the driver-level description, the top-ranked driver transcript pair of each age-associated trusted module was annotated in the same way, and the fraction of modules whose driver pair differed in each structural class was reported with its standard error across modules; these fractions carry no null.

Gene-level constraint was obtained from gnomAD v4.1^25^. When several transcripts represented a gene, the transcript with the lowest LOEUF was retained. LOEUF among switch genes was compared with the genome-wide reference by a one-sided Mann–Whitney test in each annotatable analysis, combined by Fisher’s method. For ClinVar^26^, exons found in exactly one member of an observed switch pair were compared with other exons from the same switching transcripts. Pathogenic and likely-pathogenic ClinVar VCF entries were counted per kilobase by genomic position, one count per VCF data line without deduplication by chromosome, position, reference allele, and alternate allele; the densities are therefore VCF-entry densities rather than unique-allele densities. Switched/ background labels were permuted within gene 1,000 times (seed 13), preserving the number of switched exons, and a two-sided empirical *P* value tested either enrichment or depletion; analysis-level *P* values were combined by Fisher’s method (**Fig. S19**; **Table S5**).

### BrainSEQ switch-QTL and abundance-QTL mapping

BrainSEQ *cis* switch-QTL and abundance-QTL mapping used tensorQTL^87^ with a 1-Mb window, minor-allele frequency threshold 0.01 and 10,000 permutations. Genotype-matched phenotype cohorts contained 420 caudate, 362 hippocampus and 360 DLPFC samples. Within each phenotype, values were rank-based inverse-normal transformed as *Φ*^−1^(*r*/(*n* + 1)), using average ranks for ties. Known covariates were SNP PCs 1–5, sex, diagnosis, RIN, mitochondrial mapping rate, and mapping rate. Thirty hidden factors were estimated separately for switch and abundance phenotypes using the same procedure and factor count. Transformed phenotypes were regressed on an intercept and known covariates; residual columns were centered and standardized, constant columns removed, and singular value decomposition applied to the sample-by-phenotype matrix. The first 30 sample PC scores (*UD*), standardized to unit variance, were added to the known covariates in tensorQTL. The mapping outcome remained the inverse-normal phenotype, with covariates entered jointly. Empirical permutation *P* values were Benjamini–Hochberg-adjusted within modality. Because the switch phenotype was recomputed on the genotype-matched samples and re-oriented by the same rule, each gene’s recomputed coordinate was correlated with its discovery coordinate over the shared libraries (median |*r*| ≥ 0.99 required across genes), and switch-QTL slopes were multiplied by the sign of that correlation so that effects are reported on the discovery orientation; genes with |*r*| < 0.5 were flagged as axis-unstable and carried no directional comparison.

### Allele-specific junction analysis

For BrainSEQ genes with switch-QTL evidence, fragments crossing an isoform-discriminating junction and carrying a phased heterozygous site were recounted from the same WASP-tagged STAR alignments used to generate the cohort-wide ASE release. GENCODE v47 exon models were converted to transcript introns, and junctions present in one but not the other member of a switch pair were designated isoform-discriminating. Alignments were grouped by read name so both mates were evaluated as one fragment. Every alignment had to be primary, mapped, not flagged as failing quality control, and uniquely mapped `NH:i:1`). No mate could carry a failing WASP tag, and every mate covering a heterozygous site had to carry `vW:i:1`.

A splice junction was counted only when its CIGAR `N` operation had at least eight aligned bases on each side. The fragment had to cross a junction specific to one isoform, no junction specific to the other isoform, and no junction within the pair’s genomic span that was incompatible with the assigned transcript. Haplotype was assigned from bases of quality at least 10 at phased heterozygous single-nucleotide variants (SNVs) using phASER `PW`. Overlapping mates had to agree at a site, and all informative sites carried by a fragment had to indicate the same haplotype; conflicts were excluded and retained as a QC rate. Fragments lacking a heterozygous site contributed to isoform totals but not allelic contrasts. Per-site counts from the recount were compared with the corresponding ASEReadCounter output as a concordance check.

The switch-QTL gate family comprised transcript pairs in genes with an all-samples BrainSEQ switch-QTL at *q* < 0.05. A pair passed the prespecified feasibility gate when at least 30 donors had at least five informative fragments with both isoforms represented. The allelic analysis proceeded only when at least 30 switch-QTL gate-family pairs passed in the region. Individual fitted pairs required at least ten lead-heterozygous donors informative on both haplotypes (**Data S15**).

For donor *d*, haplotype-specific transcript-1 counts *y* out of total pair counts *n* were modeled with the beta-binomial mixed model in Eq. 4,

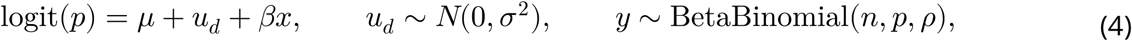

where *x* marked the haplotype carrying the switch-QTL alternate (ALT) allele. A likelihood-ratio test compared the full model of Eq. 4 with *β* = 0. Donors homozygous at the lead were fit in a separate null-control arm, using arbitrary haplotype labels and other phased heterozygous sites to assign fragments. They did not contribute an ALT-versus-reference (REF) contrast at the lead or enter the primary heterozygote effect estimate. Their separate test assessed haplotype imbalance without a lead-allele difference (**Fig. S21**); the between-donor dosage sensitivity analysis could additionally use their isoform totals. A model-free within-donor score test and between-donor dosage correlation were reported as sensitivity analyses. Fitted effects at the optimizer bound were treated as quasi-separated: their direction, but not magnitude, was retained. To orient effects to disease, the GWAS risk allele at the colocalizing locus was transferred to the fitted QTL lead only when signed in-sample LD satisfied |*r*| ≥ 0.8; effects below this gate remained unoriented. A second analysis anchored directly on the GWAS lead.

To ask whether *cis* control tracked module age association, the fitted allelic test was aggregated to IsoGraph modules without refitting. A gene was *cis*-controlled when any of its fitted gate-family pairs had *q* < 0.05. Modules were selected on linear or spline age models whose effects are not on a common scale, so absolute module age effects were ranked within each region and age trait and scaled from 0 (weakest) to 1 (strongest); traits with a single module were dropped. Within a region, the Spearman correlation between each module’s *cis*-controlled-gene fraction and its age rank required at least three ranked modules. Its null permuted gene-to-module labels 10,000 times, preserving module sizes and the number of *cis*-controlled genes. Regional correlations were averaged into a pooled statistic with a two-sided empirical *P* value. A sensitivity analysis excluded modules with fewer than five fitted genes (**Table S8**).

### Colocalization and event resolution

Selected switch genes were the member genes of phenotype-associated IsoGraph modules (module– trait FDR ≤ 0.05), taken irrespective of channel role. For the aging-anchored analyses of Alzheimer’s disease, Parkinson’s disease, dementia with Lewy bodies, amyotrophic lateral sclerosis and schizophrenia, a gene was selected when it belonged to an age-associated module in at least one of the 16 aging analyses (three BrainSEQ regions and 13 GTEx tissues); for schizophrenia, the genes of diagnosis-associated modules in the BrainSEQ caudate analysis were analyzed in addition. The same gene sets defined the stratified LD-score annotations below. Candidate disease loci contained a GWAS variant with *P* < 10^−5^ within 1 Mb of a selected switch gene; overlapping windows were merged. GWAS variants were allele-aligned to 1000 Genomes European reference LD. Strand-ambiguous and allele-mismatched variants were removed, as were the major histocompatibility complex for all traits and the *APOE* interval for Alzheimer’s disease and dementia with Lewy bodies. GWAS signals were fine-mapped with `susie_rss` using at most ten effects, fixed residual variance and locus effective sample size floored at 1,000^88,89^. Loci with fewer than 20 or more than 12,000 usable variants were not fine-mapped in the primary analysis; an Alzheimer’s disease sensitivity analysis raised the upper limit to 30,000.

GTEx expression and LeafCutter intron-excision^90^ phenotypes were re-fine-mapped from v11 nominal *cis*-association statistics on the same variant and LD universe. A QTL credible set was retained in the primary signal-level analysis only if it shared a variant with the corresponding GTEx in-sample SuSiE credible set. Every GWAS–QTL signal pair sharing at least 100 variants was tested with `coloc.susie`^21^. Default trait priors *p*_1_ = *p*_2_ = 10^−4^ were used, and *p*_12_ was swept over 10^−6^, 5 × 10^−6^, 5 × 10^−5^, 10^−4^, with 10^−6^ prespecified as primary.

Cells not estimable by signal-level colocalization were analyzed with `coloc.abf` on the complete shared *cis*-window^20^. Each cell recorded its estimator and first fallback reason. A separate credible-set posterior-overlap analysis combined GWAS SuSiE posterior inclusion probabilities (PIPs) with the GTEx QTL credible-set PIPs. For shared variants *v* in a QTL credible set *C*, the credible-set posterior-overlap score was defined as

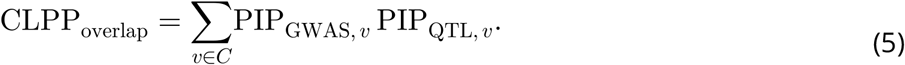

Eq. 5 follows the posterior-product principle used in eCAVIAR^22^, but uses SuSiE outputs and sums across variants; it is not a run of the eCAVIAR executable and is not a posterior probability. The main text calls this quantity the credible-set posterior-overlap score and writes it CLPP_overlap_; the supplementary tables and the analysis code label the same quantity CLPP. The highest contributing variant and its two PIPs were also saved. The screen used CLPP_overlap_ ≥ 0.01 and separately reported ≥ 0.05, summarizing the largest score within each gene and QTL class. These are screening thresholds for the implemented score, not posterior probabilities of disease mediation. Colocalization and the overlap score were presented jointly for disease interpretation, with their estimators, eligibility sets, and disagreements explicit. The overlap score did not determine the separate signal-level nomination list. Signal-level nominations required sQTL PP4 ≥ 0.8 `coloc.susie` from in at least one brain tissue at the primary prior. Cells not estimable at signal level with `coloc.abf` PP4 ≥ 0.8 were labeled fallback-supported loci and tabulated separately. A fallback posterior did not promote a locus to a signal-level nomination; maxima were summarized within estimator tier. Of the 119 gene-by-trait nominations, 33 had `coloc.susie` support in at least one tissue and 86 rested on the fallback only; both tiers are reported, labeled by estimator (**Data S16**). Paired sQTL/eQTL calls were compared without selection using exact McNemar tests and paired Wilcoxon tests of the conditional posterior PP4/(PP3 + PP4), where PP3 is the posterior probability of distinct causal variants (**Data S17**).

Event resolution was run on both layers: on the intron behind each tissue’s colocalization call (`coloc.susie`, or GTEx’s representative intron for `coloc.abf` cells) and on the overlap-score events. Colocalization was primary, and the overlap score corroborated it. Colocalizing intron identifiers were converted to genomic junctions and matched within 2 bp to introns derived from adjacent GENCODE v47 exons. Junctions were then mapped to the tissue-matched IsoGraph switch pair. An event was concordant when a transcript carrying the colocalizing junction was a member of that switch pair. Cross-cohort pair mapping was recorded when a junction transcript was also a switch-pair isoform of the same gene in the anatomically matched BrainSEQ region. This records transcript-pair overlap across independent cohorts, not replication of the event’s phenotype association (**Data S19** and **S21**). Curated literature events were verified independently against GENCODE coordinates before comparison. Nominations were classified as known mechanism recovered, context-distinct splice colocalization, disease-locus splice-linked, novel splice-led candidate or unresolved according to whether the curated event was testable and whether the colocalizing intron matched it. For loci carrying a biological claim, robustness checks assessed credible-set convergence, size and purity, boundary trimming, the *p*_12_ sweep, `estimate_s_rss`, `kriging_rss` outlier removal and residual-variance estimation.

BrainSEQ switch- and abundance-axis QTLs from the European-ancestry mapping were colocalized with the same GWAS fine-mapping using the same estimator hierarchy. QTL signals were fine-mapped with `susie_rss` on in-sample LD, computed as correlations of covariate-residualized ALT dosages in the mapped donors. Axes were compared per gene with exact McNemar tests, each gene contributing its best region on both axes. Because BrainSEQ is the discovery cohort, this was treated as same-cohort anchoring, not replication (**Fig. S22a**; **Data S18**).

SMR and HEIDI analyses were run with SMR v1.4.2 as corroboration beneath colocalization^34^. GTEx v11 ALT-allele QTL effects were harmonized to the same GWAS and 1000 Genomes European panel. Instruments required QTL *P* < 5 × 10^−8^; HEIDI used 3–20 variants at QTL *P* < 1.5654 × 10^−3^, a 2-Mb *cis*-window and allele-frequency difference below 0.2. SMR significance was Bonferroni-corrected over instrumented probes within trait and QTL class, and HEIDI *P* < 0.01 rejected a single shared variant. These results did not alter colocalization calls and were not interpreted as causal effects (**Fig. S22b**; **Data S20**).

Module-level convergence tested whether posterior-overlap-supported genes concentrated in IsoGraph modules of the BrainSEQ and GTEx caudate aging partitions, separately for the five colocalization traits. Genes were mapped to each partition’s own module assignments, and the tested pool comprised genes that received an overlap-score test for the trait and belonged to the partition. Concentration was the sum of squared per-module counts of overlap-supported genes. Its null redrew the same number of genes from the tested pool 20,000 times (seed 13), holding module sizes fixed, and gave an empirical *P* value. Per-module over-representation was tested hypergeometrically against the tested pool for modules with at least three pool genes, with Benjamini–Hochberg correction within trait and partition, and a leave-one-locus-out sensitivity removed each module’s largest contributing GWAS locus. Enrichment of overlap-supported genes in MAGMA-anchored^91^ modules (competitive gene-set *P* < 0.05; see below) was tested hypergeometrically against both the tested pool and all module genes. The tested pool was the primary denominator because MAGMA anchoring and overlap-score testing select on the same GWAS signal (**Fig. S24**; **Data S23**).

### QTL enrichment and module gene-set tests

For each cohort-region analysis, logistic regression tested whether IsoGraph membership predicted GTEx v11 *cis*-sQTL or *cis*-eQTL status among genes tested for the corresponding QTL class. Models adjusted for *cis*-variant count, gene length, retained isoform count and intron-group size where available. The primary estimand was the within-analysis ratio of the sQTL odds ratio to the eQTL odds ratio, which removed the shared *cis*-QTL depletion of network genes. Log odds ratios were pooled by inverse-variance fixed- and DerSimonian–Laird random-effects models^86^, with *I*^2^ reported. Identical analyses were applied to matched WGCNA switch-only and multiplex modules (**Fig. S23**; **Tables S6** and **S7**).

Competitive gene-set analyses of whole modules used MAGMA v1.10^91^, with the 1000 Genomes European panel for LD and European-ancestry GWAS summary statistics for schizophrenia^33^, major depressive disorder^92^, bipolar disorder^93^, Alzheimer’s disease^29^, Parkinson’s disease^30^, dementia with Lewy bodies^31^, amyotrophic lateral sclerosis^32^ and stroke^94^. Gene-level statistics were computed once per trait and each module was tested as a gene set; module *P* values were Benjamini–Hochberg-adjusted within trait and pipeline. IsoGraph modules at Leiden resolutions 2.0 and 5.0 and classical abundance-only WGCNA modules were tested; the matched switch-only and multiplex WGCNA comparators were not included in this analysis, and modules of at least 900 genes were flagged to describe the distribution of significant associations by module size (**Fig. S25**; **Data S24**).

### Stratified LD-score regression

Stratified LD-score regression^35^ tested whether common-variant heritability was concentrated in switch-gene QTL annotations conditional on baselineLD v2.2^36^. Across 13 GTEx brain tissues we constructed binary annotations for sQTL variants linked to selected switch genes, eQTL variants linked to the same genes and their union. Disease annotations used the schizophrenia-selected switch genes and aging annotations the aging-selected switch genes defined in the colocalization section. GTEx GRCh38 variants were mapped to GRCh37 using the GTEx v8 lookup table and restricted to autosomal variants in the European LD panel.

LD scores were computed from 1000 Genomes Phase 3 European genotypes and GWAS statistics were restricted to HapMap3 variants^95^. For each trait we fit baselineLD plus sQTL, baselineLD plus eQTL, baselineLD plus their union and a joint baselineLD+sQTL+eQTL model. The union was not entered with its components. The primary statistic was the one-sided value for a positive conditional per-SNP heritability coefficient, Benjamini–Hochberg corrected across the 15 single-annotation tests (five traits by sQTL, eQTL, and union annotations of the aging switch genes); joint models and the schizophrenia-module annotations were not included in this family, and conventional enrichment estimates were retained as descriptive outputs (**Data S22**).

### Candidate RBP regulons

GENCODE v47 mature-transcript sequences and strand-aware sequences from the first and last 100 nucleotides of annotated introns were scanned against human ATtRACT position-weight matrices (PWMs) with MOODS^96,97^. Uracil was represented as thymine, only the sense strand was scanned and matrices were converted to log odds with pseudocount 0.1. The composition-adjusted scan used backgrounds binned by transcript GC content (20 bins) and purine content (two bins); a uniform-background scan was retained as a sensitivity analysis. A per-position match *P* < 10^−4^ defined a motif hit. Motif families were defined by average-linkage clustering at distance 0.25, where distance was one minus the maximum mean column-wise PWM correlation over ungapped offsets with at least four overlapping columns. Only the sense orientation was compared. Hits were summed within motif family before deriving presence/absence calls; family membership could span multiple RBPs.

A gene carried an RBP site switch when a motif occurred in exactly one member of an observed transcript pair. Candidate regulons were tested both by hypergeometric over-representation against the region-wide switch-gene pool and by a binomial generalized linear model with logit link. The binary response was whether a gene carried a site switch for the tested RBP; the predictor of interest was module membership, adjusting motif opportunity for transcript length, GC content, UTR and coding composition, and transcript count. Modules with fewer than three switch genes and non-estimable separated cells were excluded. Benjamini–Hochberg correction was applied across the complete region-by-module-by-RBP family. The reported analysis used mature-transcript sequences and individual RBPs; intronic-flank and motif-family arms were sensitivity analyses. A module-by-RBP cell was reported as supported when it was significant by both tests (*q* < 0.05) with an adjusted odds ratio above 1 (**Fig. S20**; **Data S13**). Public ENCODE eCLIP^27,28^ provided orthogonal evidence of binding capacity. Because these data come from HepG2 and K562 cells, they indicate whether an RBP can occupy the relevant intervals, not regulation in brain. Motif and eCLIP results nominated candidate regulators only.

ENCODE support used genomic overlap of the union of available HepG2/K562 peaks with switched versus shared exonic intervals. Binary binding calls were compared by a two-sided exact McNemar test over unique nominated genes for each RBP, with Benjamini–Hochberg correction across testable RBPs; repeated region–module rows were descriptive rather than independent tests. An RBP was binding-supported when switched intervals were bound more often than constitutive ones at FDR ≤ 0.05 (**Data S14**).

### Statistical analysis and reproducibility

The statistical unit was the donor for cohort-level molecular analyses, the gene for gene-level enrichment and conditional models, the module for module-level tests, and the independently generated dataset for synthetic method comparisons. Sample sizes were determined by the available cohorts after prespecified quality control; no prospective power calculation was used. No observations were removed on the basis of the tested outcome. Exact correction families, random seeds, and significance thresholds were archived with each analysis. Empirical *P* values were calculated using numerator and denominator pseudo-count adjustments to avoid zero-valued estimates. Effect sizes and confidence intervals are reported alongside *P* values wherever available.

Primary analyses were executed in Python 3.12 (production environment Python 3.12.13) using IsoGraph v0.1.5 on the Pittsburgh Supercomputing Center Bridges-2 system, with allele-aware recount workflows completed on the Northwestern University Genomics Compute Cluster (Quest). The core Python stack comprised NumPy 2.4.4^98^, pandas 2.3.3^99^, SciPy 1.17.1^100^, scikit-learn 1.8.0^101^, PyTorch 2.11.0, python-igraph 1.0.0^102^, leidenalg 0.11.0, statsmodels 0.14.6^103^, and pyarrow 24.0.0.

Statistical QTL mapping, DTU, and colocalization analyses were conducted in R 4.4.3 using coloc 5.2.3^2 0^, susieR 0.14.2^21^, WGCNA 1.73^104^, and satuRn 1.18.0^8^. Figure generation was performed in R 4.5.3 using ggplot2 4.0.3^105^ and patchwork 1.3.2. Complete software environment lock files and container specifications are archived within the analysis repository.

## Data availability

No new biospecimens were collected. BrainSEQ RNA-sequencing reads, sample metadata, and SNP-array genotypes are available from the BrainSEQ Consortium under the controlled-access terms of the original studies^15,16^. GTEx v11 transcript expected counts, gene read counts, sample and subject annotations, *cis*-sQTL and *cis*-eQTL summary statistics, SuSiE credible-set summaries, and the GTEx v8 variant lookup table are available from the GTEx Portal (https://www.gtexportal.org/home/downloads/adult-gtex)^17,47^. Raw BrainSEQ and other controlled-access source files are not redistributed.

Reference resources used in this study are publicly available: GENCODE release 47 on GRCh38.p14 (https://www.gencodegenes.org/human/release_47.html), 1000 Genomes Project Phase 3 sites and the European LD panel (https://www.internationalgenome.org), TOPMed Freeze 10 through the TOPMed Imputation Server (https://imputation.biodatacatalyst.nhlbi.nih.gov), baselineLD v2.2 (https://alkesgroup.broadinstitute.org/LDSCORE/), Gene Ontology annotations (https://geneontology.org), ATtRACT RBP matrices (https://attract.cnic.es), gnomAD v4.1 constraint metrics (https://gnomad.broadinstitute.org/downloads), ClinVar (https://www.ncbi.nlm.nih.gov/clinvar/) and ENCODE eCLIP peaks (https://www.encodeproject.org). GWAS summary statistics were obtained from the Psychiatric Genomics Consortium, the GWAS Catalog and the original publications cited in Methods. Long-read Oxford Nanopore transcript quantifications of aged human DLPFC were obtained from Aguzzoli Heberle et al.^18^; the Bambu count matrix is deposited at Zenodo^83^ and was not regenerated here.

Processed IsoGraph dataset bundles, lean result tables, provenance records, and large deterministic analysis artifacts will be deposited in a public Zenodo archive at publication. Until that deposit is minted, processed manifests and reproducibility metadata are version-controlled in the analysis repository (https://github.com/heart-gen/isograph-brain-aging-benchmarking). Source data for the figures will be provided with the published article. Full-length supplementary tables are provided as Data S1–S28, each an Excel workbook with a README sheet and an accompanying CSV copy. Further information is available from the corresponding author on request.

## Code availability

IsoGraph is available at https://github.com/heart-gen/IsoGraph, on the Python Package Index as `isograph` and archived at Zenodo^70^. Documentation is maintained at https://github.com/heart-gen/<IsoGraph/wiki >and https://isograph.readthedocs.io/en/latest/. Analysis and benchmarking code for this manuscript, including the scripts that generate every figure and supplementary table, is available at https://github.com/heart-gen/isograph-brain-aging-benchmarking. The version of the analysis and benchmarking code associated with this manuscript will be archived on Zenodo, with its DOI provided at publication.

## Acknowledgements

We thank the BrainSEQ Consortium and the Lieber Institute for Brain Development for access to the BrainSEQ RNA-sequencing and genotype data, and especially the brain donors and their families whose contributions made these resources possible. The GTEx Project was supported by the Common Fund of the Office of the Director of the National Institutes of Health, and by NCI, NHGRI, NHLBI, NIDA, NIMH, and NINDS. The GTEx data used for the analyses described in this manuscript were obtained from the GTEx Portal on January 15, 2026.

High-performance computing resources were provided by the Bridges-2 system at the Pittsburgh Supercomputing Center through allocation BIO260136 from the Advanced Cyberinfrastructure Coordination Ecosystem: Services & Support (ACCESS) program, which is supported by National Science Foundation grants 2138259, 2138286, 2138307, 2137603, and 2138296. Additional computational resources and staff contributions were provided by the Genomics Compute Cluster on Quest (Northwestern University’s high-performance computing facility), which is jointly supported by the Feinberg School of Medicine, the Center for Genetic Medicine, Feinberg’s Department of Biochemistry and Molecular Genetics, the Office of the Provost, the Office for Research, the Weinberg College of Arts and Sciences, and Northwestern Information Technology.

This work was supported by the National Institute on Minority Health and Health Disparities of the National Institutes of Health (NIH) under award number R00MD016964, by Alzheimer’s Association grant 25AARG-1413315, and by a Brain & Behavior Research Foundation Young Investigator Grant to K.J.M.B. This work is the result of NIH funding, in whole or in part, and is subject to the NIH Public Access Policy.

## Author contributions

K.J.M.B. conceived and designed the study, developed IsoGraph, performed the analyses, acquired funding, and wrote the original draft. A.B. and E.K.J. contributed to code generation and figure preparation. All authors reviewed, edited, and approved the final manuscript.

## Supplementary Information

### Supplementary Figures

**Fig. S1.**
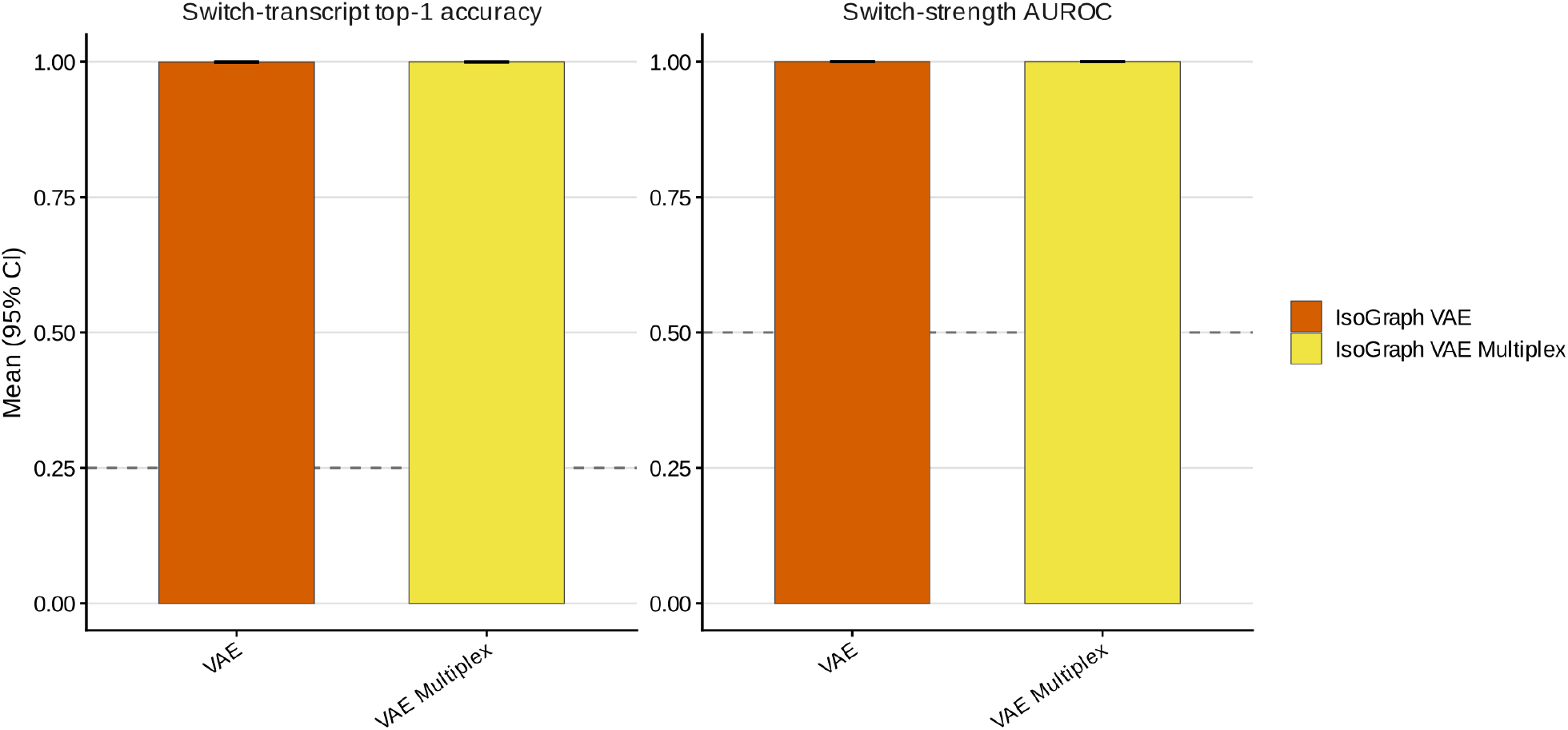
Module interpretation recovers planted switch transcripts and switching genes. Left, switch-transcript top-1 accuracy: the fraction of switching genes whose highest-loading transcript is the planted switch transcript, evaluated in multi-isoform simulations (four transcripts per gene; dashed line, chance = 0.25). Right, switch-strength area under the receiver operating characteristic curve (AUROC) for separating switching from non-switching genes within modules (dashed line, chance = 0.5). Bars show means across simulations for IsoGraph VAE (variational autoencoder; switch-only) and IsoGraph VAE Multiplex; error bars indicate 95% bootstrap confidence intervals.

**Fig. S2.**
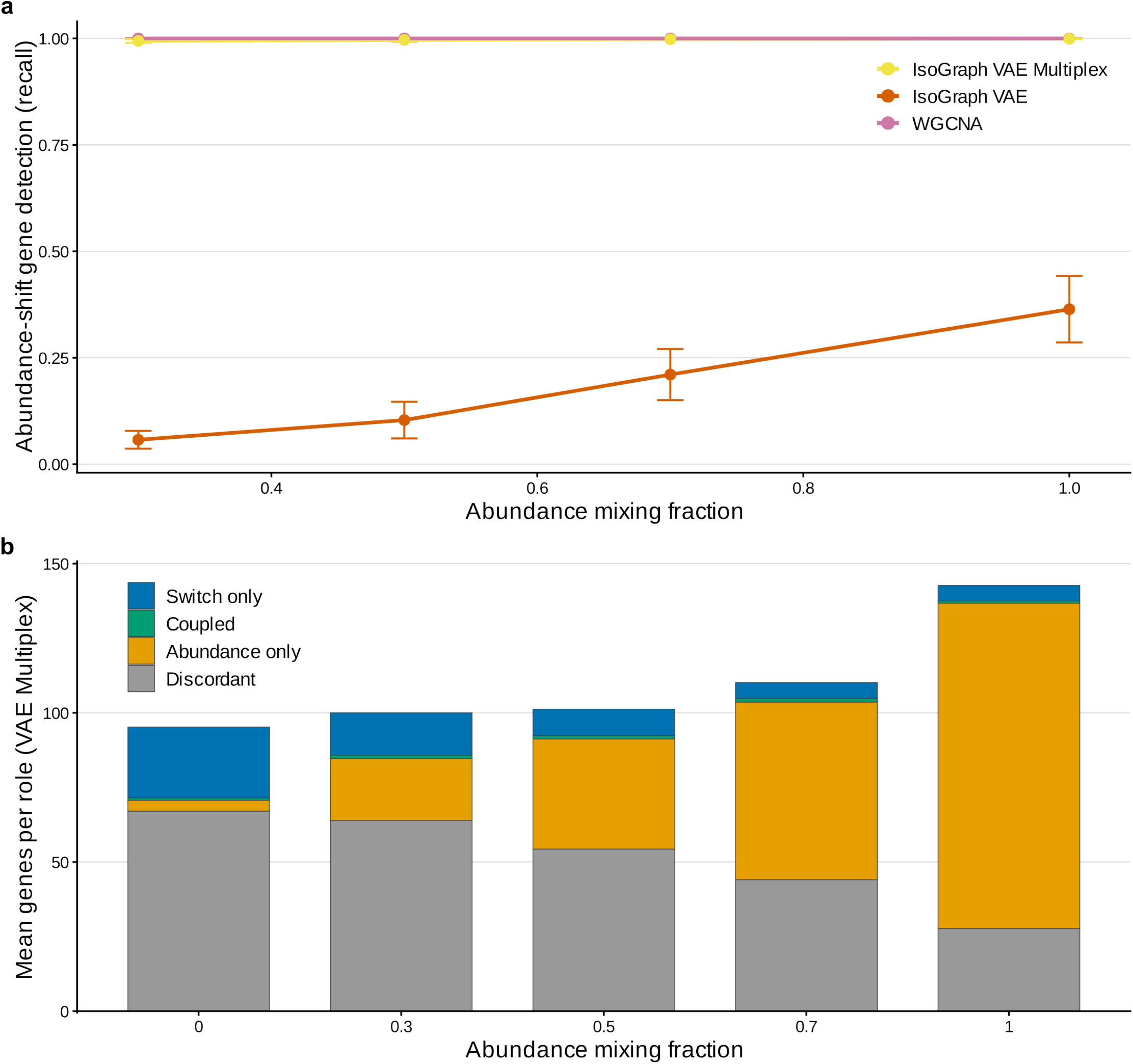
Mixed switch–abundance simulations: abundance-shift detection and channel-specific roles. Simulations mixed planted isoform switching with abundance shifts at increasing abundance mixing fractions. **a**. Recall of genes carrying a planted abundance shift for IsoGraph VAE (variational autoencoder), IsoGraph VAE Multiplex, and weighted gene co-expression network analysis (WGCNA). Points show means across simulation seeds and error bars indicate 95% confidence intervals. **b**. Mean number of genes assigned to each channel-specific role by IsoGraph VAE Multiplex: switch only, coupled (switch and abundance), abundance only, and discordant.

**Fig. S3.**
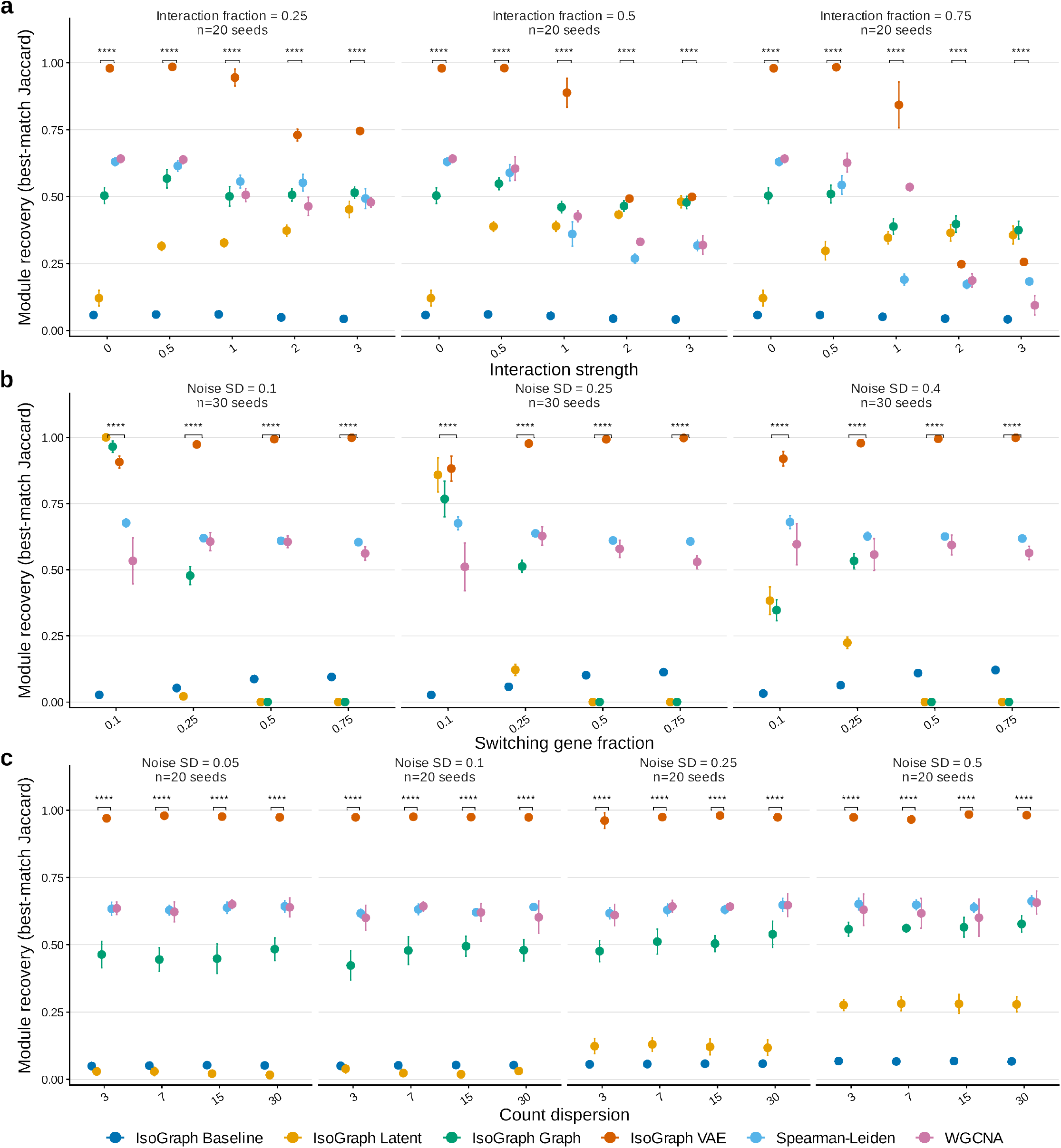
Module recovery under feature interactions, noise, and count dispersion. **a**. Recovery as a function of interaction strength between the switch and abundance features, at interaction fractions of 0.25, 0.5 and 0.75 (20 seeds per setting). **b**. Recovery as a function of the fraction of switching genes at three noise levels (30 seeds per setting). **c**. Recovery as a function of count dispersion at four noise levels (20 seeds per setting). Module recovery is the best-match Jaccard similarity between each planted module and its closest inferred module (1 = perfect recovery). Points show means across simulation seeds and error bars indicate 95% confidence intervals. Asterisks mark two-sided Wilcoxon rank-sum comparisons of IsoGraph VAE (variational autoencoder) with weighted gene co-expression network analysis (WGCNA) at each parameter value, Benjamini–Hochberg-adjusted within each panel (**** adjusted *P* [*P*_adj_] < 10^−4^); these per-panel comparisons are for visual guidance and are separate from the full-family tests in Table S1.

**Fig. S6.**
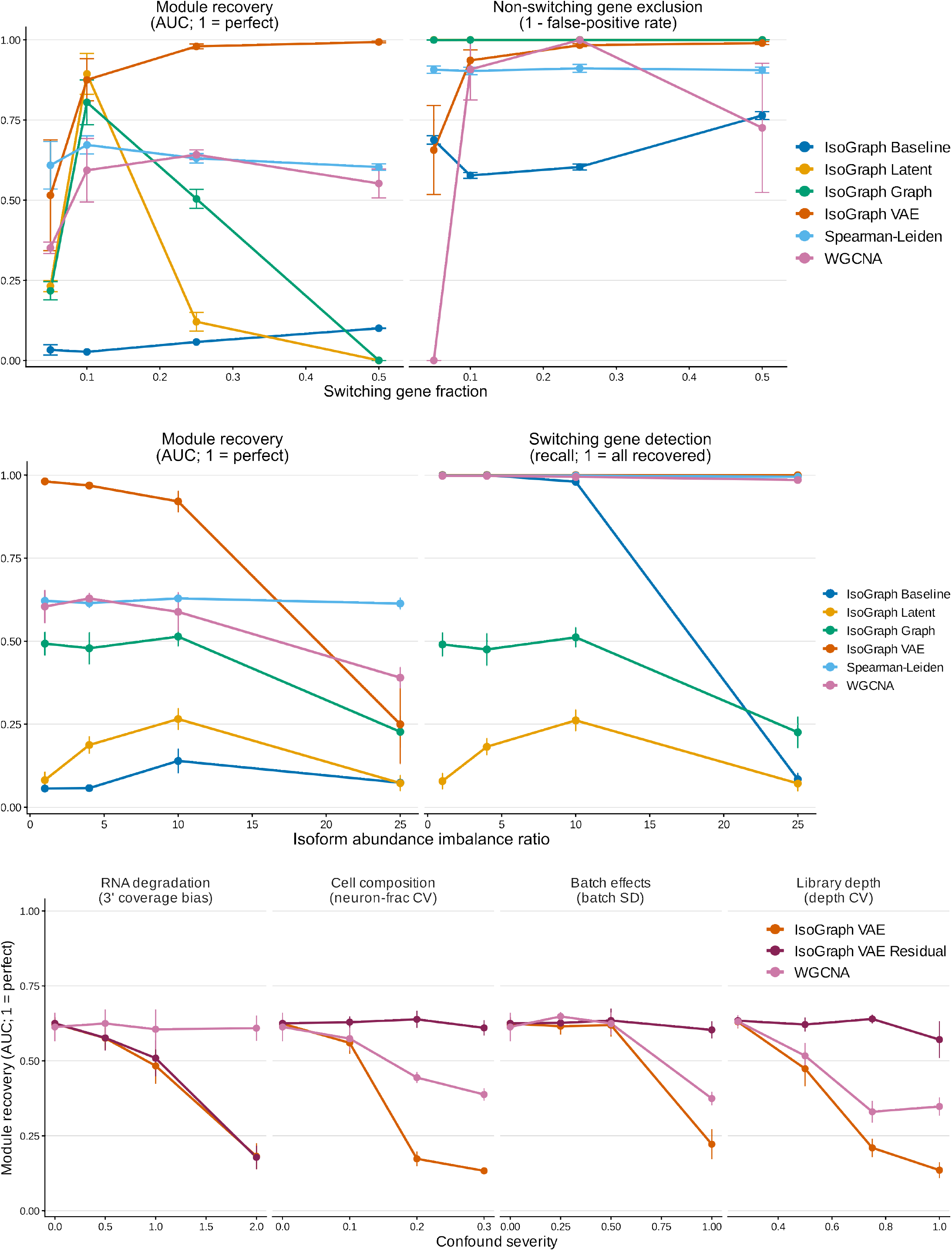
Robustness to simulated technical and compositional confounders. Module recovery as the severity of four confounders increased: RNA degradation (3′ coverage bias), cell composition (coefficient of variation of the neuronal fraction), batch effects (batch standard deviation), and library depth (coefficient of variation of depth). IsoGraph VAE (variational autoencoder), its residualized implementation (IsoGraph VAE Residual), and weighted gene co-expression network analysis (WGCNA) are shown. Module recovery is the best-match Jaccard similarity between each planted module and its closest inferred module (1 = perfect recovery). Points show means across simulation seeds and error bars indicate 95% confidence intervals.

**Fig. S7.**
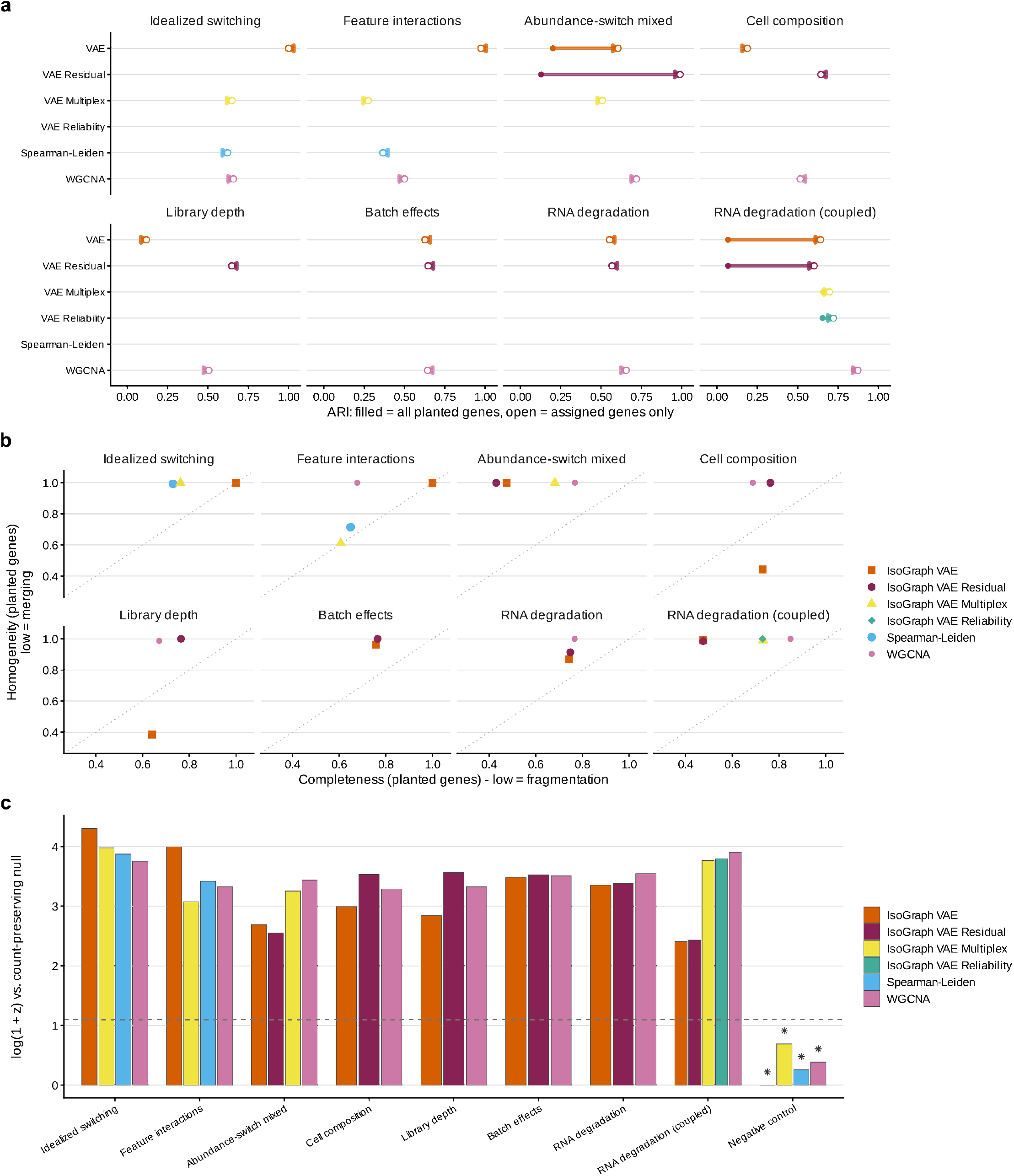
Partition diagnostics complement best-match module recovery. **a**. Adjusted Rand index (ARI) against the planted partition, computed over all planted genes (filled) or only genes assigned to a module (open). **b**. Homogeneity versus completeness for planted genes; low homogeneity indicates merging of planted modules, and low completeness indicates fragmentation. **c**. Module recovery relative to a permutation null that preserves the number of inferred modules, shown as log(1 + *z*), where *z* is the observed recovery minus the null mean, divided by the null standard deviation. The dashed line marks *z* = 2; asterisks mark a median permutation *P* > 0.05, that is, recovery fully explained by module-count structure.

**Fig. S8.**
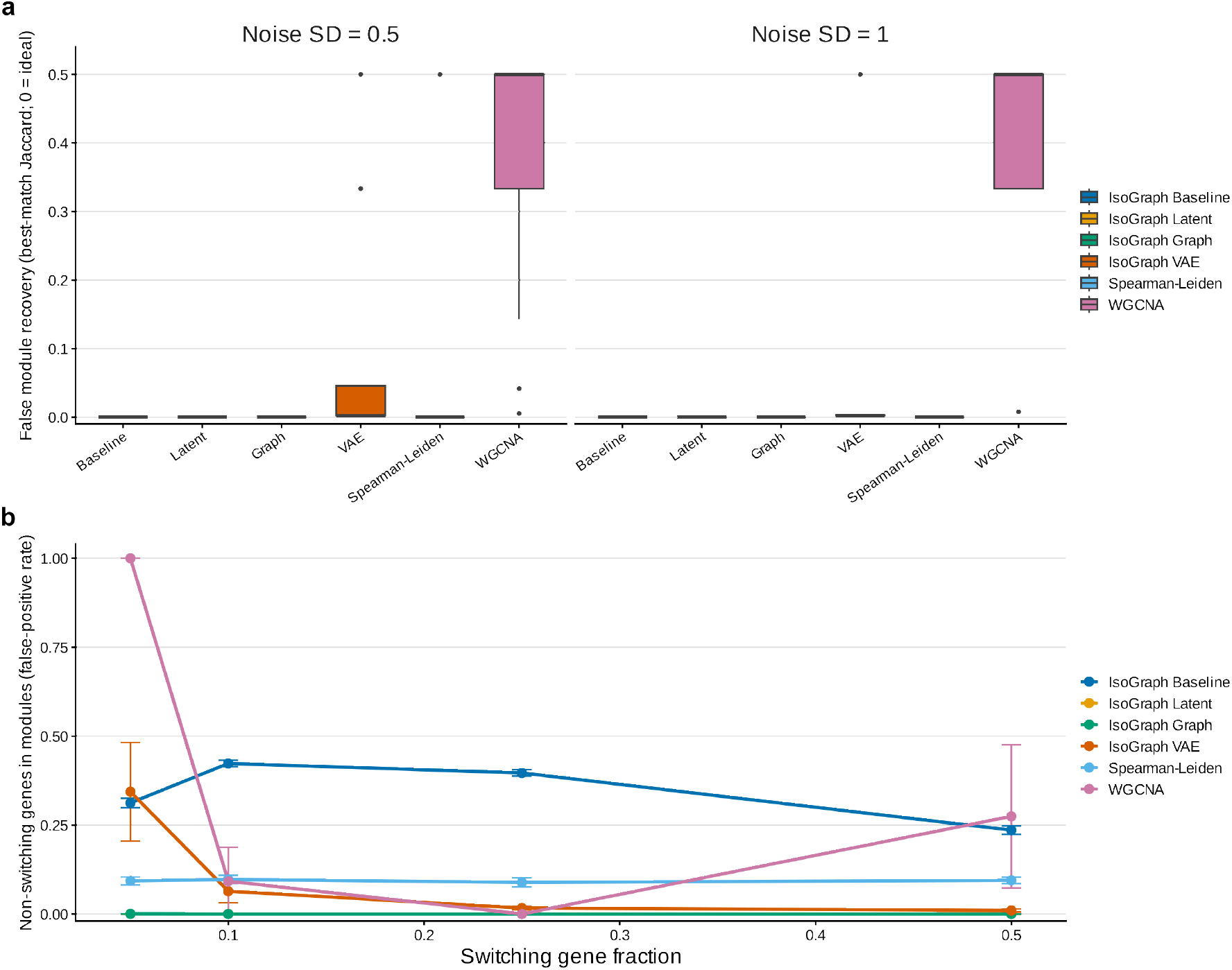
Specificity under null and non-switching settings. **a**. False module recovery in pure-noise negative-control simulations at two noise levels (0 = ideal). **b**. Fraction of non-switching genes assigned to modules (false-positive rate) as a function of the fraction of switching genes. Boxplots in **a** summarize simulation seeds; points in **b** show means across seeds, with error bars indicating 95% confidence intervals.

**Fig. S9.**
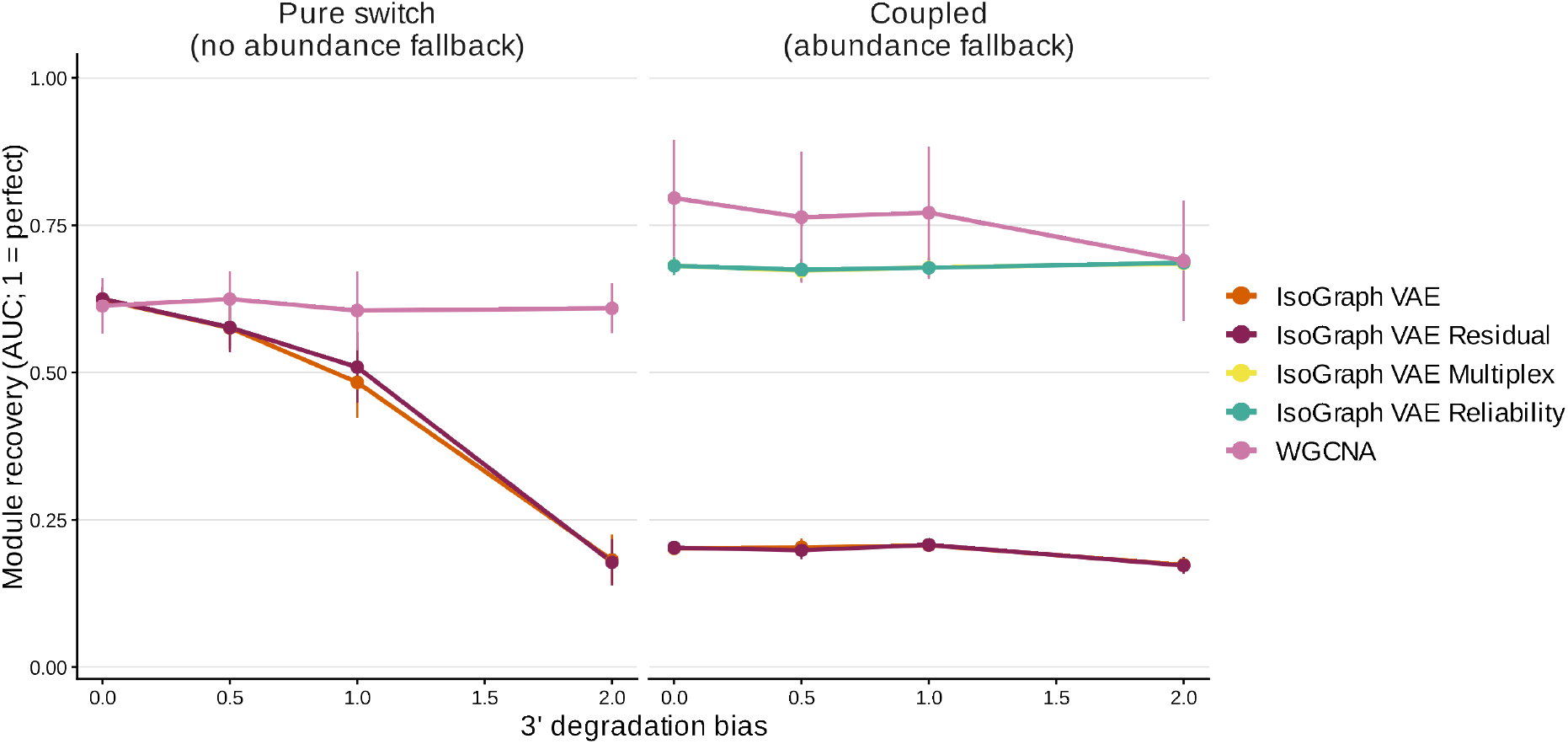
Multiplex integration provides an abundance-based fallback when RNA degradation obscures isoform-switching structure. Module recovery was evaluated across increasing 3′-degradation bias under two regulatory architectures. In the pure-switch setting, planted modules were encoded only through coordinated isoform switching and therefore lacked an abundance signal that could support recovery. In the coupled setting, switching and gene-abundance signals were concordantly embedded in the same modules, permitting abundance-informed recovery. Performance was quantified by the mean maximum Jaccard similarity to planted modules, with 1 indicating perfect recovery. IsoGraph VAE (variational autoencoder) and its residualized implementation were evaluated in both settings and compared with weighted gene co-expression network analysis (WGCNA); the coupled setting additionally evaluated IsoGraph VAE Multiplex and reliability-weighted IsoGraph VAE. Points show mean performance across independent simulation seeds, error bars indicate 95% confidence intervals, and lines connect results obtained with the same method.

**Fig. S10.**
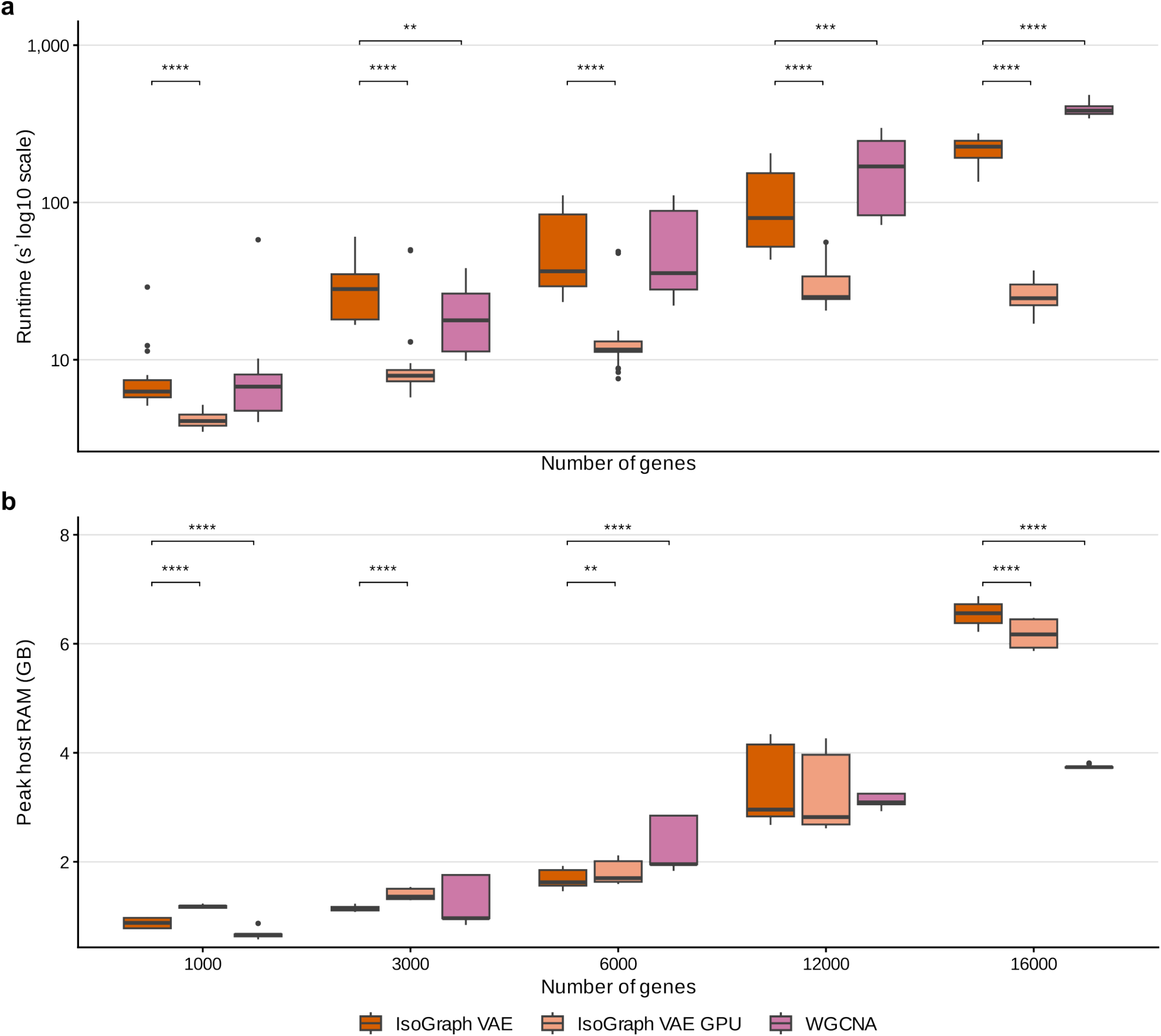
Computational scaling. **a**. Runtime (seconds, log10 scale) and **b**. peak host memory (GB) as a function of the number of genes for IsoGraph VAE (variational autoencoder) on a central processing unit (CPU), IsoGraph VAE on a graphics processing unit (GPU), and weighted gene co-expression network analysis (WGCNA). Boxplots summarize replicate runs. Asterisks mark two-sided Wilcoxon rank-sum comparisons with IsoGraph VAE (CPU) at each gene count, Benjamini–Hochberg-adjusted within each panel (** adjusted *P* [*P*_adj_] < 0.01; *** *P*_adj_ < 0.001; **** *P*_adj_ < 10^−4^). Summary statistics are given in Table S2.

**Fig. S11.**
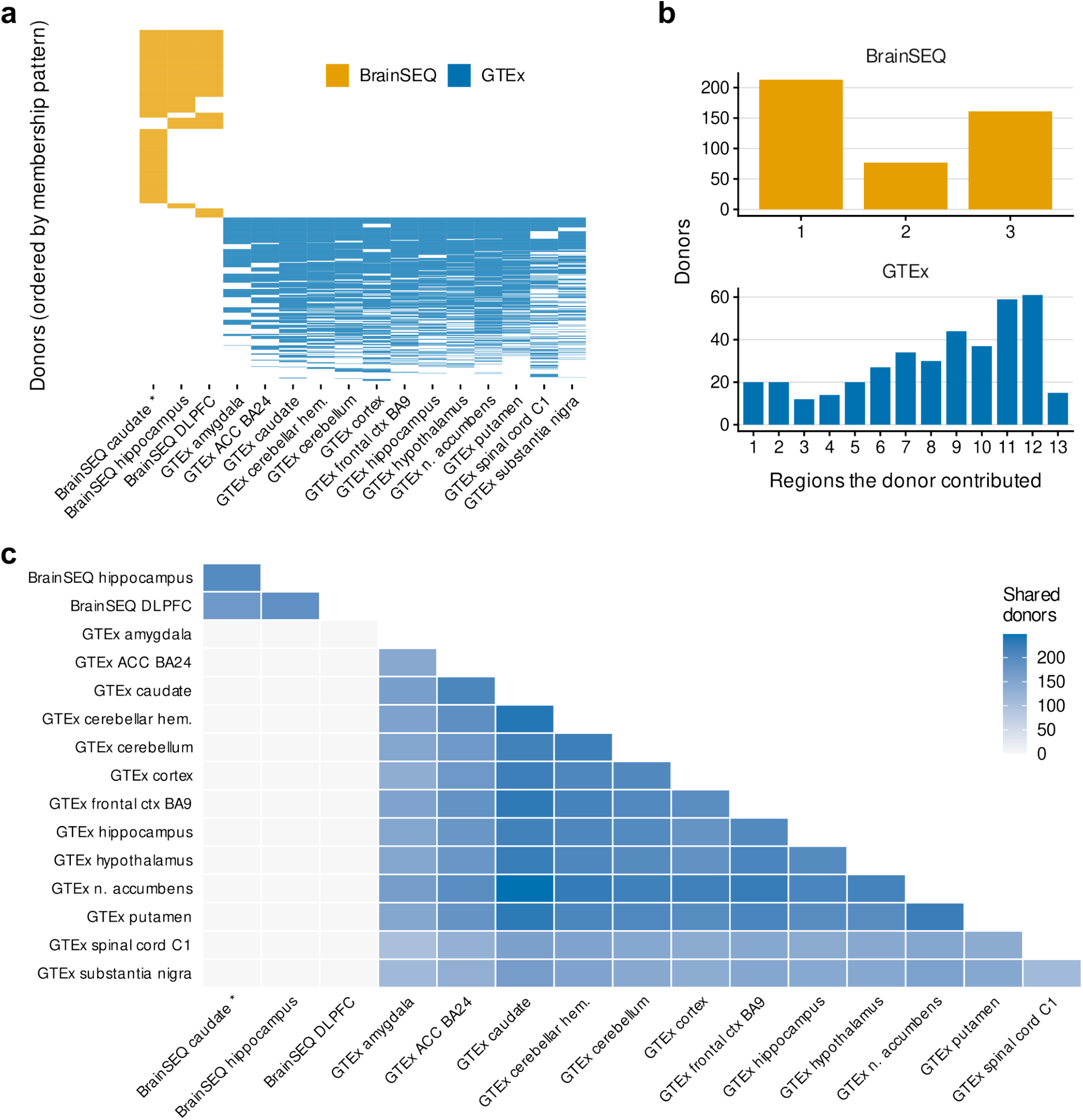
Donor sharing across the 16 brain regions and 17 analyses. **a**. Donor-by-region incidence. Each row is a donor and each column a region; a filled cell indicates that the donor contributed tissue to that region. Donors are ordered by the set of regions they contributed, so shared-donor blocks are contiguous. Asterisks mark a region analyzed more than once. **b**. Number of regions each donor contributed, by cohort. **c**. Shared-donor counts for every pair of regions; blank cells indicate no shared donors. The 16 regions comprise three BrainSEQ and 13 Genotype-Tissue Expression (GTEx) regions, and are covered by 17 analyses because BrainSEQ caudate is analyzed twice: once for aging over its 238 control samples and once for schizophrenia over those same 238 control samples together with 152 samples from donors with schizophrenia. The 4,322 sample-analysis rows therefore correspond to 4,084 distinct tissue samples from 844 donors. Donor identifiers (BrNum for BrainSEQ, SUBJID for GTEx) are shared across regions within a cohort and are never matched across cohorts, whereas the sample identifier (RNum for BrainSEQ) is specific to a donor and region. The median GTEx donor contributed 9 of the 13 GTEx regions and the median BrainSEQ donor 2 of the 3 BrainSEQ regions, so agreement between regions within a cohort is re-measurement of a shared donor pool, whereas BrainSEQ-versus-GTEx comparisons carry independent donors.

**Fig. S12.**
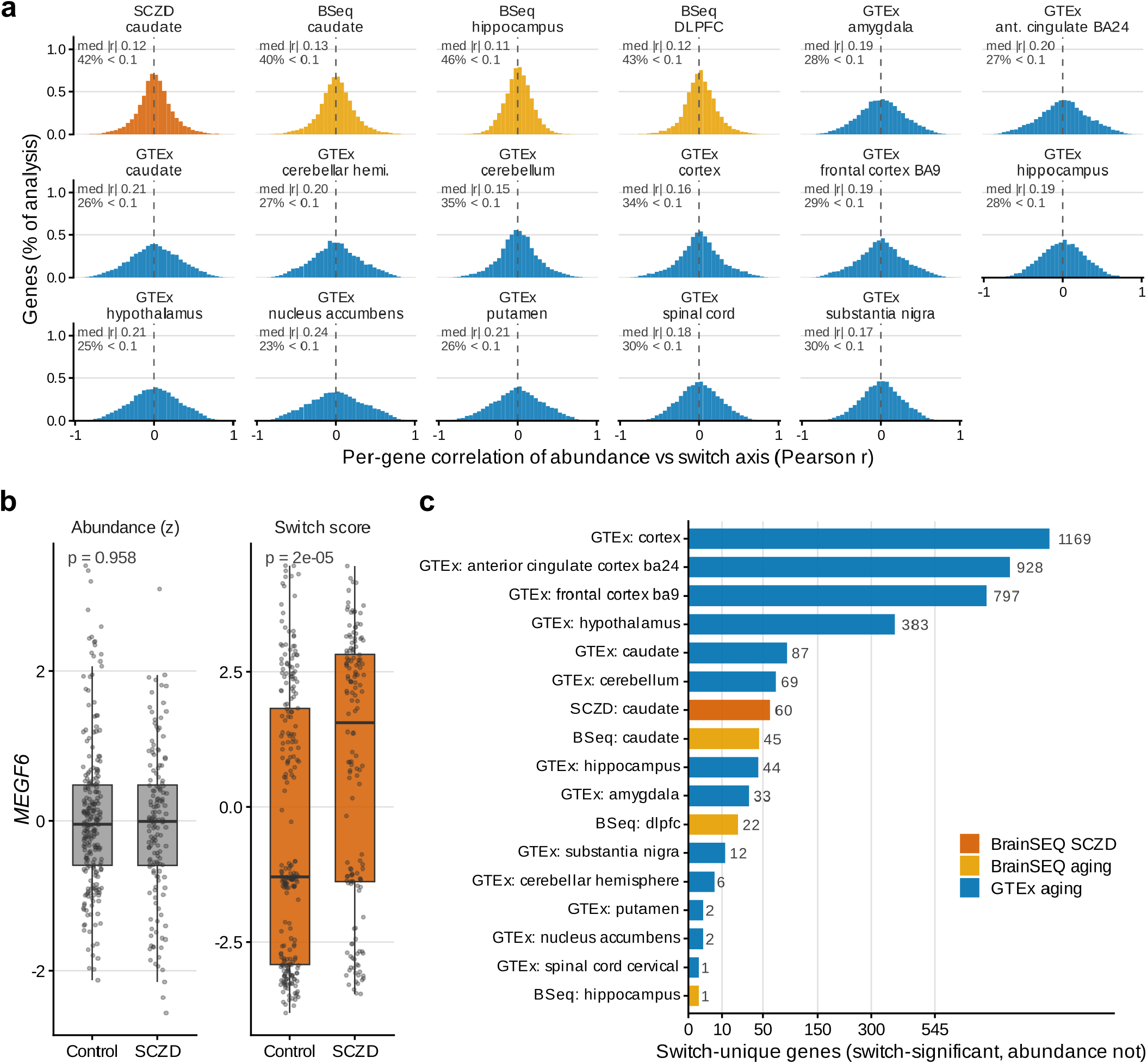
The isoform-switch and gene-abundance channels are partly non-redundant. **a**. Distribution of per-gene Pearson correlations between the gene-abundance and isoform-switch coordinates in each of the 17 analyses (12,042– 13,222 genes per analysis; the *y* axis is the percentage of that analysis’s genes per bin). Each panel is annotated with its median |*r*| and the percentage of genes with |*r*| < 0.1. **b**. *MEGF6* shows schizophrenia-associated isoform usage in caudate without a detectable conditional gene-abundance association. **c**. Switch-unique genes – those whose switch coordinate is phenotype-associated conditional on abundance while the corresponding abundance test is not – for all 17 analyses, including the five Genotype-Tissue Expression (GTEx) regions that are absent from the composition analyses in Fig. 2c,d because they were not deconvolved. The *x* axis is square-root scaled.

**Fig. S13.**
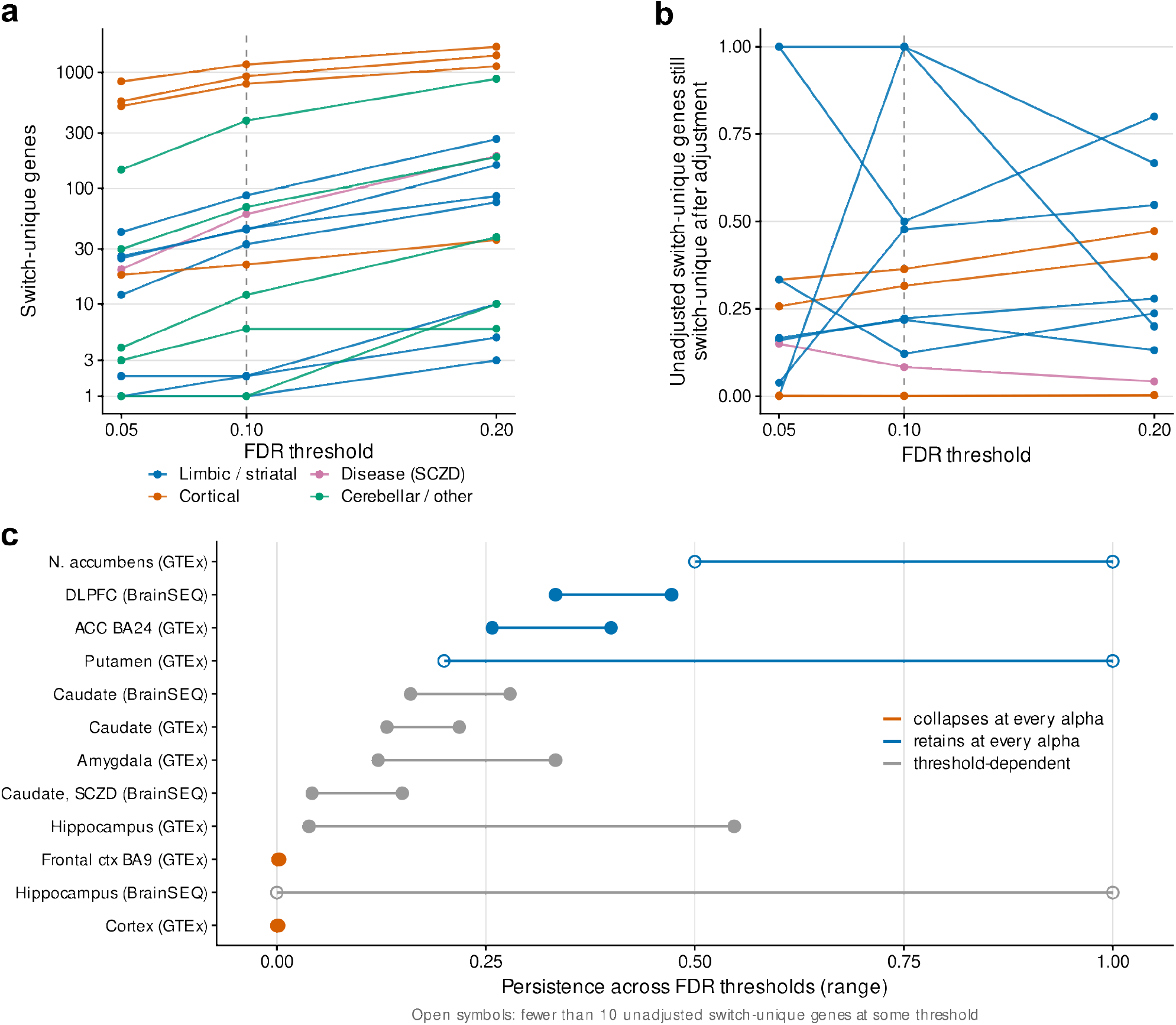
Sensitivity of the switch-unique classification to the FDR threshold. Genes were reclassified at false discovery rate (FDR) thresholds of 0.05, 0.10 (the value used throughout), and 0.20. Because the Benjamini–Hochberg-adjusted values are computed once across all genes, changing the threshold moves only the cut and no model is refit. **a**. Switch-unique genes per analysis against threshold; the *y* axis is pseudo-log scaled and the dashed line marks 0.10. **b**. Gene-level persistence after cell-composition adjustment against threshold, for the 12 analyses with a composition-adjusted arm. **c**. Range of persistence across the three thresholds per analysis, colored by whether the analysis retains, collapses, or changes call between thresholds. Open symbols mark analyses with fewer than 10 unadjusted switch-unique genes at some threshold, where persistence is a ratio of single-digit counts. The ranking of analyses by switch-unique count is essentially unchanged across thresholds (Spearman ρ = 0.97–0.98 against 0.10), and the two cortical collapses are threshold-independent (persistence ≤ 0.004 at every threshold in Genotype-Tissue Expression (GTEx) cortex and frontal cortex BA9, versus ≥ 0.26 at every threshold in anterior cingulate cortex BA24 and BrainSEQ dorsolateral prefrontal cortex [DLPFC]). The ordering of the intermediate analyses is not threshold-independent, and GTEx hippocampus in particular ranges from 0.04 to 0.55 across the three thresholds.

**Fig. S14.**
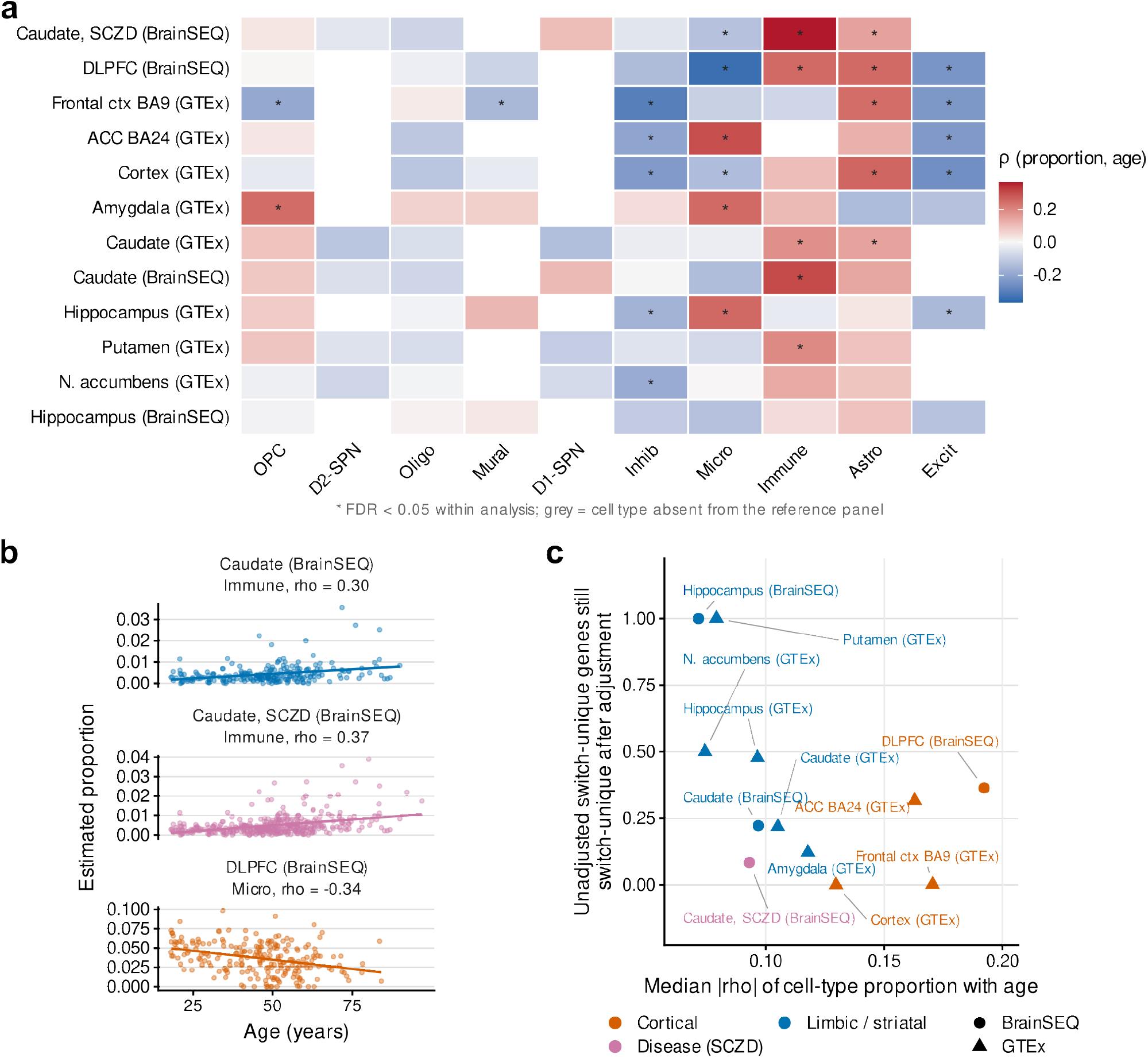
Estimated cell-type composition is itself age-coupled, but coupling alone does not predict which analyses lose their switch-unique genes. **a**. Spearman correlation between each estimated cell-type proportion and donor age, per analysis; asterisks mark false discovery rate (FDR) < 0.05 within analysis, and gray cells indicate cell types absent from that region’s reference panel. **b**. The sample-level scatter behind the strongest age-coupled cell type in each anatomical class, with an ordinary least-squares line. **c**. Gene-level persistence of the switch-unique set against the analysis’s median absolute correlation of cell-type proportion with age. Median absolute correlation is used rather than a count of significant cell types because reference panels differ in size between regions. Cortical analyses carry the most age-coupled composition, yet anterior cingulate cortex BA24 and BrainSEQ dorsolateral prefrontal cortex (DLPFC) retain about a third of their switch-unique genes while frontal cortex BA9 and cortex retain almost none, so composition–age coupling bounds the risk of over-adjustment without resolving it. **d**. Module-by-cell-type marker tests reaching FDR < 0.05 for enrichment and for depletion of cell-type marker genes among module members, pooled by cohort, with the number significant over the number tested. This control does not depend on the covariate model. Only 12 of 2,061 tests showed enrichment and none showed depletion, indicating that modules were not generally enriched for the tested marker sets. This result does not exclude contributions of cellular composition to module structure or establish which biological signals composition adjustment removes.

**Fig. S15.**
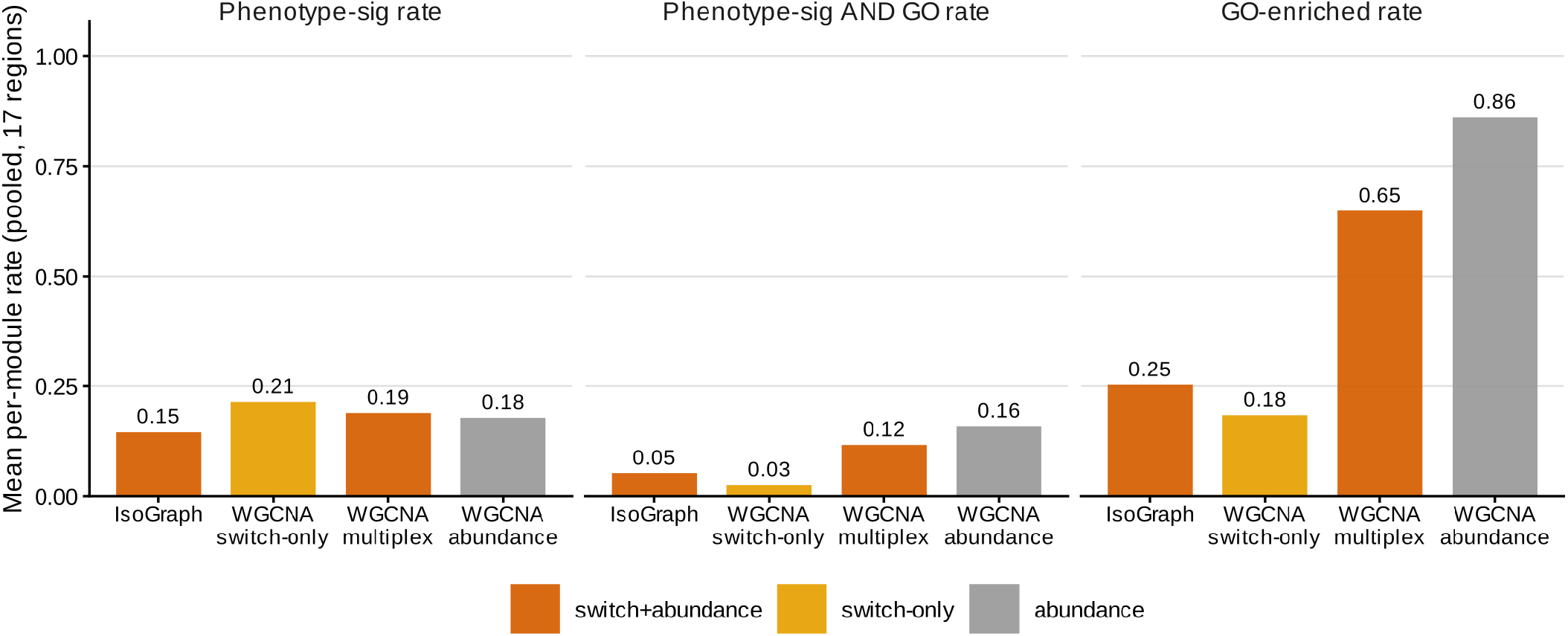
Per-module rates for the four module sets, on identical samples. Each analysis was partitioned four ways: IsoGraph on switch and abundance features, and classical weighted gene co-expression network analysis (WGCNA) on the same switch-only features, on the same switch and abundance features, and on abundance alone. Bars are the mean per-module rate across the 17 analyses, colored by the feature class each method consumed (switch and abundance, dark orange; switch-only, light orange; abundance, gray). Rates are per-module rather than totals because the four methods partition at different granularity (IsoGraph median 25 modules of median size 108; the WGCNA baselines 8–25 modules of size 159–468), which makes raw counts non-comparable. A module counts as phenotype-significant at module-level false discovery rate (FDR) ≤ 0.1 and as Gene Ontology (GO)-enriched with at least one enriched term. IsoGraph does not carry the highest phenotype-association rate, and the two cohorts rank the methods in opposite orders (IsoGraph 0.176 in BrainSEQ against 0.137 in Genotype-Tissue Expression [GTEx], where 13 of the 17 analyses lie), so the pooled ordering should not be read as a method ranking. This figure is the scope bound on the rest of the results: the defensible contribution of IsoGraph is the content of its modules, not a higher module-level rate.

**Fig. S16.**
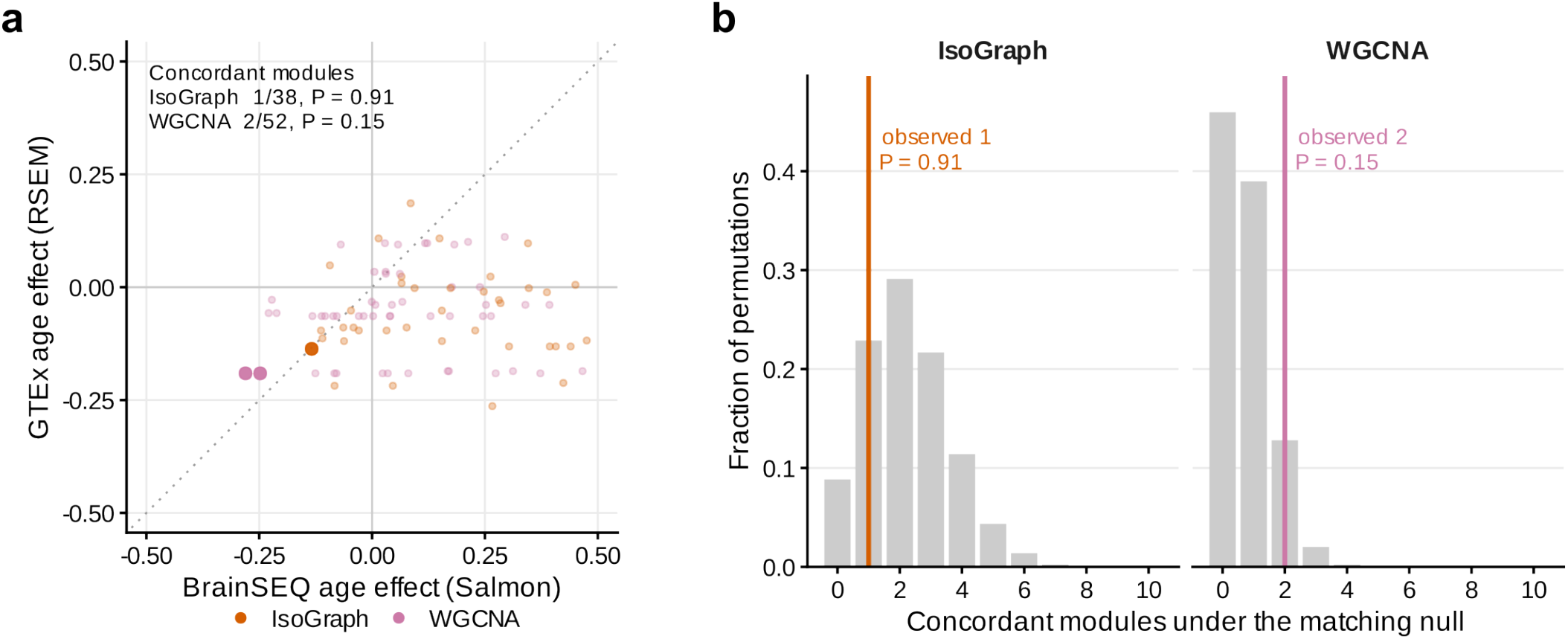
Cross-cohort module matching by gene membership, and its permutation null. **a**. Age effect of each matched BrainSEQ module against its best-overlapping Genotype-Tissue Expression (GTEx) module in the anatomically paired region, for both methods (IsoGraph and weighted gene co-expression network analysis [WGCNA]). Larger opaque points are pairs that are nominally significant in both cohorts with a concordant sign. **b**. The observed count of such pairs against the permutation null that holds every age statistic fixed and permutes which target module each source module is matched to. Neither method separates from its null. The two cohorts are quantified by different pipelines – BrainSEQ transcripts by Salmon, GTEx by RSEM – so a module whose age effect fails to carry across them has failed a test that confounds quantification with biology. This comparison is therefore reported as a limitation, and the within-cohort split-half analysis (**Fig. 3d**) carries the reproducibility claim.

**Fig. S17.**
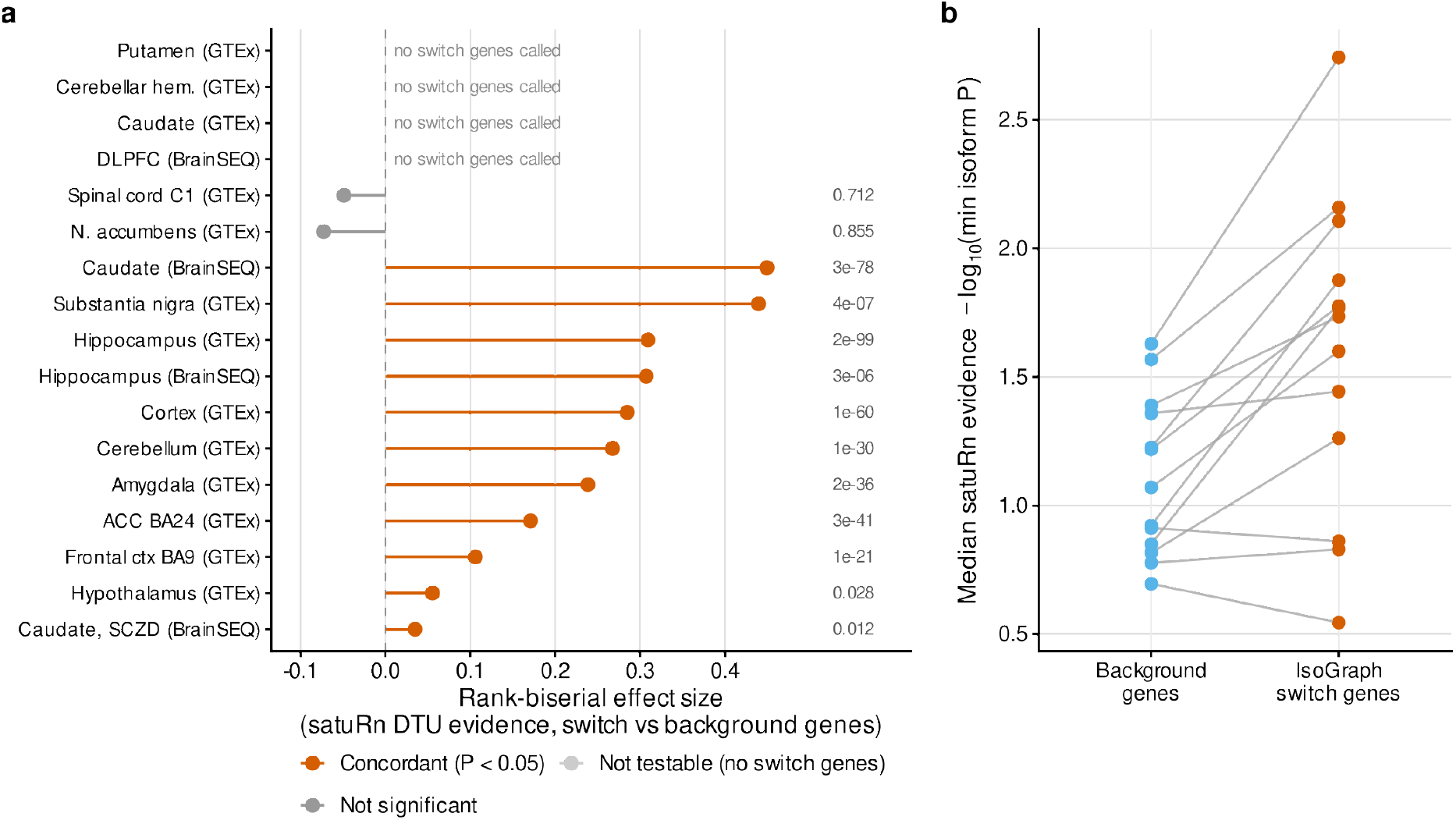
IsoGraph switch genes carry elevated DTU evidence under satuRn. satuRn, the differential transcript usage (DTU) test run by IsoformSwitchAnalyzeR v2, was fit to the same transcript quantifications with age as a single continuous coefficient, or schizophrenia (SCZD) against control in the BrainSEQ caudate disease analysis, using covariates matched to IsoGraph’s discovery design. Each gene’s evidence is −log10 of its minimum isoform *P*. **a**. Rank-biserial effect size comparing that evidence between genes in trait-associated IsoGraph switch modules and all other tested genes, for each of the 17 analyses. Labels give the one-sided Mann–Whitney *P*. Four analyses had no switch genes; they are shown as not testable rather than as null results, so the denominator is 13. **b**. Median satuRn evidence for background and switch genes in each testable analysis, with lines joining the two values from the same analysis. satuRn’s empirical-null false discovery rate (FDR) is not used to binarize the calls: it assumes sparse differential usage, which pervasive age-related DTU in brain violates. Because both methods use the discovery transcript counts, this figure shows agreement between analytical approaches, not independent replication.

**Fig. S18.**
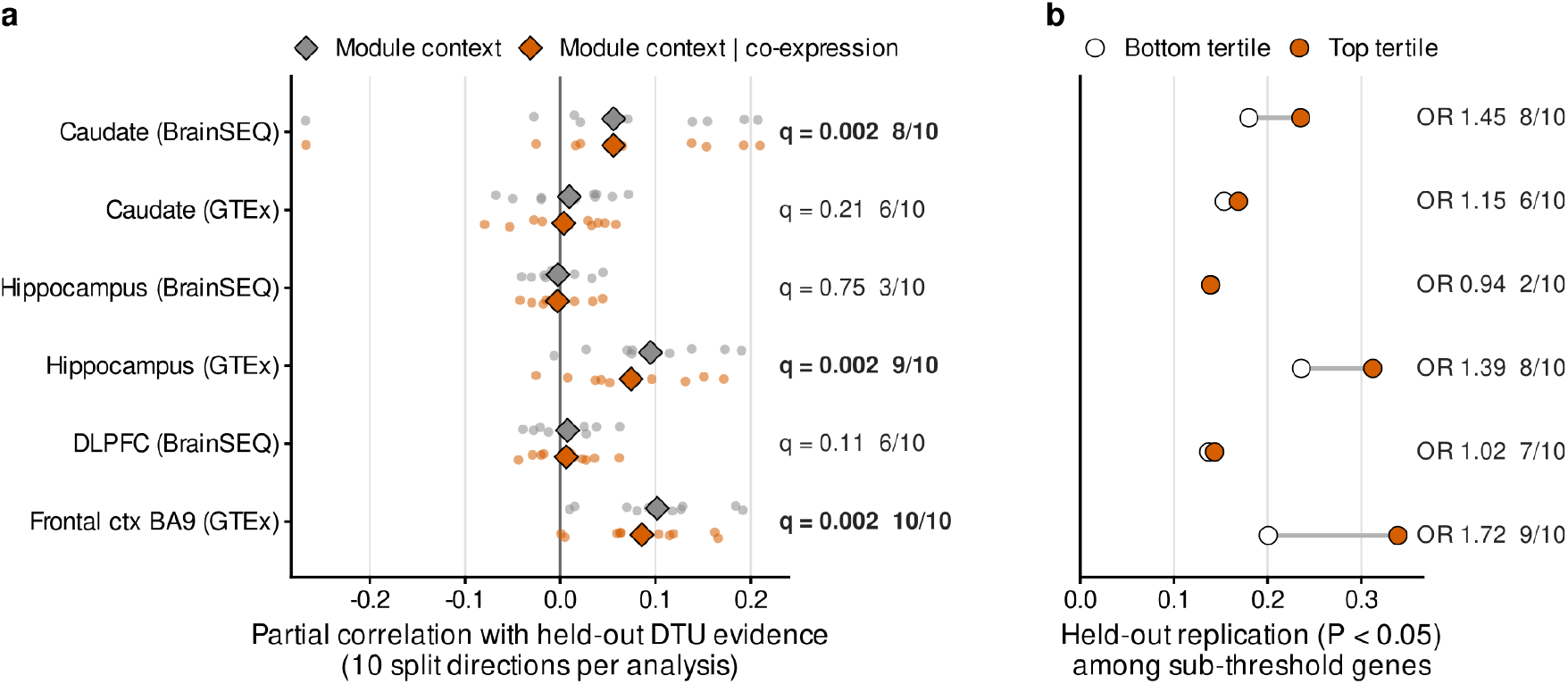
Module context predicts held-out DTU evidence in three of six analyses. satuRn was fit separately in the two donor halves of five random splits of each of the six anatomically paired cohort-region analyses, giving 10 discovery-to-held-out directions per analysis. **a**. Partial correlation between a gene’s module context (the mean discovery-half differential transcript usage [DTU] evidence of its module neighbors, leaving the gene out) and its held-out DTU evidence. The correlation is conditioned on a natural spline of the gene’s own discovery evidence, isoform count, expression and minor-isoform usage (gray), and additionally on the discovery evidence of its 50 most co-expressed genes (orange). Small points are individual split directions and diamonds their mean, which is the test statistic. The labels give the Benjamini–Hochberg *q* of the co-expression-adjusted mean against 1,000 permutations of module labels within technical strata, and the number of positive directions; bold marks *q* < 0.05. **b**. Practical reading of **a**. Among genes that did not meet the discovery DTU criterion (Benjamini–Hochberg *q* ≥ 0.05), the fraction reaching *P* < 0.05 in the held-out half, for the bottom (white) and top (orange) tertiles of module context. Labels give the median odds ratio over directions, adjusted for the gene’s own discovery evidence and technical covariates, and the number of directions with an odds ratio above 1. This panel is descriptive; **a** is the inference. The held-out threshold is nominal and does not define false discovery rate (FDR)-significant replication. Module neighbors share discovery-half sampling noise with the gene, and factors shared by both donor halves can contribute to the association.

**Fig. S19.**
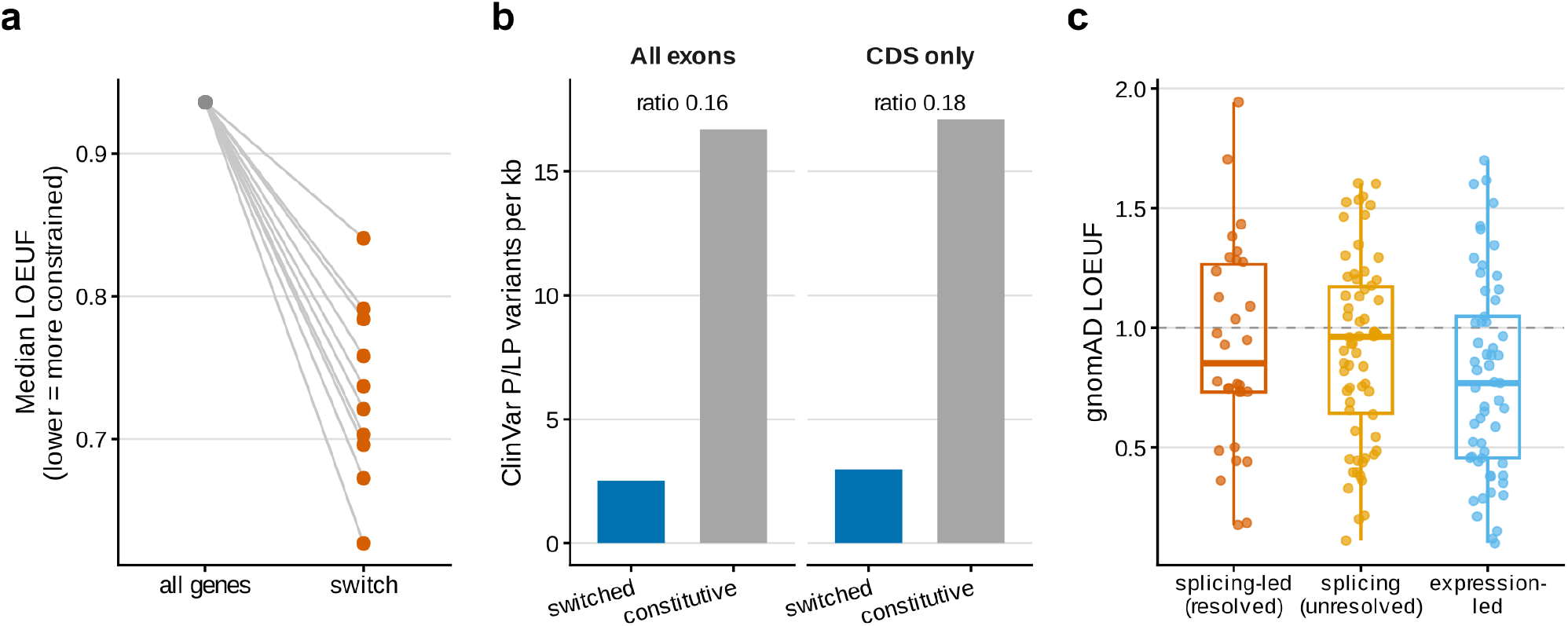
Switch genes are constrained, but switched exons are depleted of known pathogenic variants. **a**. Median gnomAD v4.1 loss-of-function observed/expected upper-bound fraction (LOEUF) of switch genes in each of the ten annotatable analyses (orange) against the genome-wide median (gray); lower values indicate stronger loss-of-function constraint. **b**. ClinVar pathogenic or likely-pathogenic entry density per kilobase in exons unique to one member of a switch pair (switched) versus other exons of the same transcripts (constitutive), pooled across analyses, for all exons and for coding exons only; labels give the median within-analysis density ratio. Densities count variant call format (VCF) entries, not unique alleles. **c**. Gene-level LOEUF of colocalization-nominated genes by nomination class (splicing-led with a resolved switch pair, splicing-linked but unresolved, expression-led); this panel is descriptive. The dashed line marks LOEUF 1 (**Table S5**).

**Fig. S20.**
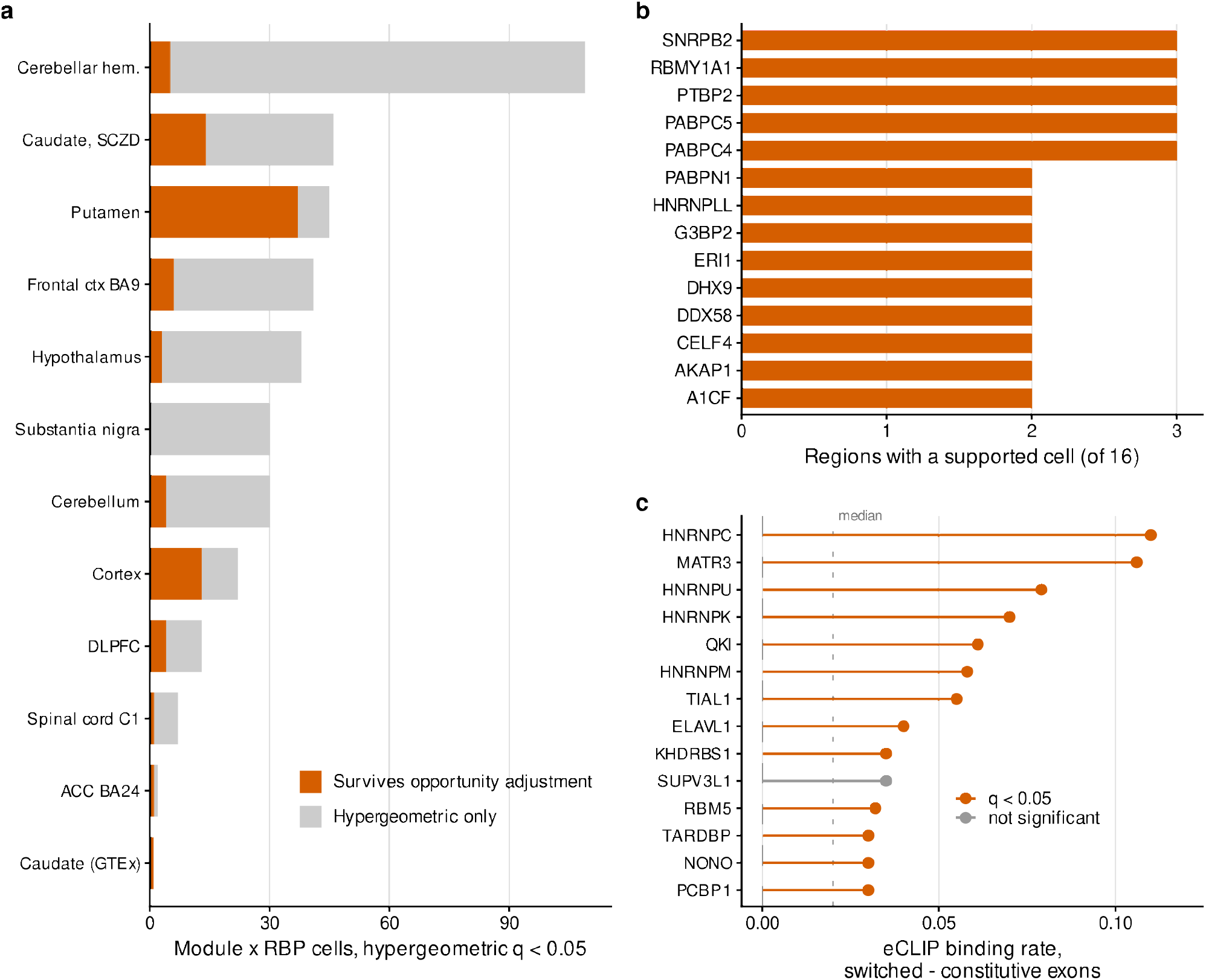
Candidate RNA-binding-protein regulons of the switch modules. **a**. Per analysis, module-by-RNA-binding protein (RBP) cells enriched for RBP motif-site switches by the hypergeometric test (Benjamini–Hochberg *q* < 0.05), split by whether they also survive a binomial model adjusting motif opportunity for transcript length, GC content, untranslated-region (UTR) and coding composition, and transcript count (adjusted *q* < 0.05, odds ratio > 1; orange). Analyses with no enriched cell are not shown. **b**. RBPs whose supported cells recur across analyses. **c**. ENCODE enhanced crosslinking and immunoprecipitation (eCLIP; HepG2 and K562 cell lines) binding at switched versus constitutive exon intervals of each RBP’s nominated genes, shown as the difference in binding rate for the 14 RBPs with the largest difference. Orange marks false discovery rate (FDR) ≤ 0.05 by an exact McNemar test per RBP, and the dotted line marks the median over all 35 testable RBPs. eCLIP indicates binding capacity in cell lines, not regulation in brain.

**Fig. S21.**
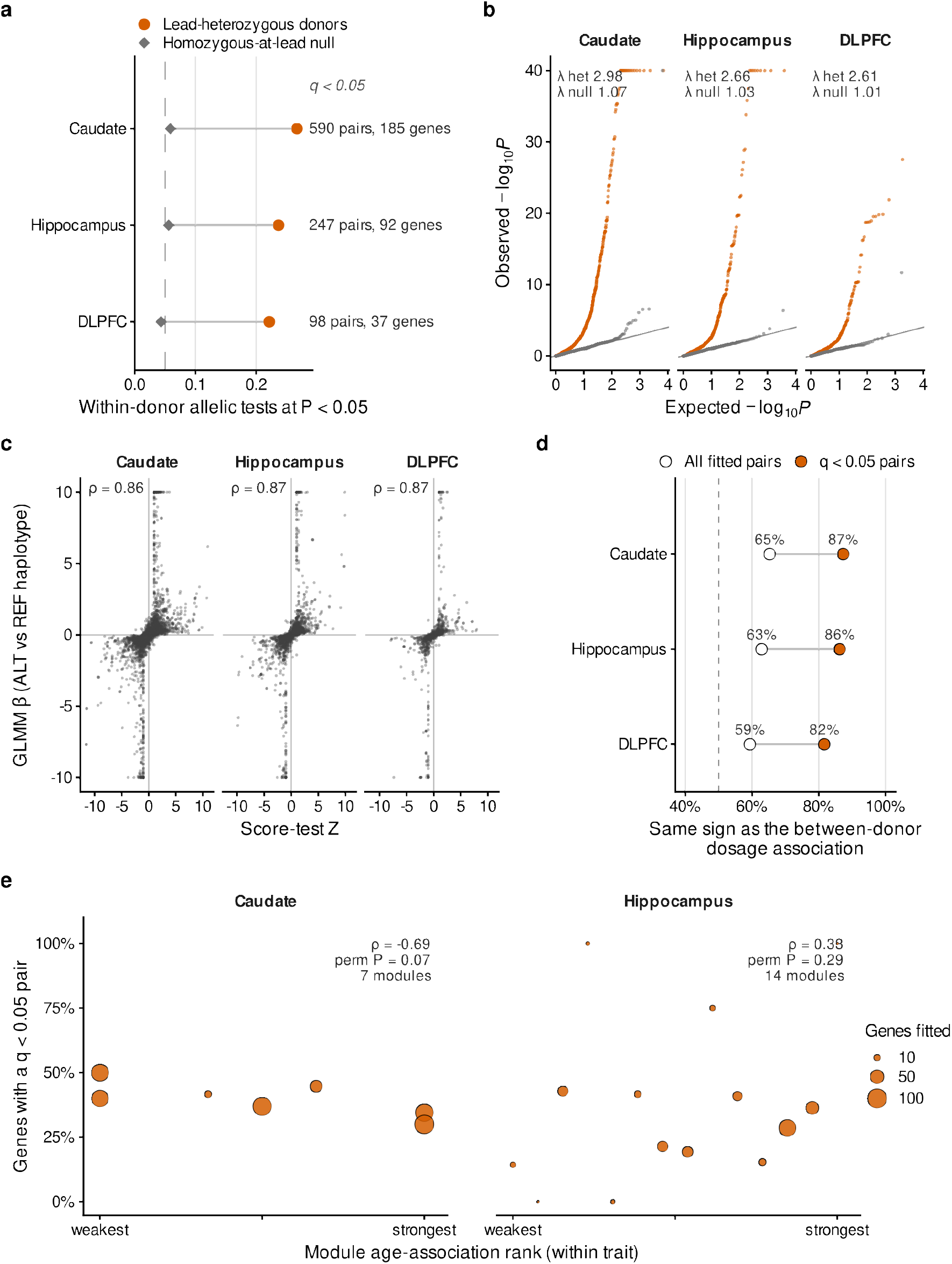
Calibration and agreement of the within-donor allelic test. BrainSEQ allele-aware junction recount. For each switch pair, fragments crossing isoform-specific junctions were counted on the two haplotypes of donors heterozygous at the switch quantitative trait locus (switch-QTL) lead and compared with a beta-binomial mixed model with a random donor intercept, which absorbs each donor’s baseline isoform ratio and cell composition. **a**. As **Fig. 5a**. **b**. Quantile–quantile plots of all fitted pairs for the lead-heterozygous test (orange) and for the same model fitted to donors homozygous at the lead (gray), with genomic inflation factors. Observed −log10 *P* is capped at 40 for display. **c**. Mixed-model allelic effect (β, alternate [ALT] against reference [REF] haplotype) against the model-free stratified score-test *Z*, with Spearman ρ. Fits at |β| = 10 are quasi-separated (one haplotype carries only one isoform): their sign is informative but their magnitude is not. The 160 pairs without a defined score statistic are not drawn. **d**. Fraction of pairs whose within-donor effect has the same sign as the between-donor association of lead dosage with isoform fraction, estimated from the same reads, over all fitted pairs (white) and pairs at Benjamini–Hochberg *q* < 0.05 (orange); the dashed line marks 50%. **e**. Per IsoGraph module, the fraction of fitted genes with a pair at *q* < 0.05 against the module’s age-association rank within its trait; point size gives the genes fitted. ρ is the Spearman correlation, and the permutation *P* comes from 10,000 permutations of gene-to-module labels that hold module sizes and the number of *cis*-controlled genes (genes with a pair at *q* < 0.05) fixed. Dorsolateral prefrontal cortex (DLPFC) had too few modules per trait to rank. With 7 and 14 modules this panel is descriptive. Pair counts are not independent: pairs of one gene share donors and fragments.

**Fig. S22.**
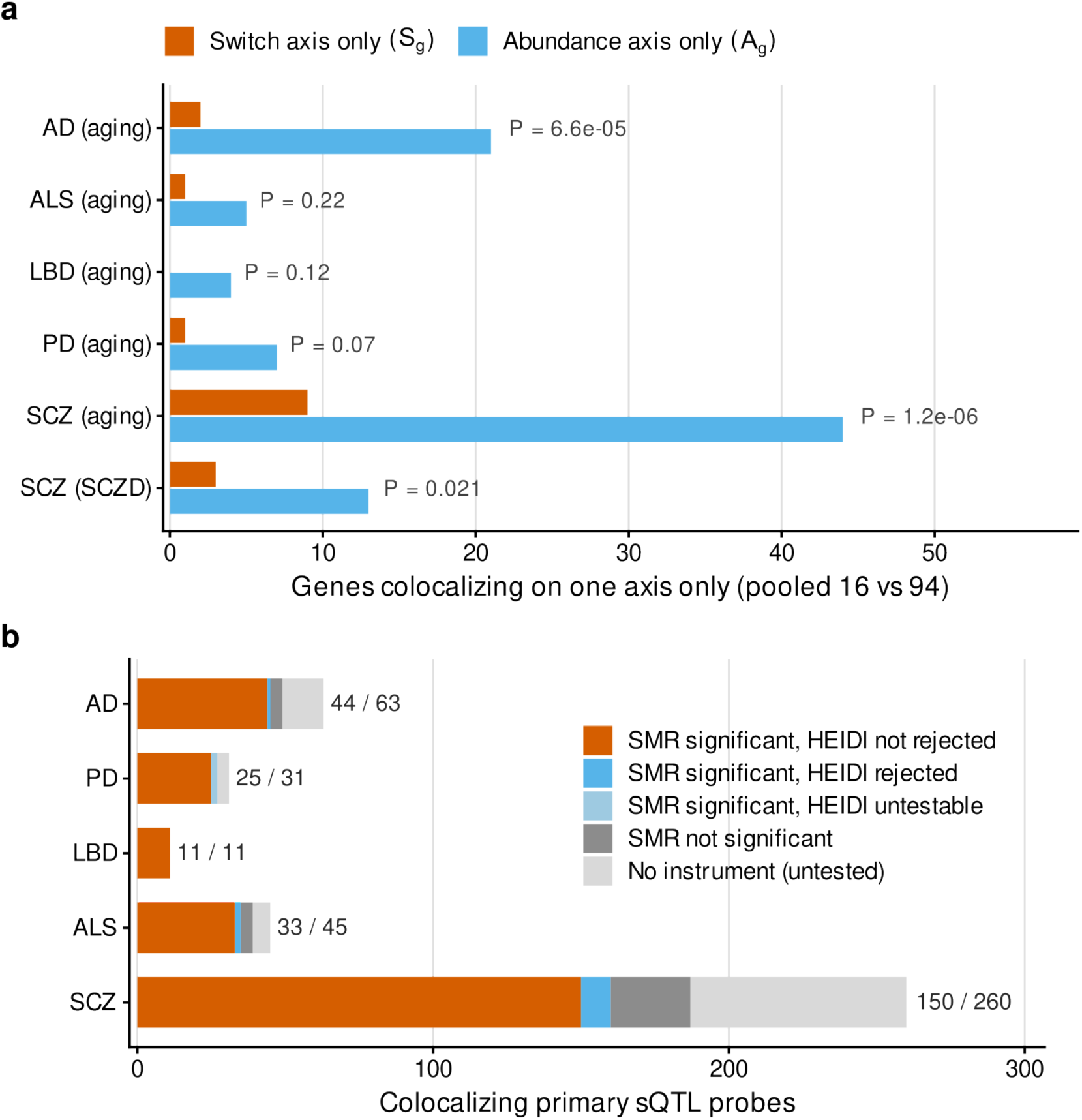
Corroboration of the colocalization layer. **a**. In European-ancestry BrainSEQ donors, genes that colocalize with each genome-wide association study (GWAS) on the IsoGraph switch axis only (orange) or the abundance axis only (blue), with exact McNemar *P*. BrainSEQ is the discovery cohort, so this is same-cohort anchoring rather than replication. **b**. Summary-data-based Mendelian randomization (SMR) and heterogeneity in dependent instruments (HEIDI) tests for the primary confirmatory probe of each colocalizing splicing quantitative trait locus (sQTL) gene, by trait. Bars count probes that SMR supports with HEIDI not rejecting a single shared variant (orange), probes that HEIDI rejects or cannot test (blue), probes that SMR leaves non-significant (dark gray) and probes without an instrument at QTL *P* < 5 × 10^−8^ (light gray), which are untested rather than negative. Labels give supported over all probes. HEIDI is underpowered at Genotype-Tissue Expression (GTEx) brain sample sizes, so non-rejection is not evidence of a shared variant.

**Fig. S23.**
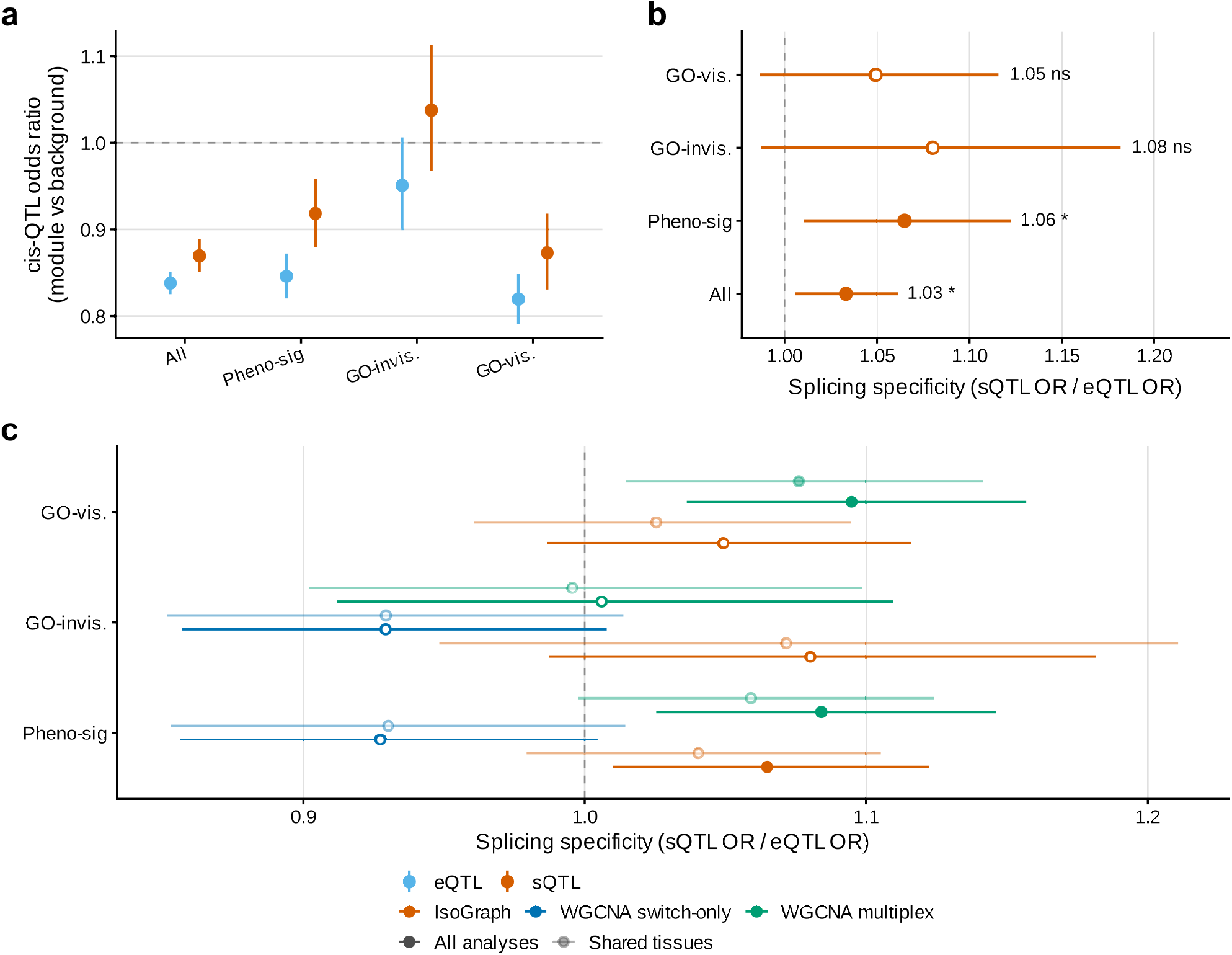
Splicing-QTL specificity of module genes. **a**. Pooled odds ratios (fixed-effect inverse-variance meta-analysis across analyses) for *cis*-expression quantitative trait locus (eQTL; blue) and *cis*-splicing QTL (sQTL; orange) status of IsoGraph module genes relative to the tested background, by module set: all modules (17 analyses), phenotype-associated modules (10), Gene Ontology (GO)-invisible modules (7), and GO-visible modules (9). Both QTL classes are depleted. **b**. The paired within-analysis ratio of the sQTL to the eQTL odds ratio, the primary estimand; bars are 95% confidence intervals and filled points have *P* < 0.05. **c**. The same ratio for IsoGraph (orange), weighted gene co-expression network analysis (WGCNA) on the same switch-only features (blue), and WGCNA on the same switch-plus-abundance features (green), over all analyses (dark) and over the analyses shared by all three methods (light). The contrast is present for both methods that consume the paired features and absent for the switch-only representation. Logistic models adjusted for *cis*-variant count, gene length, retained isoform count and intron-group size (**Tables S6** and **S7**).

**Fig. S24.**
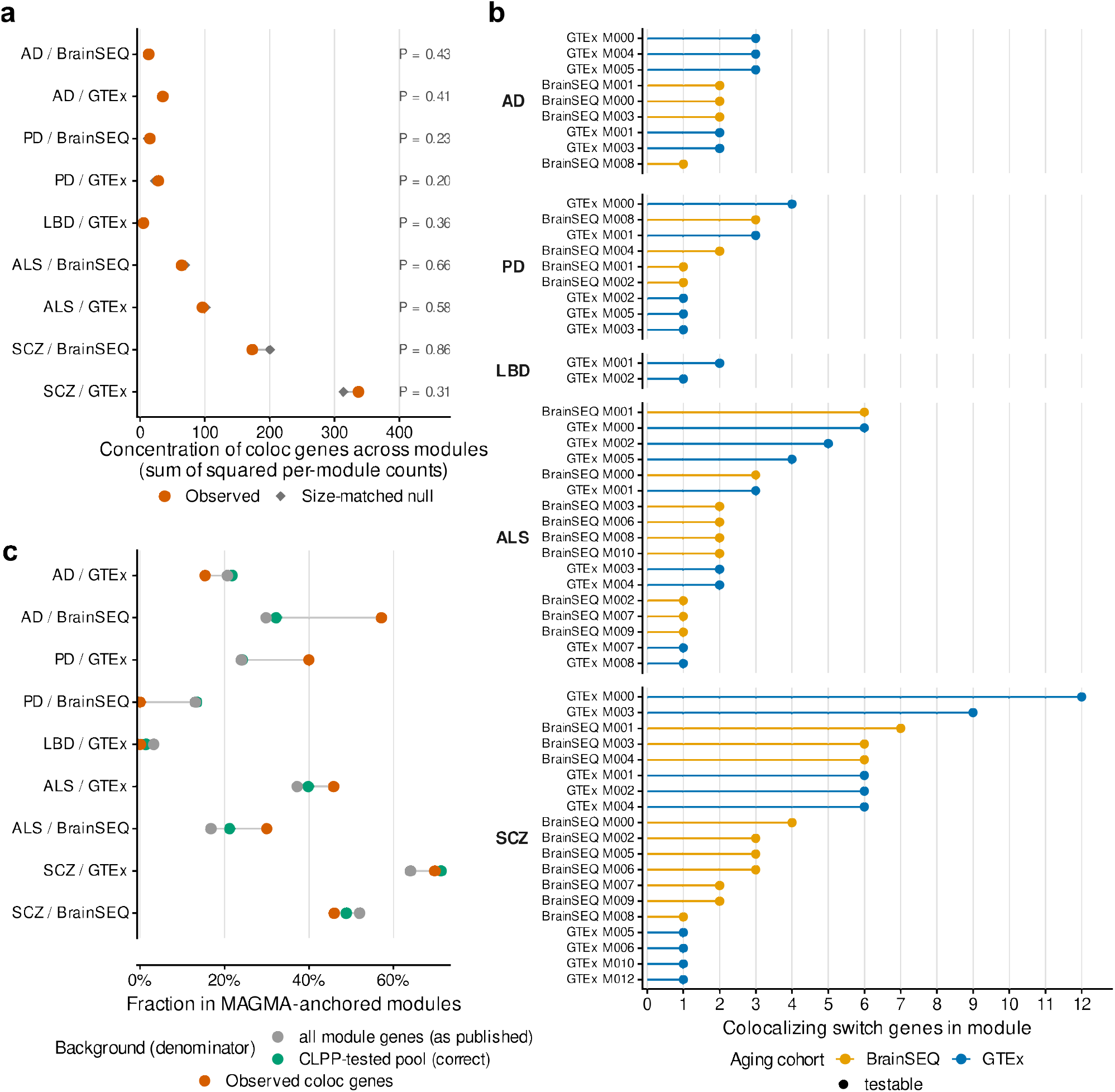
Posterior-overlap-supported genes show no detectable concentration in IsoGraph modules. **a**. For each trait and aging cohort, the concentration of posterior-overlap-supported module-member genes across modules (sum of squared per-module counts; orange) against a null that redraws the same number of genes from the overlap-score-tested pool with module sizes fixed (gray, null mean). *P* values are empirical. Dementia with Lewy bodies (LBD) had no overlap-supported genes in BrainSEQ and is not tested. **b**. Overlap-supported genes per module, by trait and cohort (BrainSEQ, yellow; Genotype-Tissue Expression [GTEx], blue). **c**. Fraction of overlap-supported genes that fall in MAGMA-anchored modules (modules with a competitive gene-set *P* < 0.05; orange), against two backgrounds: all module genes (gray) and the overlap-score-tested pool (green). Genes enter the overlap-score screen partly through the same genome-wide association study (GWAS) that defines MAGMA anchoring, so the tested pool is already enriched for anchored modules and is the correct denominator. Against it, no trait shows enrichment.

**Fig. S25.**
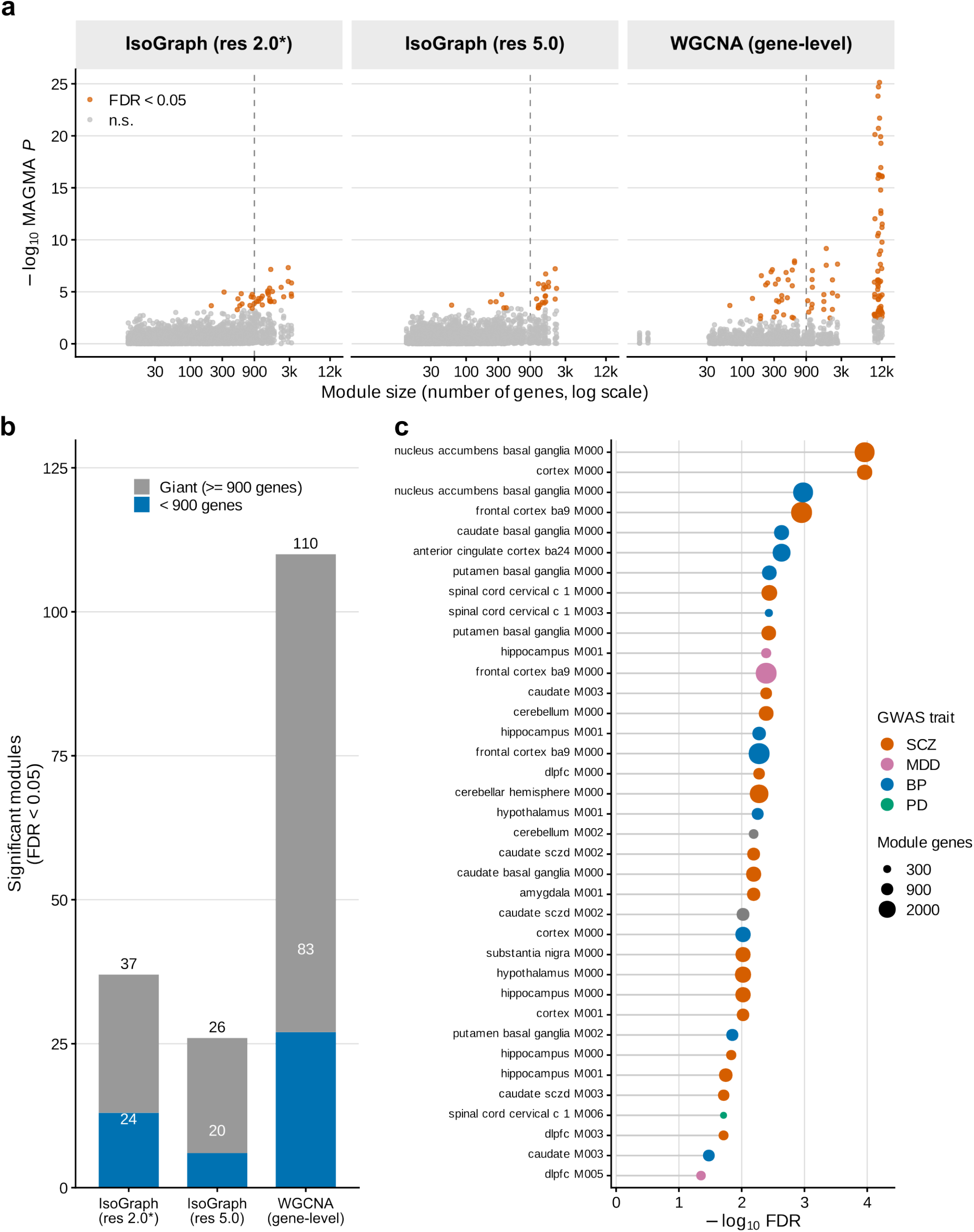
Significant module-level GWAS associations are concentrated among large modules. **a**. MAGMA competitive gene-set −log_10_ *P* against module size for IsoGraph modules at the production Leiden resolution 2.0, the resolution-5.0 sensitivity arm, and classical abundance-only weighted gene co-expression network analysis (WGCNA), over eight genome-wide association study (GWAS) traits; orange marks false discovery rate (FDR) < 0.05 and the dashed line the 900-gene cutoff. **b**. FDR-significant module-by-trait sets split by size: 24 of 37 IsoGraph sets at resolution 2.0, 20 of 26 at resolution 5.0, and 83 of 110 WGCNA sets fall in modules of at least 900 genes. **c**. The 37 FDR-significant IsoGraph sets at resolution 2.0 by −log_10_ FDR, colored by trait (gray, amyotrophic lateral sclerosis [ALS]) and sized by module genes. Module identifiers are specific to this partition (**Data S24**).

### Supplementary Tables

**Table S1.** Synthetic benchmark performance across scenarios and methods. Performance metrics are reported as means with 95% confidence intervals (CIs) in brackets. Runtime is reported as the median in seconds. *N* is the number of benchmark runs summarized in each row. Module recovery is the best-match Jaccard similarity between planted and inferred modules (1 = perfect recovery); switch detection is the recall of planted switching genes; the non-switching gene module rate is the fraction of non-switching genes assigned to modules, and lower values indicate better performance. Methods are the IsoGraph backends (Baseline, Latent, Graph, and VAE), a Spearman-correlation network with Leiden clustering (Spearman-Leiden), and WGCNA. ARI, adjusted Rand index; CPU, central processing unit; NA, unavailable; VAE, variational autoencoder; WGCNA, weighted gene co-expression network analysis.

| Scenario | Method | Compute | *N* | Module recovery, mean (95% CI) | ARI (planted), mean (95% CI) | ARI (assigned genes only), mean (95% CI) | Homogeneity, mean (95% CI) | Completeness, mean (95% CI) | Switch detection, mean (95% CI) | Non-switching gene module rate (lower is better), mean (95% CI) | Runtime (s), median |
| --- | --- | --- | --- | --- | --- | --- | --- | --- | --- | --- | --- |
| Idealized switching | IsoGraph Baseline | CPU | 360 | 0.074<br>[0.070, 0.077] | 0.001<br>[0.001, 0.002] | 0.001<br>[0.001, 0.001] | 0.118<br>[0.100, 0.137] | 0.684<br>[0.650, 0.719] | 0.898<br>[0.881, 0.914] | 0.297<br>[0.282, 0.311] | 0.7 |
| Idealized switching | IsoGraph Latent | CPU | 360 | 0.217<br>[0.183, 0.253] | 0.468<br>[0.408, 0.530] | 0.996<br>[0.991, 1.000] | 0.999<br>[0.998, 1.000] | 0.683<br>[0.650, 0.717] | 0.217<br>[0.182, 0.253] | 0.000<br>[0.000, 0.000] | 5.7 |
| Idealized switching | IsoGraph Graph | CPU | 360 | 0.300<br>[0.265, 0.336] | 0.590<br>[0.555, 0.626] | 0.995<br>[0.989, 1.000] | 0.999<br>[0.998, 1.000] | 0.706<br>[0.683, 0.730] | 0.300<br>[0.265, 0.336] | 0.000<br>[0.000, 0.000] | 6.5 |
| Idealized switching | IsoGraph VAE | CPU | 360 | 0.968<br>[0.961, 0.974] | 0.998<br>[0.996, 1.000] | 0.998<br>[0.996, 1.000] | 0.999<br>[0.998, 1.000] | 1.000<br>[1.000, 1.000] | 1.000<br>[1.000, 1.000] | 0.021<br>[0.018, 0.024] | 4.1 |
| Idealized switching | Spearman-Leiden | CPU | 360 | 0.632<br>[0.628, 0.637] | 0.615<br>[0.611, 0.620] | 0.630<br>[0.624, 0.636] | 0.989<br>[0.987, 0.991] | 0.736<br>[0.733, 0.739] | 0.985<br>[0.982, 0.988] | 0.089<br>[0.086, 0.092] | 1 |
| Idealized switching | WGCNA | CPU | 360 | 0.572<br>[0.557, 0.586] | 0.649<br>[0.645, 0.653] | 0.669<br>[0.663, 0.674] | 0.991<br>[0.988, 0.994] | 0.766<br>[0.764, 0.768] | 0.980<br>[0.976, 0.984] | 0.232<br>[0.192, 0.274] | 3.4 |
| Noise stress | IsoGraph Baseline | CPU | 320 | 0.057<br>[0.056, 0.057] | 0.000<br>[-0.000, 0.000] | 0.000<br>[0.000, 0.000] | 0.000<br>[0.000, 0.000] | 0.997<br>[0.991, 1.000] | 1.000<br>[1.000, 1.000] | 0.419<br>[0.410, 0.428] | 0.7 |
| Noise stress | IsoGraph Latent | CPU | 320 | 0.113<br>[0.101, 0.126] | 0.078<br>[0.068, 0.088] | 1.000<br>[1.000, 1.000] | 1.000<br>[1.000, 1.000] | 0.473<br>[0.471, 0.476] | 0.111<br>[0.099, 0.123] | 0.000<br>[0.000, 0.000] | 6.3 |
| Noise stress | IsoGraph Graph | CPU | 320 | 0.500<br>[0.490, 0.511] | 0.421<br>[0.411, 0.430] | 1.000<br>[1.000, 1.000] | 1.000<br>[1.000, 1.000] | 0.580<br>[0.576, 0.584] | 0.497<br>[0.487, 0.508] | 0.000<br>[0.000, 0.000] | 6.3 |
| Noise stress | IsoGraph VAE | CPU | 320 | 0.974<br>[0.971, 0.977] | 1.000<br>[0.999, 1.000] | 1.000<br>[0.999, 1.000] | 1.000<br>[0.999, 1.000] | 1.000<br>[1.000, 1.000] | 1.000<br>[1.000, 1.000] | 0.020<br>[0.018, 0.022] | 4.1 |
| Noise stress | Spearman-Leiden | CPU | 320 | 0.636<br>[0.631, 0.641] | 0.614<br>[0.609, 0.618] | 0.615<br>[0.611, 0.620] | 0.991<br>[0.989, 0.993] | 0.732<br>[0.730, 0.734] | 0.998<br>[0.998, 0.999] | 0.092<br>[0.089, 0.094] | 1 |
| Noise stress | WGCNA | CPU | 320 | 0.629<br>[0.619, 0.638] | 0.663<br>[0.659, 0.666] | 0.665<br>[0.662, 0.669] | 1.000<br>[1.000, 1.000] | 0.768<br>[0.767, 0.769] | 0.997<br>[0.997, 0.998] | 0.046<br>[0.027, 0.067] | 3.4 |
| Feature interactions | IsoGraph Baseline | CPU | 300 | 0.052<br>[0.051, 0.053] | 0.000<br>[0.000, 0.000] | 0.000<br>[0.000, 0.000] | 0.000<br>[0.000, 0.000] | 1.000<br>[1.000, 1.000] | 1.000<br>[1.000, 1.000] | 0.489<br>[0.476, 0.503] | 0.7 |
| Feature interactions | IsoGraph Latent | CPU | 300 | 0.326<br>[0.312, 0.340] | 0.238<br>[0.225, 0.250] | 0.766<br>[0.750, 0.782] | 1.000<br>[1.000, 1.000] | 0.532<br>[0.528, 0.537] | 0.437<br>[0.416, 0.457] | 0.000<br>[0.000, 0.000] | 6.2 |
| Feature interactions | IsoGraph Graph | CPU | 300 | 0.482<br>[0.473, 0.491] | 0.385<br>[0.375, 0.396] | 0.856<br>[0.844, 0.868] | 1.000<br>[1.000, 1.000] | 0.580<br>[0.576, 0.584] | 0.565<br>[0.553, 0.576] | 0.000<br>[0.000, 0.000] | 6.8 |
| Feature interactions | IsoGraph VAE | CPU | 300 | 0.769<br>[0.739, 0.799] | 0.670<br>[0.626, 0.715] | 0.671<br>[0.626, 0.715] | 0.786<br>[0.755, 0.817] | 0.847<br>[0.825, 0.869] | 0.999<br>[0.999, 0.999] | 0.009<br>[0.008, 0.010] | 5.3 |
| Feature interactions | Spearman-Leiden | CPU | 300 | 0.449<br>[0.428, 0.469] | 0.335<br>[0.307, 0.363] | 0.336<br>[0.308, 0.364] | 0.628<br>[0.590, 0.665] | 0.560<br>[0.539, 0.581] | 0.998<br>[0.998, 0.999] | 0.089<br>[0.087, 0.092] | 0.9 |
| Feature interactions | WGCNA | CPU | 300 | 0.476<br>[0.456, 0.496] | 0.425<br>[0.397, 0.453] | 0.427<br>[0.400, 0.455] | 0.795<br>[0.761, 0.829] | 0.614<br>[0.593, 0.636] | 0.990<br>[0.988, 0.992] | 0.088<br>[0.058, 0.118] | 3.1 |
| Non-switching background | IsoGraph Baseline | CPU | 80 | 0.054<br>[0.047, 0.062] | 0.002<br>[0.000, 0.004] | 0.002<br>[0.000, 0.004] | 0.018<br>[0.011, 0.027] | 0.835<br>[0.769, 0.893] | 0.990<br>[0.986, 0.994] | 0.342<br>[0.325, 0.359] | 0.7 |
| Non-switching background | IsoGraph Latent | CPU | 80 | 0.312<br>[0.237, 0.393] | 0.481<br>[0.388, 0.571] | 0.890<br>[0.829, 0.945] | 0.972<br>[0.954, 0.987] | 0.727<br>[0.677, 0.776] | 0.346<br>[0.269, 0.426] | 0.000<br>[0.000, 0.000] | 6.2 |
| Non-switching background | IsoGraph Graph | CPU | 80 | 0.382<br>[0.314, 0.453] | 0.555<br>[0.493, 0.621] | 0.865<br>[0.787, 0.935] | 0.965<br>[0.941, 0.985] | 0.736<br>[0.703, 0.771] | 0.420<br>[0.354, 0.487] | 0.000<br>[0.000, 0.000] | 6.9 |
| Non-switching background | IsoGraph VAE | CPU | 80 | 0.841<br>[0.778, 0.898] | 0.911<br>[0.849, 0.964] | 0.911<br>[0.849, 0.964] | 0.924<br>[0.867, 0.972] | 1.000<br>[1.000, 1.000] | 1.000<br>[1.000, 1.000] | 0.109<br>[0.067, 0.157] | 4.6 |
| Non-switching background | Spearman-Leiden | CPU | 80 | 0.629<br>[0.609, 0.649] | 0.609<br>[0.592, 0.628] | 0.698<br>[0.667, 0.731] | 0.990<br>[0.985, 0.994] | 0.761<br>[0.750, 0.772] | 0.905<br>[0.870, 0.937] | 0.093<br>[0.088, 0.099] | 0.9 |
| Non-switching background | WGCNA | CPU | 80 | 0.535<br>[0.499, 0.570] | 0.633<br>[0.611, 0.655] | 0.650<br>[0.626, 0.673] | 0.985<br>[0.976, 0.993] | 0.784<br>[0.776, 0.793] | 0.983<br>[0.976, 0.990] | 0.342<br>[0.242, 0.442] | 4.2 |
| Unequal abundance | IsoGraph Baseline | CPU | 80 | 0.082<br>[0.071, 0.095] | 0.013<br>[0.006, 0.022] | 0.220<br>[0.141, 0.306] | 0.274<br>[0.184, 0.372] | 0.748<br>[0.687, 0.809] | 0.766<br>[0.675, 0.848] | 0.240<br>[0.201, 0.278] | 0.6 |
| Unequal abundance | IsoGraph Latent | CPU | 80 | 0.152<br>[0.131, 0.173] | 0.086<br>[0.071, 0.102] | 1.000<br>[1.000, 1.000] | 1.000<br>[1.000, 1.000] | 0.476<br>[0.472, 0.480] | 0.148<br>[0.128, 0.170] | 0.000<br>[0.000, 0.000] | 5.8 |
| Unequal abundance | IsoGraph Graph | CPU | 80 | 0.428<br>[0.395, 0.458] | 0.336<br>[0.307, 0.364] | 1.000<br>[1.000, 1.000] | 1.000<br>[1.000, 1.000] | 0.554<br>[0.544, 0.564] | 0.425<br>[0.393, 0.456] | 0.000<br>[0.000, 0.000] | 5.7 |
| Unequal abundance | IsoGraph VAE | CPU | 80 | 0.780<br>[0.707, 0.849] | 0.818<br>[0.738, 0.894] | 0.818<br>[0.738, 0.894] | 0.833<br>[0.755, 0.904] | 0.999<br>[0.998, 1.000] | 1.000<br>[1.000, 1.000] | 0.151<br>[0.107, 0.198] | 4 |
| Unequal abundance | Spearman-Leiden | CPU | 80 | 0.620<br>[0.611, 0.628] | 0.604<br>[0.596, 0.611] | 0.606<br>[0.598, 0.614] | 0.990<br>[0.986, 0.993] | 0.730<br>[0.726, 0.734] | 0.998<br>[0.997, 0.999] | 0.096<br>[0.091, 0.101] | 0.9 |
| Unequal abundance | WGCNA | CPU | 80 | 0.553<br>[0.524, 0.580] | 0.600<br>[0.579, 0.620] | 0.605<br>[0.584, 0.624] | 0.999<br>[0.998, 1.000] | 0.737<br>[0.726, 0.747] | 0.994<br>[0.991, 0.997] | 0.093<br>[0.050, 0.141] | 4 |
| Negative control | IsoGraph Baseline | CPU | 40 | 0.000 [0.000, 0.000] | NA [NA, NA] | NA [NA, NA] | NA [NA, NA] | NA [NA, NA] | 0.000 [0.000, 0.000] | 0.000 [0.000, 0.000] | 1.4 |
| Negative control | IsoGraph Latent | CPU | 40 | 0.000 [0.000, 0.000] | NA [NA, NA] | NA [NA, NA] | NA [NA, NA] | NA [NA, NA] | 0.000 [0.000, 0.000] | 0.000 [0.000, 0.000] | 25.8 |
| Negative control | IsoGraph Graph | CPU | 40 | 0.000 [0.000, 0.000] | NA [NA, NA] | NA [NA, NA] | NA [NA, NA] | NA [NA, NA] | 0.000 [0.000, 0.000] | 0.000 [0.000, 0.000] | 21 |
| Negative control | IsoGraph VAE | CPU | 40 | 0.054 [0.017, 0.100] | NA [NA, NA] | NA [NA, NA] | NA [NA, NA] | NA [NA, NA] | 0.925 [0.825, 1.000] | 0.790 [0.687, 0.884] | 4.7 |
| Negative control | Spearman-Leiden | CPU | 40 | 0.013 [0.000, 0.037] | NA [NA, NA] | NA [NA, NA] | NA [NA, NA] | NA [NA, NA] | 0.025 [0.000, 0.075] | 0.095 [0.089, 0.101] | 1 |
| Negative control | WGCNA | CPU | 40 | 0.405 [0.359, 0.446] | NA [NA, NA] | NA [NA, NA] | NA [NA, NA] | NA [NA, NA] | 1.000 [1.000, 1.000] | 1.000 [1.000, 1.000] | 24.3 |

**Table S2.** Runtime and memory use across network sizes and compute settings. Runtime, peak host RAM, and peak reserved GPU VRAM are reported as medians with interquartile ranges (IQRs) in brackets. *N* is the number of benchmark runs summarized in each row. Runtime is measured in seconds and memory in gigabytes (GB). NA indicates that GPU VRAM measurements are not applicable to CPU runs. CPU, central processing unit; GPU, graphics processing unit; RAM, random-access memory; VAE, variational autoencoder; VRAM, video RAM; WGCNA, weighted gene co-expression network analysis.

| Genes (*n*) | Method | Compute | *N* | Runtime (s), median [IQR] | Peak host RAM (GB), median [IQR] | Peak reserved GPU VRAM (GB), median [IQR] |
| --- | --- | --- | --- | --- | --- | --- |
| 1000 | IsoGraph VAE | CPU | 30 | 6.3 [5.8, 7.4] | 0.9 [0.8, 1.0] | NA |
| 1000 | IsoGraph VAE GPU | GPU | 30 | 4.1 [3.8, 4.5] | 1.2 [1.2, 1.2] | 0.0 [0.0, 0.1] |
| 1000 | WGCNA | CPU | 30 | 6.7 [4.7, 8.1] | 0.7 [0.6, 0.7] | NA |
| 3000 | IsoGraph VAE | CPU | 30 | 28.2 [18.0, 35.0] | 1.1 [1.1, 1.2] | NA |
| 3000 | IsoGraph VAE GPU | GPU | 30 | 7.9 [7.3, 8.6] | 1.4 [1.3, 1.5] | 0.2 [0.1, 0.3] |
| 3000 | WGCNA | CPU | 30 | 17.8 [11.3, 26.4] | 1.0 [1.0, 1.8] | NA |
| 6000 | IsoGraph VAE | CPU | 30 | 36.7 [29.3, 84.0] | 1.6 [1.6, 1.8] | NA |
| 6000 | IsoGraph VAE GPU | GPU | 30 | 11.6 [11.2, 13.1] | 1.7 [1.6, 2.0] | 0.2 [0.2, 0.4] |
| 6000 | WGCNA | CPU | 30 | 35.6 [28.0, 88.5] | 2.0 [2.0, 2.8] | NA |
| 12000 | IsoGraph VAE | CPU | 30 | 79.7 [52.3, 153.7] | 3.0 [2.8, 4.2] | NA |
| 12000 | IsoGraph VAE GPU | GPU | 30 | 25.0 [24.3, 33.9] | 2.8 [2.7, 4.0] | 0.4 [0.4, 0.8] |
| 12000 | WGCNA | CPU | 30 | 171.3 [82.9, 246.2] | 3.1 [3.1, 3.3] | NA |
| 16000 | IsoGraph VAE | CPU | 30 | 226.3 [192.3, 246.8] | 6.6 [6.4, 6.7] | NA |
| 16000 | IsoGraph VAE GPU | GPU | 30 | 24.6 [22.2, 30.1] | 6.2 [5.9, 6.4] | 1.2 [1.0, 1.2] |
| 16000 | WGCNA | CPU | 30 | 383.5 [365.2, 409.3] | 3.7 [3.7, 3.7] | NA |

**Table S3.**
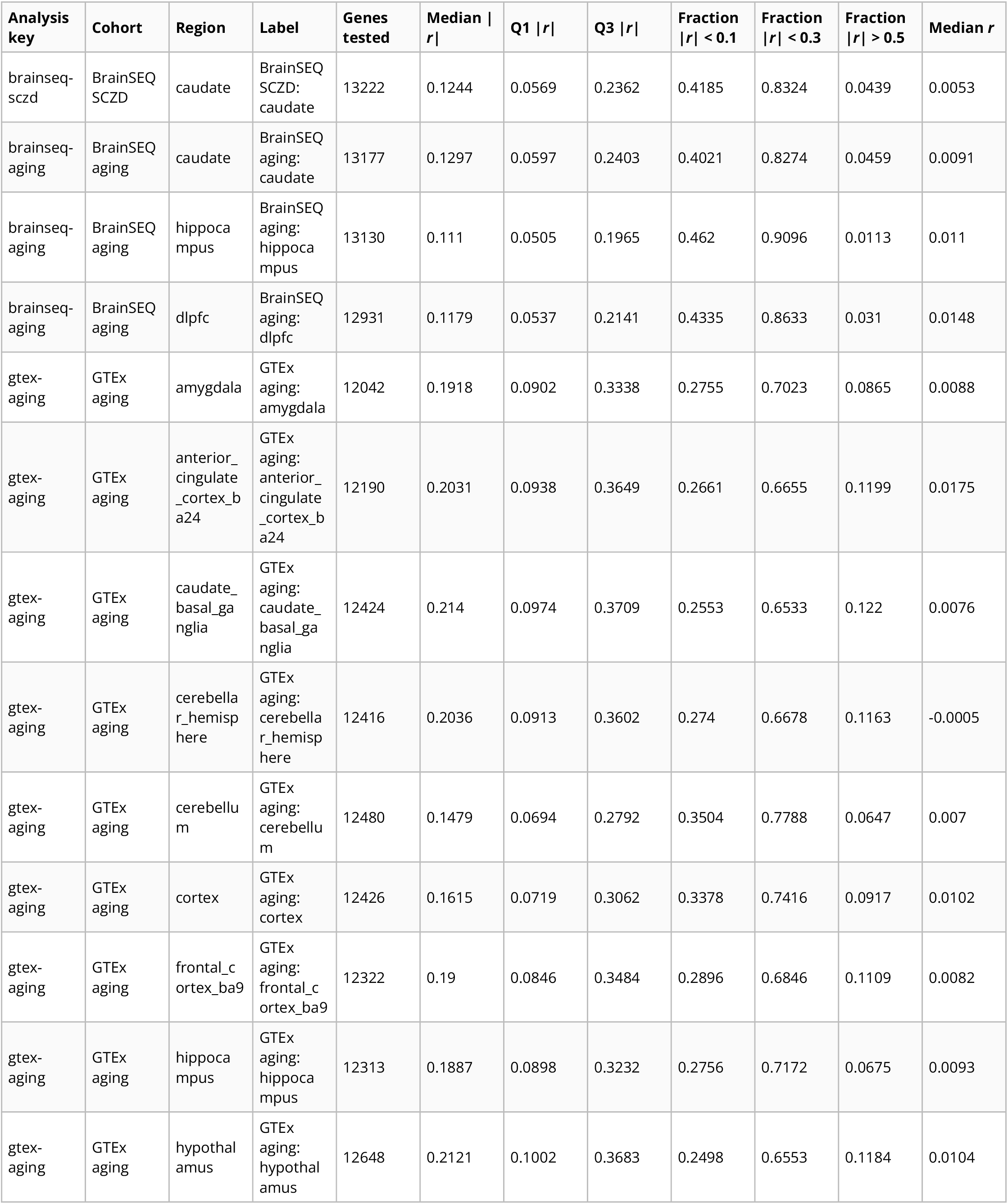
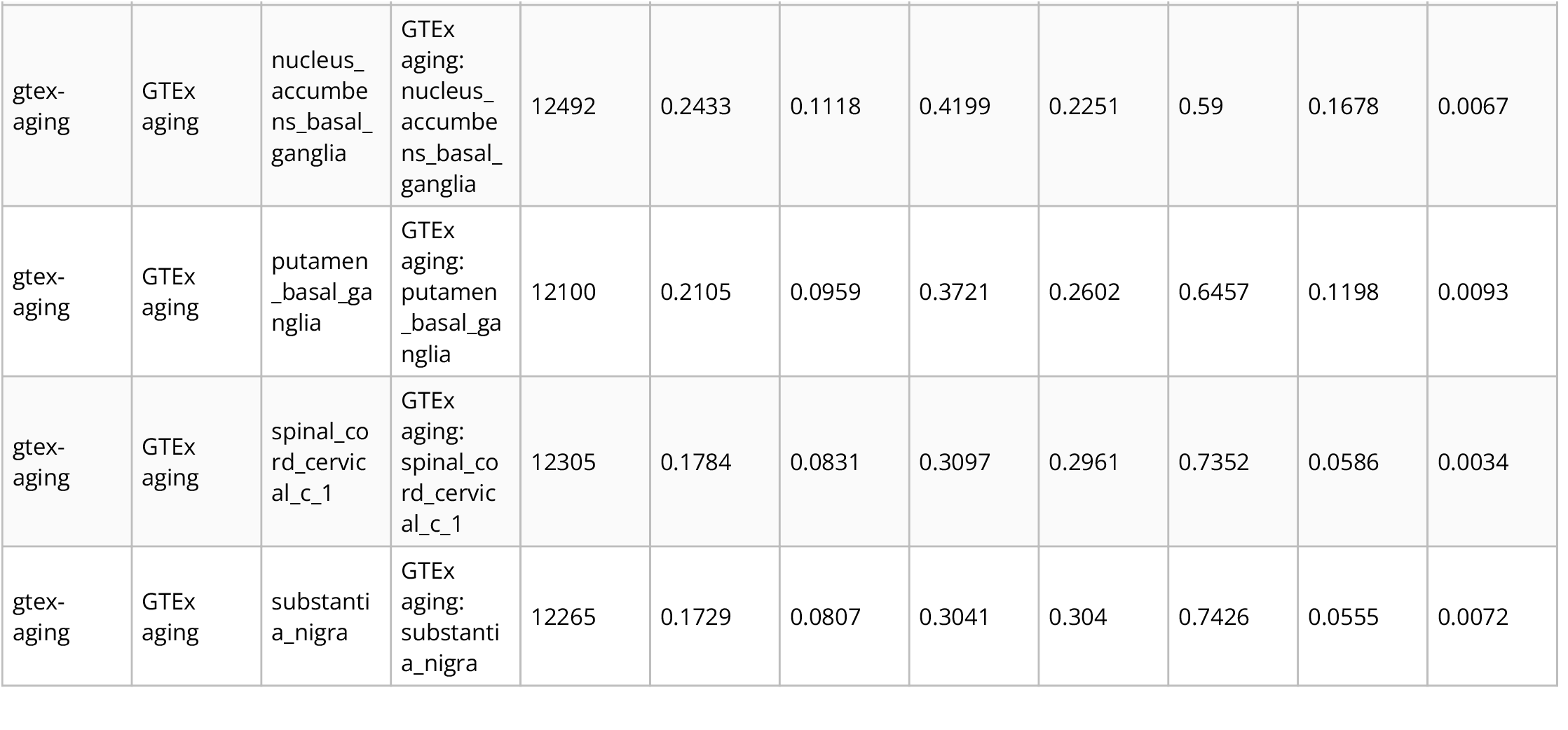
Separation of the switch and abundance coordinates, per analysis. One row per analysis (17 rows): genes tested, the median absolute Pearson correlation (|*r*|) between the gene-abundance and isoform-switch coordinates with its first and third quartiles (Q1 and Q3), the fraction of genes with |*r*| below 0.1, below 0.3, and above 0.5, and the median signed correlation. The analysis key and region are the identifiers used in the analysis code; the label combines cohort and region. Median |*r*| ranges from 0.111 in BrainSEQ aging hippocampus to 0.243 in GTEx nucleus accumbens, with 23–46% of genes below 0.1 and 1.1–16.8% above 0.5, over 12,042–13,222 genes per analysis. The quartiles and tail fractions are given because the claim concerns the bulk of the distribution rather than a central value: the two coordinates are largely distinct in every analysis, and close to orthogonal in none. This test requires no deconvolution, so unlike Data S2 it covers all 17 analyses. BrainSEQ aging and BrainSEQ SCZD (schizophrenia) are the BrainSEQ aging and schizophrenia analyses; GTEx, Genotype-Tissue Expression. Backs **Fig. 2b** and **Fig. S12a**.

**Table S4.** Pooled per-module association rates across the 17 analyses. One row per module set, pooled over all analyses: the number of analyses, the median module count and module size, the mean per-analysis fraction of modules associated with the phenotype, the mean fraction associated on both the switch and abundance coordinates, the mean fraction with an enriched Gene Ontology (GO) term, and the corresponding totals. Methods are IsoGraph (isograph) and weighted gene co-expression network analysis (WGCNA) on gene abundance (wgcna_gene), on switch-only features (wgcna_switch_only), and on the same switch and abundance features (wgcna_multiplex). The comparison is not a like-for-like ranking: the four sets partition the same samples at very different granularities (median 25 modules of 108 genes for IsoGraph against 8 modules of 468 genes for abundance WGCNA), so a per-module rate is computed over different numbers of modules in each row. Backs **Fig. S15**.

| Method | Features | Analyses (*n*) | Median modules (*n*) | Median module size | Mean fraction phenotype-associated | Mean fraction both coordinates | Mean fraction GO-enriched | Total phenotype-associated | Total both coordinates |
| --- | --- | --- | --- | --- | --- | --- | --- | --- | --- |
| isograph | switch+abundance | 17 | 25 | 108 | 0.146 | 0.052 | 0.253 | 54 | 20 |
| wgcna_gene | abundance | 17 | 8 | 468 | 0.178 | 0.159 | 0.862 | 28 | 24 |
| wgcna_switch_only | switch-only | 17 | 18 | 159 | 0.214 | 0.025 | 0.184 | 64 | 8 |
| wgcna_multiplex | switch+abundance | 17 | 25 | 242 | 0.189 | 0.117 | 0.649 | 75 | 48 |

**Table S5.**
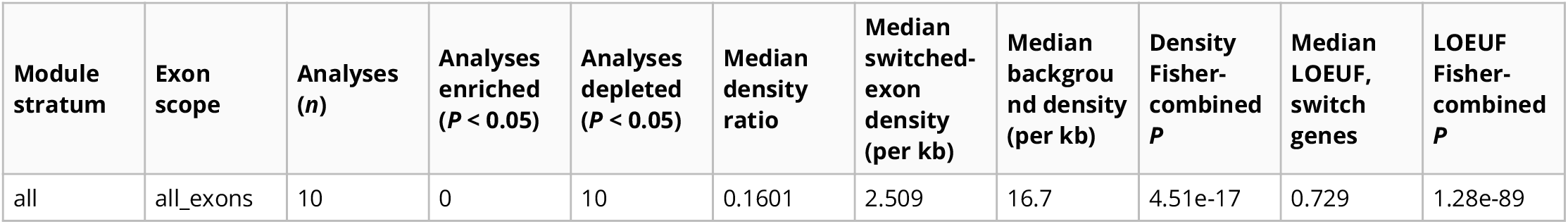
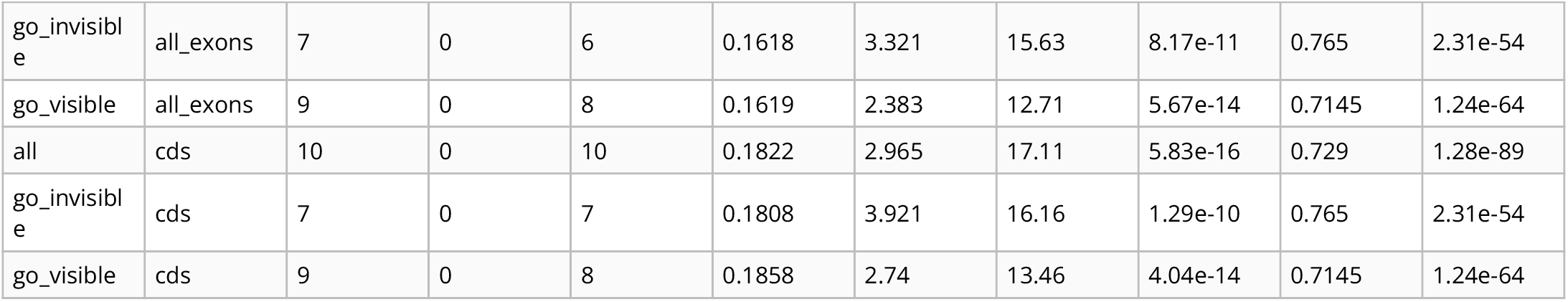
Constraint and ClinVar density of switch genes and switched exons. One row per module stratum (all switch modules [all], Gene Ontology [GO]-invisible modules [go_invisible], and GO-visible modules [go_visible]) and exon scope (all switch-pair exons [all_exons], or coding exons only [cds]). Columns give the number of annotatable analyses, how many showed switched exons enriched or depleted for ClinVar pathogenic or likely-pathogenic entries relative to other exons of the same transcripts at within-gene permutation *P* < 0.05, the median within-analysis density ratio (switched over background exons, per kilobase [kb]) with its Fisher-combined *P*, and the median gnomAD loss-of-function observed/expected upper-bound fraction (LOEUF) of switch genes with the Fisher-combined one-sided Mann–Whitney *P* against the genome-wide reference. Densities count variant call format (VCF) entries, not unique alleles. Backs **Fig. S19a,b**.

**Table S6.** Splicing-QTL specificity contrast, all analyses. One row per network method and module set: the number of analyses pooled, the fixed-effect inverse-variance pooled ratio of the splicing quantitative trait locus (sQTL) odds ratio to the expression QTL (eQTL) odds ratio for module membership (paired within analysis, which removes the shared *cis*-QTL depletion of network genes), its 95% confidence interval (CI) and *P* value, and between-analysis heterogeneity (*I*^2^). Methods are IsoGraph (isograph) and classical weighted gene co-expression network analysis (WGCNA) on the same switch-only (wgcna_switch_only) or switch-plus-abundance (wgcna_multiplex) features. Module sets are all modules, phenotype-associated modules (pheno_sig_modules), and Gene Ontology (GO)-invisible and GO-visible phenotype-associated modules. Backs **Fig. S23b,c**.

| Method | Module set | Analyses (n) | sQTL/eQTL odds-ratio ratio | 95% CI, lower | 95% CI, upper | $P$ | $I^2$ |
| --- | --- | --- | --- | --- | --- | --- | --- |
| isograph | all_modules | 17 | 1.033 | 1.006 | 1.062 | 0.0179 | 0.6036 |
| isograph | pheno_sig_modules | 10 | 1.065 | 1.01 | 1.123 | 0.0198 | 0.104 |
| wgcna_switch_only | pheno_sig_modules | 6 | 0.9273 | 0.856 | 1.005 | 0.0646 | 0.4648 |
| wgcna_multiplex | pheno_sig_modules | 8 | 1.084 | 1.025 | 1.146 | 0.00444 | 0.6386 |
| isograph | go_invisible_modules | 7 | 1.08 | 0.9873 | 1.182 | 0.0927 | 0 |
| wgcna_switch_only | go_invisible_modules | 6 | 0.9292 | 0.8567 | 1.008 | 0.0762 | 0.4589 |
| wgcna_multiplex | go_invisible_modules | 4 | 1.006 | 0.9121 | 1.109 | 0.905 | 0.6106 |
| isograph | go_visible_modules | 9 | 1.049 | 0.9866 | 1.116 | 0.126 | 0.3384 |
| wgcna_switch_only | go_visible_modules | 2 | 0.9694 | 0.7851 | 1.197 | 0.773 | 0 |
| wgcna_multiplex | go_visible_modules | 8 | 1.095 | 1.036 | 1.157 | 0.00126 | 0.5034 |

**Table S7.**
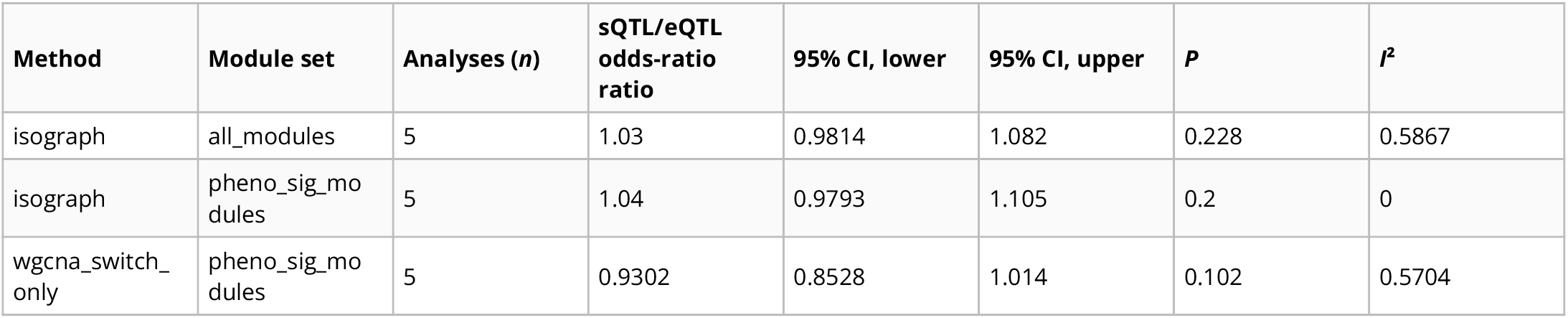
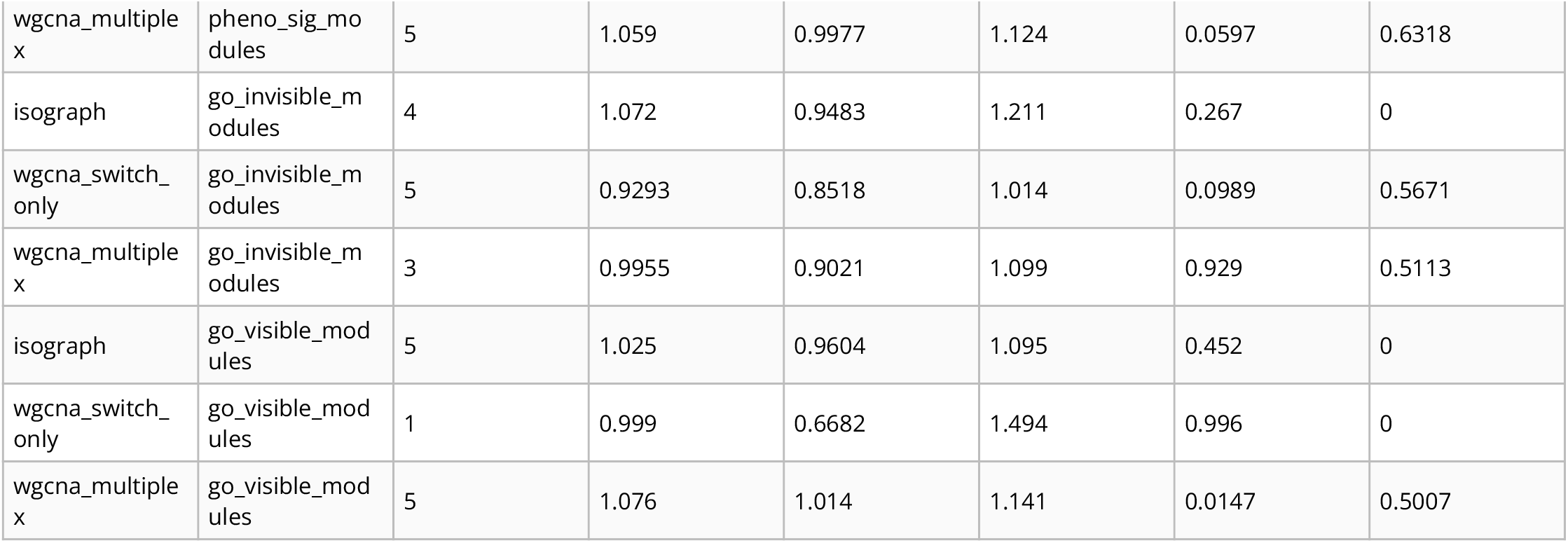
Splicing-QTL specificity contrast on the analyses shared by all methods. As Table S6, restricted to the analyses in which IsoGraph and both matched weighted gene co-expression network analysis (WGCNA) comparators produced the relevant module set, so that the three methods are compared on identical inputs. Columns give the number of analyses pooled, the fixed-effect pooled ratio of the splicing quantitative trait locus (sQTL) odds ratio to the expression QTL (eQTL) odds ratio for module membership, its 95% confidence interval (CI) and *P* value, and between-analysis heterogeneity (*I*^2^). Method and module-set labels are as in Table S6. Backs **Fig. S23c**.

**Table S8.**
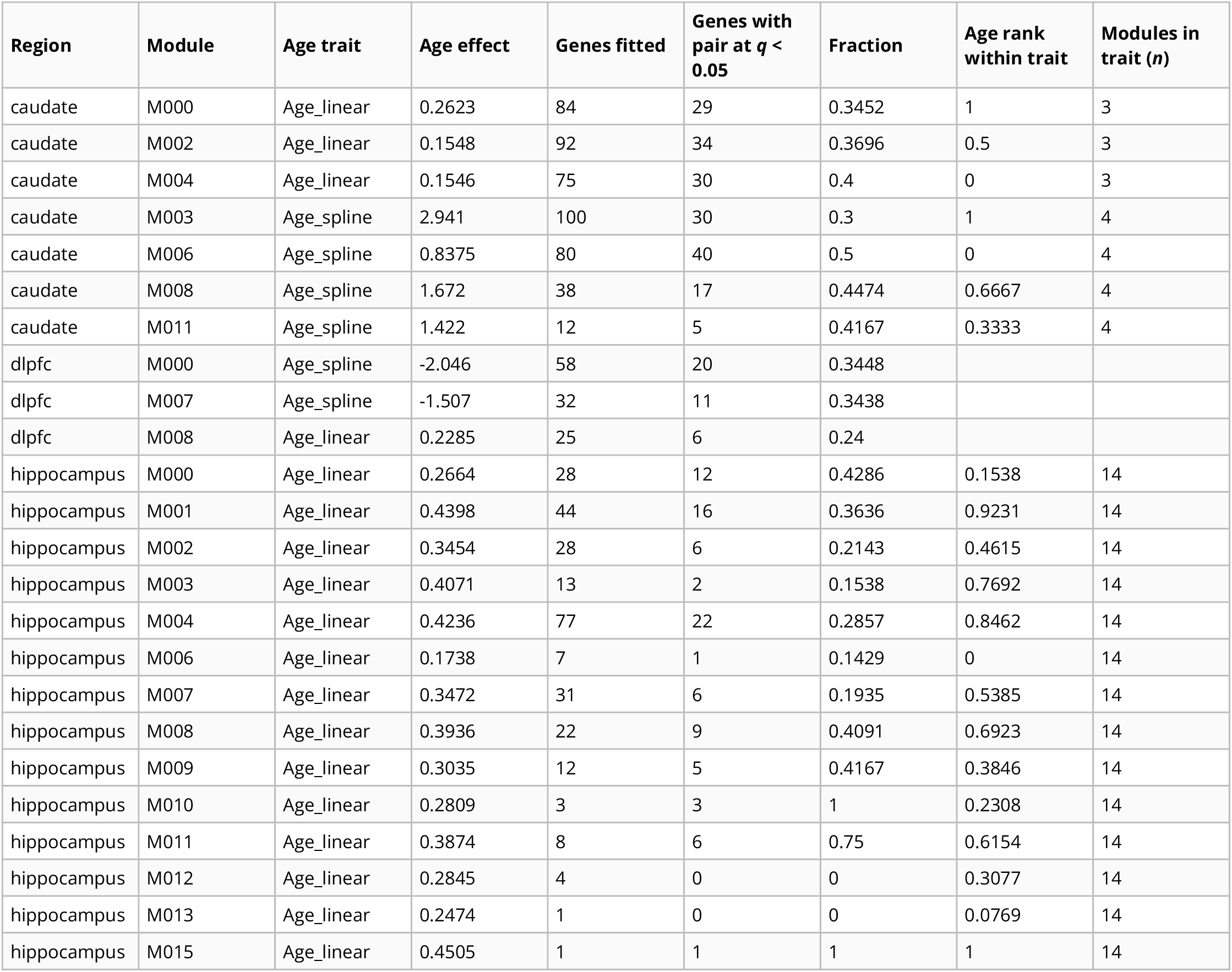
Module-level *cis* control of isoform choice. One row per age-selected IsoGraph module with a fitted gene in the BrainSEQ within-donor allelic test. Columns give the module’s age trait (linear or spline age model) and effect, the genes fitted, the genes with a transcript pair at Benjamini–Hochberg *q* < 0.05 and their fraction, the module’s age-association rank within its trait (0 weakest, 1 strongest), and the number of modules in that trait. Dorsolateral prefrontal cortex (dlpfc) ranks are missing because it has too few modules per trait to rank. Backs **Fig. S21e**.

| Region | Module | Age trait | Age effect | Genes fitted | Genes with pair at $q < 0.05$ | Fraction | Age rank within trait | Modules in trait (n) |
| --- | --- | --- | --- | --- | --- | --- | --- | --- |
| caudate | M000 | Age_linear | 0.2623 | 84 | 29 | 0.3452 | 1 | 3 |
| caudate | M002 | Age_linear | 0.1548 | 92 | 34 | 0.3696 | 0.5 | 3 |
| caudate | M004 | Age_linear | 0.1546 | 75 | 30 | 0.4 | 0 | 3 |
| caudate | M003 | Age_spline | 2.941 | 100 | 30 | 0.3 | 1 | 4 |
| caudate | M006 | Age_spline | 0.8375 | 80 | 40 | 0.5 | 0 | 4 |
| caudate | M008 | Age_spline | 1.672 | 38 | 17 | 0.4474 | 0.6667 | 4 |
| caudate | M011 | Age_spline | 1.422 | 12 | 5 | 0.4167 | 0.3333 | 4 |
| dlpfc | M000 | Age_spline | -2.046 | 58 | 20 | 0.3448 |  |  |
| dlpfc | M007 | Age_spline | -1.507 | 32 | 11 | 0.3438 |  |  |
| dlpfc | M008 | Age_linear | 0.2285 | 25 | 6 | 0.24 |  |  |
| hippocampus | M000 | Age_linear | 0.2664 | 28 | 12 | 0.4286 | 0.1538 | 14 |
| hippocampus | M001 | Age_linear | 0.4398 | 44 | 16 | 0.3636 | 0.9231 | 14 |
| hippocampus | M002 | Age_linear | 0.3454 | 28 | 6 | 0.2143 | 0.4615 | 14 |
| hippocampus | M003 | Age_linear | 0.4071 | 13 | 2 | 0.1538 | 0.7692 | 14 |
| hippocampus | M004 | Age_linear | 0.4236 | 77 | 22 | 0.2857 | 0.8462 | 14 |
| hippocampus | M006 | Age_linear | 0.1738 | 7 | 1 | 0.1429 | 0 | 14 |
| hippocampus | M007 | Age_linear | 0.3472 | 31 | 6 | 0.1935 | 0.5385 | 14 |
| hippocampus | M008 | Age_linear | 0.3936 | 22 | 9 | 0.4091 | 0.6923 | 14 |
| hippocampus | M009 | Age_linear | 0.3035 | 12 | 5 | 0.4167 | 0.3846 | 14 |
| hippocampus | M010 | Age_linear | 0.2809 | 3 | 3 | 1 | 0.2308 | 14 |
| hippocampus | M011 | Age_linear | 0.3874 | 8 | 6 | 0.75 | 0.6154 | 14 |
| hippocampus | M012 | Age_linear | 0.2845 | 4 | 0 | 0 | 0.3077 | 14 |
| hippocampus | M013 | Age_linear | 0.2474 | 1 | 0 | 0 | 0.0769 | 14 |
| hippocampus | M015 | Age_linear | 0.4505 | 1 | 1 | 1 | 1 | 14 |

**Table S9.**
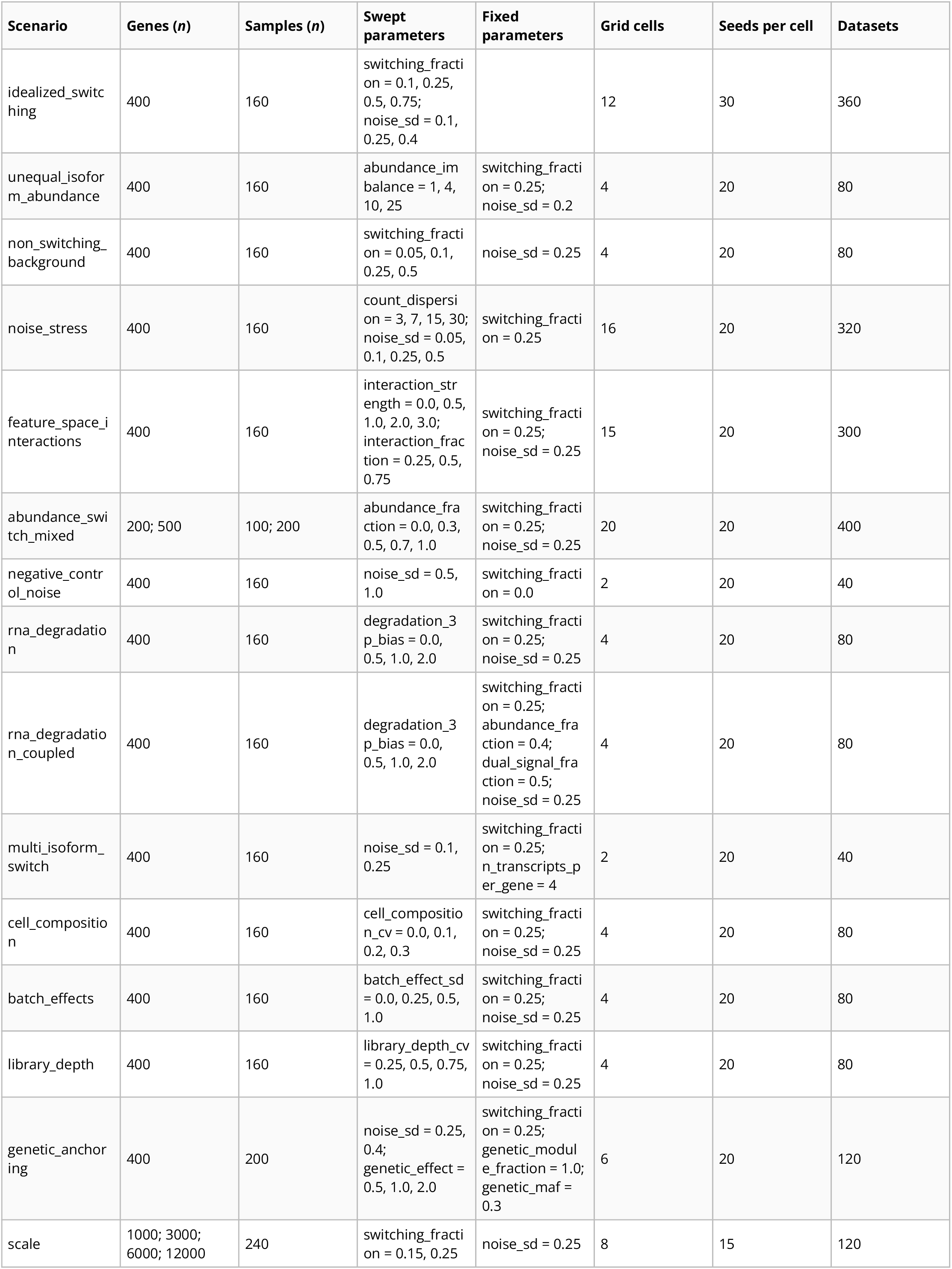
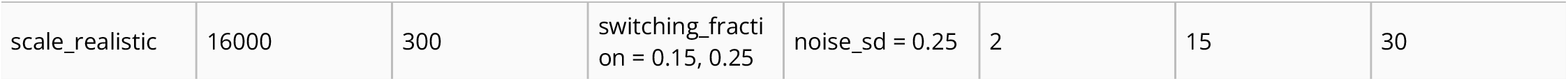
Synthetic benchmark scenarios. One row per scenario of the synthetic grid. Swept parameters were crossed within a scenario; the number of datasets is the number of grid cells times seeds per cell (30 for idealized switching, 15 for the scale scenarios, and 20 otherwise), and each method was run once per dataset. Scenario and parameter names are the identifiers used in the committed grid configuration, whose values are listed here.

### Supplementary Data

**Data S1. Discovery cohorts.** One row per analysis (17 analyses over 16 region-level cohorts; the BrainSEQ schizophrenia caudate analysis includes the 238 control donors of the aging caudate analysis). Each donor contributes one sample per analysis. Age is in years; Genotype-Tissue Expression (GTEx) exact age is released top-coded at 70. BrainSEQ reports postmortem interval (PMI, h) and GTEx tissue ischemic time (min). Genes and transcripts are counts after the bundle expression filter, before the prevalence filter applied at module fitting. The CSV adds library kit, self-reported ancestry, manner of death or Hardy scale, feature counts before filtering, and the single-nucleus RNA-sequencing (snRNA-seq) reference used for MuSiC deconvolution.

**Data S2. Switch-unique genes before and after adjustment for estimated cellular composition.** One row per analysis for the 12 analyses with an anatomically matched single-nucleus reference: genes tested, switch-unique genes before and after adding the estimated cell-type proportions to the conditional model, the number of unadjusted genes that remain `n_overlap`) and the number that adjustment newly introduces `n_new`), genes significant on both the switch and abundance coordinates in each arm, the number of cell types and samples covered by the deconvolution, and the number of module-by-cell-type marker tests with the counts reaching false discovery rate (FDR) < 0.05 for depletion and for enrichment. Because adjustment both removes and adds genes, the adjusted set is not nested in the unadjusted one, so `n_overlap` divided by the unadjusted count – not the ratio of the two counts – is the persistence statistic. The five Genotype-Tissue Expression (GTEx) regions with no matched reference are absent by design rather than entered as zeros. Backs **Fig. 2c,d** and **Fig. S14d**.

**Data S3. Per-region module trust funnel.** One row per cohort, region and method for the six anatomically paired cohort-region analyses. Columns cover split-half stability (module count, trusted count, and fraction above a size-matched permutation null at false discovery rate [FDR] < 0.05, with the median co-assignment density and its null), switch-driver reproducibility (median shared-gene driver-loading Spearman correlation and the fraction of positive pairs; IsoGraph only, as classical weighted gene co-expression network analysis [WGCNA] has no transcript-level drivers), the split-half aging counts (pairs tested, pairs significant for age in both halves, and pairs agreeing in sign), and complementarity to the abundance baseline. Best-match Jaccard is reported for description only: it is confounded by partition granularity, which is why the trust criterion is the chance-calibrated co-assignment test. Backs **Fig. 3a–d**.

**Data S4. Per-analysis module association rates for all four module sets.** The per-region ledger behind Table S4: one row per cohort, region, and method (68 rows) giving the module count, median module size, Gene Ontology enrichment count and fraction, and the counts and fractions of modules associated with the phenotype and with both coordinates. It is included because the pooled means hide the fact that the two cohorts order the methods differently, which is why the pooled ordering is not read as a method ranking. Backs **Fig. S15**.

**Data S5. Split-half module ledger.** One row per module per method (190 rows): module size, co-assignment density, the mean of its size-matched permutation null, permutation *P*, Benjamini– Hochberg *q*, the trusted flag and the best-match Jaccard against the most-overlapping half-fit module. Every module behind the 93 of 118 and 62 of 72 trusted counts is listed, so the counts can be audited module by module. Backs **Fig. 3b**.

**Data S6. Split-half agreement across the Leiden resolution sweep.** Mean adjusted Rand index (ARI) and normalized mutual information between the two halves of each donor split, over five seeds, at eight Leiden resolutions (0.5, 1, 2, 3, 5, 8, 12, and 20) for each cohort and region. The production resolution of 2.0 is flagged in `is_production`. The weighted gene co-expression network analysis (WGCNA) rows carry no resolution, because there is none to set, and are kept so the sweep is read against the fixed baseline rather than against itself. Averaged across regions, agreement varies little over the whole sweep (mean ARI 0.37–0.47, production 0.42), and the best resolution differs by region (0.5, 0.5, 1, 3, 20 and 20), so no resolution is uniformly best and the production value is not a tuned optimum. Per-region production values span ARI 0.19–0.57 for IsoGraph against 0.29–0.71 for abundance WGCNA on the same splits, the same granularity contrast the funnel reports throughout.

**Data S7. Cross-cohort eigengene projection ledger.** One row per trusted module per transfer direction (155 rows): the frozen-weight signed eigengene correlation against its feature-type-matched null with Benjamini–Hochberg *q*, the module’s age correlation in each cohort, that correlation expressed as a *z* statistic against same-weight random projections, and whether the two cohorts agree in sign. The final columns give the same sign agreement computed on the raw projected age correlations. These are the reason the reported statistic is standardized: among the modules significant in both cohorts on raw correlations, the three BrainSEQ-to-Genotype-Tissue Expression (GTEx) region pairs give sign agreement of 8 of 8 (caudate), 8 of 8 (dorsolateral prefrontal cortex to frontal cortex BA9) and 0 of 13 (hippocampus) – two regions unanimous in one direction and one unanimous in the other, which is a property of each target cohort’s own age-correlated structure rather than of the modules (**Data S27**). Backs **Fig. 3e**.

**Data S8. Cross-cohort matched-pair count against its permutation nulls.** The full grid of two methods by three statistics (covariate-free Pearson correlation, covariate-adjusted linear age term, covariate-adjusted natural cubic spline block *F* test) by three covariate modes by two nulls. The `age` null regenerates the target-cohort age association under the age-null while holding module construction and matching fixed; the `matching` null holds every age statistic fixed and permutes which target module each source module is matched to, and is the stricter of the two. The `is_published_statistic` flag marks the covariate-free Pearson arm quoted in the main text. Against the matching null that arm gives 1 of 38 (*P* = 0.91) for IsoGraph and 2 of 52 (*P* = 0.15) for weighted gene co-expression network analysis (WGCNA). The spline sensitivity arm with the matching null and `complement` covariate mode gives 0 of 38 (*P* = 1) and 5 of 52 (*P* = 0.16). The `complement` mode adds covariates outside the discovery residualization set; `full` adds the complete covariate set and `none` adds none. Spline significance uses a three-degree-of-freedom block *F* test, with direction taken from the corresponding linear age coefficient. Neither method exceeds the matching null in the reported Pearson or spline arm.

**Data S9. Functional preservation of matched cross-cohort module pairs.** Matched BrainSEQ– Genotype-Tissue Expression (GTEx) module pairs compared with a null that re-pairs each module with a random target module drawn from the same gene-count decile, because all three similarity measures increase with module size. Median gene Jaccard across matched pairs is 0.04, the gene-level overlap these measures are asked to look past. Rows with `n_finite` of zero are untested rather than negative: transcript-structure similarity had no computable pair in any region, and cell-type composition profiles are produced only for the IsoGraph arm. Both the linear and the spline age model are included; the main text reports the linear arm. Backs **Fig. 3f**.

**Data S10. satuRn concordance of IsoGraph switch genes, per analysis.** One row per cohort, region, and trait (17 analyses). Columns give the number of satuRn-tested genes, of switch genes (members of trait-associated IsoGraph switch modules), and of background genes, and the median satuRn differential transcript usage evidence (−log10 minimum isoform *P*) in each set. The primary test is the one-sided Mann–Whitney *P* with its rank-biserial effect size. The secondary is the odds ratio of switch-module membership per unit of evidence, adjusted for transcript count, with its *P*. Analyses with no switch genes are kept with empty statistics because nothing could be tested; they are not null results. Backs **Fig. S17**.

**Data S11. Gene-level corroboration of the switch layer by junction usage.** One row per analysis and switch set. Switch-positive genes are members of a trait-associated IsoGraph module (module set) or genes whose own switch coordinate is trait-associated (marginal set; BrainSEQ caudate only). Percent-spliced-in (PSI)-positive genes have at least one junction-level event (a PSI event in BrainSEQ, or a within-gene junction-usage feature in Genotype-Tissue Expression [GTEx]) associated with the trait under the same covariate model (within-gene Bonferroni, then Benjamini–Hochberg across genes, false discovery rate [FDR] ≤ 0.05). The adjusted odds ratio is from a logistic model of PSI status on switch status and log transcript count. The CSV adds the Fisher exact test, the PSI-positive rates in switch and background genes, and, for BrainSEQ, the junction-level resolution counts. Analyses with no switch-positive genes are listed with empty statistics.

**Data S12. Module context and held-out DTU evidence, per analysis.** One row per anatomically paired cohort-region analysis (10 split directions each). Columns give the tested genes and their coverage of the satuRn-tested universe, and the partial correlation of module context with held-out differential transcript usage (DTU) evidence (mean, median, range, and positive directions). Each correlation is given with its stratified permutation *P* and Benjamini–Hochberg *q*, before and after conditioning on the co-expression context, followed by the co-expression context’s own correlation and the resulting pattern. The sub-threshold columns give the fraction reaching held-out nominal Simes *P* < 0.05 among genes with discovery Benjamini–Hochberg *q* ≥ 0.05, in the top and bottom tertiles of module context, and the median adjusted odds ratio across split directions. These descriptive odds ratios adjust for discovery evidence and transcript features, without the additional co-expression predictor; nominal held-out support is not a false discovery rate (FDR)-significant replication call. The final columns are a post hoc check that recomputes the statistic without modules of 900 or more genes and without each split’s largest module. The decision rule recorded in the analysis code (*q* < 0.05 in at least four of six analyses, including one per cohort) applied to the model before co-expression adjustment and was met by four analyses. Three remained significant after co-expression adjustment; this additional model did not meet the four-of-six criterion. Backs **Fig. S18**.

**Data S13. RNA-binding-protein motif regulons of the switch modules.** One row per analysis, module, and RNA-binding protein (RBP; 11,040 cells). Columns give the module size, Gene Ontology (GO) visibility, and switch-gene counts; the hypergeometric test of RBP site-switch over-representation (expected count, enrichment, *P*, and Benjamini–Hochberg *q*); the opportunity-adjusted binomial model (coefficient, standard error, odds ratio with 95% confidence interval [CI], *P, q*, and fit status); and whether the cell is supported by both tests. Backs **Fig. S20a,b**.

**Data S14. ENCODE eCLIP binding at switched and constitutive exons.** One row per testable RNA-binding protein (RBP; 35). Columns give the nominated genes; the fraction with an enhanced crosslinking and immunoprecipitation (eCLIP) peak over switched and over constitutive exon intervals and their difference; the discordant gene counts, exact McNemar *P*, matched odds ratio with 95% confidence interval (CI), and false discovery rate (FDR); peak density per kilobase in each interval class with a gene-level bootstrap CI of their ratio; and whether binding is preferential and supported. Backs **Fig. S20c**.

**Data S15. Within-donor allelic test of isoform choice, per region.** One row per BrainSEQ region for the Benjamini–Hochberg-tested set of switch pairs. Columns give the pairs tested and fitted, genes fitted, pairs at nominal *P* < 0.05, and pairs and genes at Benjamini–Hochberg *q* < 0.05; the sign agreement of the within-donor effect with the between-donor dosage association (all fitted pairs and *q* < 0.05 pairs); fits that did not converge or reached the |β| = 10 bound; the homozygous-at-lead null (fits, fraction at *P* < 0.05, and the genomic inflation factor λGC) and λGC of the heterozygous test; the Spearman correlation of the mixed-model effect with the score-test *Z*; and the median lead-to-heterozygous-site distance. The module-axis columns test whether, within a gene, the effect tracks each pair’s module polarity, against a within-gene permutation null. The module-*cis* columns give the correlation of module *cis*-controlled-gene rate with age-association rank and its permutation *P*; the final row pools caudate and hippocampus. The last two columns count the separately fitted colocalization-nominated pairs, which carry a *q* only where they are also in the tested set. Backs **Fig. 5a** and **Fig. S21**.

**Data S16. sQTL colocalization nominations by signal-level and regional fallback support.** Every analysis-by-gene combination with a splicing quantitative trait locus (sQTL) posterior probability of a shared causal variant (PP4) ≥ 0.8 in the all-introns colocalization (127 rows). The aging and schizophrenia (SCZD) gene sets overlap, so the rows collapse to 119 gene-by-trait cells in 116 genes. Columns give the module and Gene Ontology (GO) visibility, the sQTL and expression QTL (eQTL) PP4, whether the call is sQTL-preferential (eQTL PP4 < 0.5), the tissues tested and colocalizing on each modality, and the headline tissue, intron, estimator `coloc.susie` or the `coloc.abf` fallback), and prior robustness. The headline PP4 is a maximum over correlated tissues and should be read with the tissue count. The sQTL-preferential flag is a selection on the sQTL call, not an estimate of splicing specificity; Data S17 is the unbiased comparison. Of the 119 cells, 33 were scored by `coloc.susie` in at least one tissue; the remaining 86 rest on the `coloc.abf` fallback.

**Data S17. Paired sQTL and eQTL colocalization contrast.** One row per analysis and a pooled row. Columns give the genes tested; the genes colocalizing with splicing quantitative trait loci (sQTLs), expression QTLs (eQTLs), both, only one modality, or neither (posterior probability of a shared causal variant [PP4] ≥ 0.8); the McNemar *P* of the discordant counts and the splicing share among them; and paired Wilcoxon comparisons of the conditional and raw PP4 of the two modalities.

**Data S18. BrainSEQ switch- and abundance-axis colocalization contrast.** One row per analysis and a pooled row, in European-ancestry BrainSEQ donors with in-sample linkage disequilibrium (LD). Columns give the genes testable on both IsoGraph axes; the genes colocalizing (posterior probability of a shared causal variant [PP4] ≥ 0.8) on the switch axis, the abundance axis, both, one axis only, or neither; the exact McNemar *P* and the switch share of the discordant genes; and paired Wilcoxon comparisons of the conditional and raw PP4. Backs **Fig. S22a**.

**Data S19. Colocalization events and their mapping to IsoGraph switch pairs.** One row per colocalization event, defined as a gene-by-trait nomination in one tissue with a splicing quantitative trait locus (sQTL) posterior probability of a shared causal variant (PP4) ≥ 0.8 (410 events over the 119 nominations of Data S16). Rows are sorted with the 115 events whose colocalizing junction is carried by an isoform of the tissue-matched switch pair first; these events cover 56 nominations, 21 of them signal-level. Columns give the analysis, trait, gene, and tissue; whether the nomination is signal-level (`coloc.susie` in at least one tissue); the estimator, sQTL and expression QTL (eQTL) PP4, prior robustness, and fallback reason of the event; the intron and junction; the junction-carrying transcripts and Gene Ontology (GO) visibility; whether the junction is carried by a switch-pair transcript, the switch pairs (up to 15 per gene) and the correlation of junction usage with switch polarity; the matched BrainSEQ region and whether the junction-carrying transcript also maps to a switch pair there; and whether the posterior-overlap layer (Data S21) also resolves the same gene-by-trait cell to a switch pair. `coloc.abf` rows name the representative intron of the Genotype-Tissue Expression (GTEx) release only. Risk-allele direction is resolved only in the posterior-overlap layer. Cross-cohort pair mapping does not establish replication of the phenotype association. Locus notes: *TMEM175* (amyotrophic lateral sclerosis [ALS], PP4 = 0.998) lies in a gene-dense region that also colocalized with Parkinson’s and Alzheimer’s disease at different lead variants, and few of its probes passed summary-data-based Mendelian randomization (SMR) without rejection by the heterogeneity in dependent instruments (HEIDI) test (Data S20); *SNCA* (dementia with Lewy bodies, PP4 = 0.970) maps to a switch pair in hypothalamus and cerebellum but not in cortex; *C9orf72* (ALS) colocalized on splicing in eight brain tissues (sQTL PP4 0.83–0.99, eQTL PP4 < 10-8) through `coloc.abf` only, maps to a cerebellar switch pair and also colocalized on the BrainSEQ switch axis (dorsolateral prefrontal cortex PP4 = 0.98; Data S18), and because the ALS association there tags the repeat-expansion haplotype it is read as a haplotype-linked splicing effect. Backs **Fig. 5b,c**.

**Data S20. SMR and HEIDI for the colocalization nominations.** One row per primary confirmatory probe, one per gene and quantitative trait locus (QTL) class (410 splicing QTL [sQTL] and 410 expression QTL [eQTL] probes). Columns give the analysis, trait, gene, QTL class, probe, and tissue; the top *cis*-QTL variant; the colocalization posterior probability of a shared causal variant (PP4) and estimator for the probe; the summary-data-based Mendelian randomization (SMR) estimate, standard error, and *P* with its Bonferroni threshold; the heterogeneity in dependent instruments (HEIDI) test *P* and number of variants; the instrument *F* statistic and weak-instrument flag; and the SMR status and its agreement with colocalization. “No instrument” means untested, not negative. Backs **Fig. S22b**.

**Data S21. Posterior-overlap-nominated isoform events and their mapping to IsoGraph switch pairs.** One row per overlap-supported quantitative trait locus (QTL) event (467 events), sorted with the 76 events whose junction maps into a tissue-matched switch pair first. Columns give the analysis, trait, and case; the gene, QTL type, and tissue; the lead variant, its risk allele, and the risk allele’s QTL effect; the junction and its credible-set posterior-overlap score (CLPP); Gene Ontology (GO) visibility; whether the junction is carried by a switch-pair transcript, the switch pairs (up to 15 per gene) and the junction-carrying transcripts; the correlation of junction usage with the switch polarity; and the matched BrainSEQ region with whether the junction-carrying transcript also maps to a switch pair there. This is cross-cohort pair mapping, not replication of the phenotype association. Corroborates the colocalization events of Data S19.

**Data S22. Partitioned heritability of the switch-derived QTL annotations.** Stratified linkage disequilibrium (LD) score regression of five genome-wide association study (GWAS) traits on variants that are Genotype-Tissue Expression (GTEx) brain quantitative trait loci (QTLs) of the aging switch genes (splicing QTL [sQTL], expression QTL [eQTL], or either) and, for schizophrenia, of the schizophrenia (SCZD) switch genes, conditional on baselineLD v2.2. Single-annotation models test each annotation alone; joint models include the sQTL and eQTL annotations together. Columns give the proportion of variants and of heritability, the enrichment and its *P*, and the per-variant coefficient with its standard error, *Z* and one-sided *P*. `coef_q_bh` is the Benjamini–Hochberg *q* across the 15 single-annotation tests of the aging switch genes; joint-model and SCZD-module rows are descriptive and have no *q*. Backs **Fig. 5e**.

**Data S23. Module concentration of posterior-overlap-supported genes.** One row per trait and aging cohort. Columns give the overlap-score-tested pool, the supported genes and the modules containing them, the observed concentration (sum of squared per-module counts) with its null mean and empirical *P*, and the fraction of overlap-supported genes in MAGMA-anchored modules (modules with a competitive gene-set *P* < 0.05) with the hypergeometric *P* under the overlap-score-tested pool and under all module genes. Dementia with Lewy bodies (LBD) in BrainSEQ had no overlap-supported genes. Backs **Fig. S24**.

**Data S24. MAGMA competitive gene-set tests of every module.** One row per module, trait and pipeline (8,064 rows): IsoGraph modules at Leiden resolution 2.0 (production) and 5.0 (sensitivity) and classical abundance-only weighted gene co-expression network analysis (WGCNA) modules, tested against eight genome-wide association studies (GWAS; schizophrenia, bipolar disorder, major depressive disorder, Alzheimer’s disease, Parkinson’s disease, dementia with Lewy bodies, amyotrophic lateral sclerosis, and stroke). Columns give the cohort, region and module identifier, module size, whether the module has at least 900 genes, the MAGMA coefficient, its standardized form and standard error, the *P* value, and the Benjamini–Hochberg false discovery rate (FDR) within trait and pipeline. Module identifiers are specific to the fit that produced them. Backs **Fig. S25**.

**Data S25. Long-read transcript detection and reciprocal usage of colocalization-anchored pairs.** One row per gene (55) whose colocalizing junction is carried by an isoform of a tissue-matched switch pair. Columns give the traits and the maximum splicing quantitative trait locus (sQTL) posterior probability of a shared causal variant (PP4); the distinct transcript pairs, those detected in the long-read data, and those that are switch-like, overall and restricted to pairs with an anchored isoform whose mean within-gene fraction is at least 0.05 (the well-expressed subset). Distinct unordered pairs are the primary unit. The table also gives counts over anchored–partner orientations, in which a pair anchored from both transcripts appears twice (sensitivity analysis); the strongest usage correlation; the anchored isoform’s maximum mean fraction; and whether the gene has at least one switch-like pair overall and at that abundance (the source “confirmed” fields). These calls do not test age association. Backs **Fig. 6a,b**.

**Data S26. Short-read junction corroboration of colocalization-prioritized transcript pairs.** One row per nominated transcript pair and primary BrainSEQ region (260 rows) for the 55 genes whose signal-level splicing quantitative trait locus (sQTL) colocalization junction is carried by an isoform of a tissue-matched IsoGraph switch pair. Columns give the gene, trait(s), Genotype-Tissue Expression (GTEx) tissue of the colocalization and its maximum sQTL posterior probability of a shared causal variant (PP4); the implicated junction and the two versioned transcripts of the pair; the BrainSEQ region and its anatomical match class (exact or generic-cortex proxy); whether the implicated junction itself discriminates the two alternatives and whether the recount model is two-sided; the number of nominating events; the coverage thresholds; the donors with completed recounts, the covered donors (at least 10 pair-discriminating fragments), the donors with at least one fragment for each alternative, and their fraction; the median total fragments and minor-form usage; the measurement status (measured, pair not counted, not two-sided, or insufficient coverage); and the descriptive donor-level Spearman concordance between junction-based and transcript-quantification usage, raw and covariate-adjusted, with its status. Counts pool haplotypes, including unassigned fragments, over adult control donors. Unmeasured rows are untestable rather than negative, and the concordance correlations share reads with the transcript estimates and carry no *P* values. Backs **Fig. 6c**.

**Data S27. Raw against null-standardized projected-age sign agreement, per region.** One row per method, transfer direction and region pair, with the switch-axis sign convention appended to the IsoGraph rows. The two arms are counted over different denominators and both are given: raw sign agreement over the modules whose projected age correlation is significant in both cohorts, and standardized sign agreement over every age-testable module. The raw arm is the reason the reported statistic is standardized – it is near-unanimous within a region and can be unanimous in either direction (8 of 8, 8 of 8, and 0 of 13 for the three BrainSEQ-to-Genotype-Tissue Expression [GTEx] pairs), which tracks the target cohort’s own age-correlated structure rather than the modules. The orientation columns give the second reason: the switch axis of a gene is fitted independently in each cohort, and about 45% of the genes that can be oriented (2,505–2,616 of 5,450–5,739 per region) carry the opposite sign in the two cohorts, so a raw projected coefficient inherits a sign the projection did not earn. These columns are empty for weighted gene co-expression network analysis (WGCNA), which projects abundance features and has no switch axis to orient.

**Data S28. Short-read junction-usage support for posterior-overlap-anchored pairs.** One row per candidate BrainSEQ percent-spliced-in (PSI) event (173 rows) for the 30 genes whose posterior-overlap-supported junction is carried by an isoform of the tissue-matched IsoGraph switch pair. Columns give the gene, trait, Genotype-Tissue Expression (GTEx) tissue of the colocalization, and the BrainSEQ region tested with its anatomical match class; the anchored junction, its credible-set posterior-overlap score (CLPP), and risk allele; the event and its type; the number of samples; the median PSI and minor-form usage (min(median PSI, 1 − median PSI)); the fraction of samples with minor-form usage of at least 0.05; and the verdict. A positive usage verdict denotes detectable alternative usage at the specified threshold, not phenotype replication. Events in GTEx tissues without a BrainSEQ counterpart are untestable rather than negative.

